# Experience shifts value updating from synaptic plasticity to neural dynamics

**DOI:** 10.64898/2025.11.30.691382

**Authors:** Jaeeon Lee, Jay A. Hennig, Vanessa Frelih, Samuel J. Gershman, Naoshige Uchida

**Affiliations:** Department of Molecular and Cellular Biology, Harvard University, Cambridge, MA, USA; Center for Brain Science, Harvard University, Cambridge, MA, USA; Department of Psychology, Harvard University, Cambridge, MA, USA; Department of Neuroscience, Baylor College of Medicine, Houston, TX, USA; Neuroengineering Initiative, Rice University, Houston, TX, USA

## Abstract

Adaptive behavior requires durable value memories and rapid updating. Experience in dynamic environments accelerates value updating, but the underlying neuronal mechanisms remain unclear. Mice exposed to repeated reversals of odor–outcome contingencies acquired faster value updating at the cost of faster memory decay. Recurrent neural networks (RNNs) trained with continual plasticity revealed two routes to faster learning, one through more efficient synaptic plasticity and one through a transition to value computation by neural dynamics. Only the latter reproduced the trade-off between updating speed and memory persistence observed in mice. Consistent with this transition, inhibiting dopamine neurons at reward delivery or pharmacologically inhibiting CaMKII in the amygdala impaired initial learning but largely spared expert reversal learning. In vivo recordings in the amygdala and hippocampus revealed representations of cue value and latent context necessary for rapid value updating. Context and value signals decayed over prolonged inter-trial intervals, paralleling behavioral forgetting. Conditional deletion of CaMKIIα in Emx1-cre-expressing neurons spared initial associative learning but impaired expert reversal performance, as did freezing the recurrent weights of RNNs, suggesting distinct loci for initial value learning and meta-learning. Together, these findings reveal how experience shifts value computation from synaptic plasticity toward neural dynamics, providing a mechanistic framework for the trade-off between memory stability and behavioral flexibility.

## Main

Animals must continually learn the value of sensory cues and actions to guide adaptive behavior. Reinforcement learning provides an algorithmic framework for this process^1–3^. Value memories should be durable when contingencies are stable, yet updated rapidly when contingencies change. How the brain implements value learning to meet these competing demands remains unclear.

Two mechanistic models of value updating have been proposed. In the ‘synaptic model’, the values of states or actions are stored in synaptic weights and modified incrementally during learning^4–11^. An influential version of this model posits that reward prediction errors encoded by midbrain dopamine neurons gate synaptic plasticity in downstream regions such as the striatum and the amygdala, converting state representations into value representations^4,5,12,13^. We use ‘synaptic model’ to refer to the broader family of models in which value updating is mediated by long-term synaptic plasticity. By contrast, in the ‘neural dynamics model’, value is updated through recurrent dynamics and maintained via persistent activity akin to working memory^2,14,15^. Synaptic plasticity would still be needed to learn the task structure, but once acquired, recurrent neural networks (RNNs) trained in this regime can update value purely through their activation dynamics^14,15^. Such networks integrate outcome information and maintain a persistent representation of context or recent reward history during the inter-trial interval (ITI), which is read out at the moment of choice to compute value.

Two observations are commonly cited in support of the neural dynamics model in decision-making tasks. First, animals can reverse choice within a few trials of a contingency change, a behavioral signature that has been argued to be too rapid for synaptic plasticity^15–18^. Second, single units in regions implicated in value computation exhibit persistent activity during the ITI that may convey a context or value signal used at the time of choice^16,18–21^. However, neither observation is uniquely diagnostic. Recently described forms of rapid synaptic plasticity, such as behavioral timescale plasticity (BTSP) in the hippocampus and neocortex^22–25^, could in principle drive value updates on the timescale of a few trials. Furthermore, persistent ITI activity reflecting context is equally consistent with the synaptic model if it reflects the readout of recently modified synaptic weights rather than a plasticity-independent integrator. Consistent with this, computational models^26,27^ suggest the two models can be indistinguishable at the behavioral level without direct causal manipulation of the underlying mechanism (e.g. blocking synaptic plasticity or persistent activity). Thus, computational accounts in which fast updating is carried purely by network dynamics, with the weights fixed after training^14,15,20^, rest on an untested assumption that synaptic plasticity no longer contributes once the task is learned. Overall, although the two models are not mutually exclusive, experimentally distinguishing their respective contributions to value computation remains a challenge.

Here, through a combination of modelling, causal manipulations and in vivo extracellular recordings, we dissociate the two mechanisms of value updating.

### Timescale of value updating and decay

We trained head-fixed mice on a classical conditioning task in which an odor cue (conditioned stimulus, CS+ or CS-) was followed by a water reward or no reward (Fig. 1a-c). In the stable task, the reward contingency was fixed, and mice had to initially learn a fixed value and maintain the value memory in the subsequent sessions. In the dynamic task, the contingency reversed in the middle of every session, and each session started with the contingency from the end of the previous one, so that mice had an incentive to remember previously learned values. Anticipatory licking during the odor (CS licks) served as a proxy for value. Mice discriminated the cues from day 1 of the stable task with small further improvement up to day 3. Mice took about 2 weeks to learn the dynamic task, after which they discriminated the cues as well as stable-task experts (Fig. 1d, f; EDF 1a-c).

**Fig 1.**
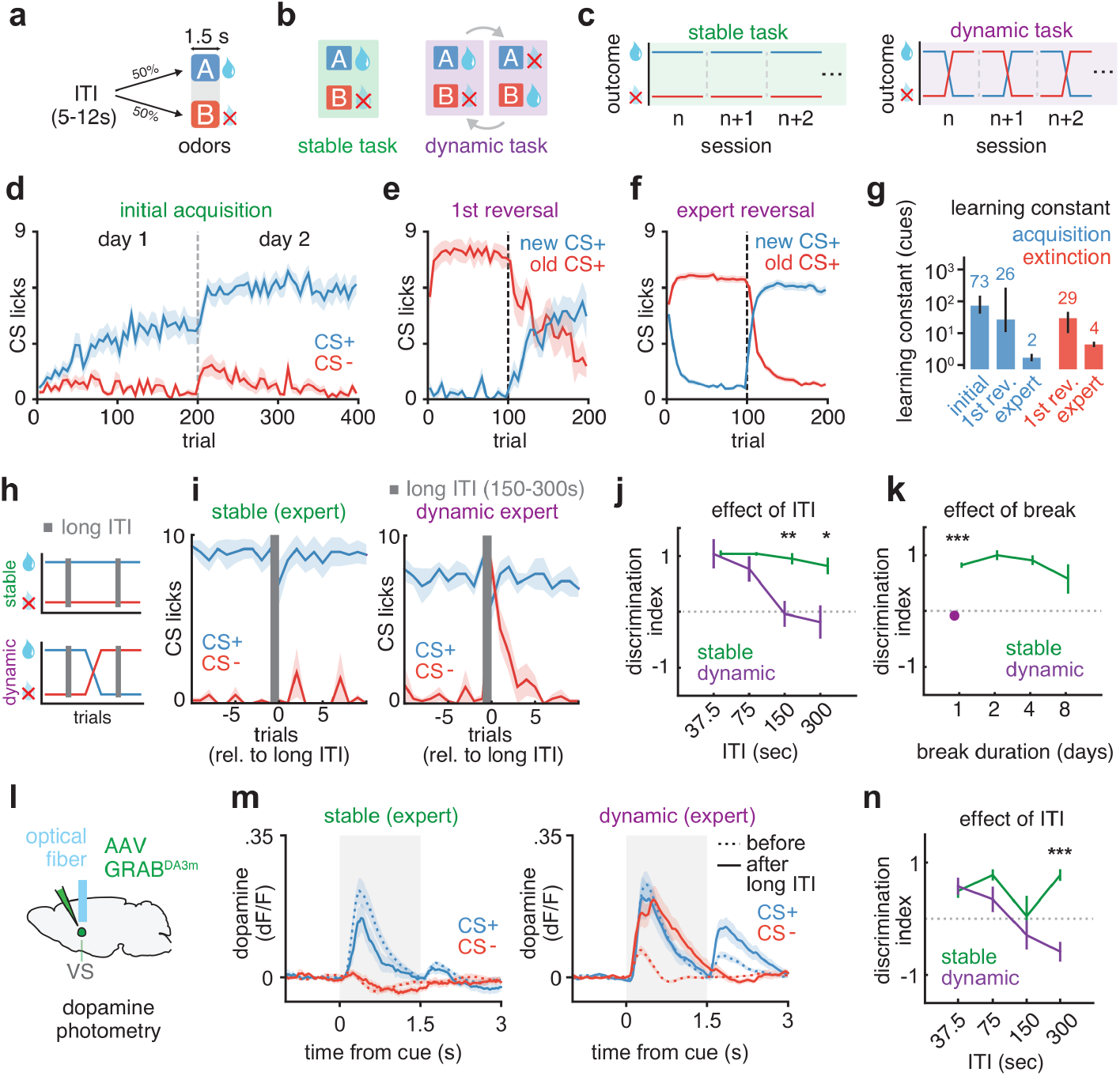
Environmental stability sets distinct timescales for value updating and forgetting. **a**, Trial structure (ITI, inter-trial interval). **b**, Task types (arrows, one reversal per session). **c**, Outcomes across sessions. **d**, Initial acquisition. CS licks (licks during the odor; 5-trial bins of session trials), stable task (n=16 mice). **e**, First reversal (n=8 mice). **f**, Expert reversals (n=188 sessions from 27 mice). Dashed lines, session boundaries or reversal. **g**, Learning constant in cue presentations (see Methods). Bars and numbers, medians; error bars, 95% bootstrap confidence interval (initial and 1st reversal, n as in d, e; expert, n=179 acquisition and 183 extinction sessions). **h**, Long-ITI test (see Methods). **i**, CS licks around a long ITI (trial, the n-th presentation of that cue; n=6 stable and 7 dynamic expert mice, one session each). **j**, Discrimination index, the difference in CS licks between the first CS+ and CS- after a long ITI divided by the difference over the last five presentations of each cue before it (1, retained; 0, lost; see Methods; per duration, n=6 stable mice, one long ITI each, and 7 dynamic mice, 7-8 long ITIs). **k**, Discrimination index after a break, normalized by the mean difference over that session (n=4 stable mice; n=15 dynamic sessions from 7 mice; ***, 1-day break, dynamic vs stable). **l**, Dopamine photometry in the ventral striatum (VS). **m**, Dopamine cue response, last five presentations before and first after a 150-300-s ITI (grey shading, odor; n=12 stable and 15 dynamic long ITIs from 6 and 7 mice). **n, Same as j, for the dopamine cue response** (mean dF/F, 0-1.5 s; index clipped to [−1, 1], see Methods; n=24 stable and 30 dynamic long ITIs from 6 and 7 mice). Dotted lines, 0. Two-tailed Student’s t-tests (j, k, n). *, *P*<0.05; **, *P*<0.01; ***, *P*<0.001. All other data shown are mean ± s.e.m.

We quantified the speed of learning by fitting an exponential to the CS licks as a function of the presentations of each cue (learning constant; see Methods). The learning constant fell from 73 presentations at initial acquisition to 26 at the first reversal and about 2 at expert reversals for the new CS+, and from 29 to 4 for the slower extinction of the old CS+ (Fig. 1d-g). Thus, as mice moved from a stable to a dynamic environment, value updating became faster by more than an order of magnitude, a speed-up of learning with experience that we refer to as meta-learning (i.e. learning-to-learn).

To understand how durable the value memory is in the two tasks, we inserted long inter- trial intervals (ITIs) during a session and quantified the CS lick response on the first CS+ and CS- presentations after each long interval (Fig. 1h; see Methods). Stable-task experts differentiated CS+ and CS- regardless of ITI duration (Fig. 1i, j). However, dynamic-task experts licked to the CS- on the first trials after the long ITI, with unchanged CS+ licking, before CS- licking subsided again (Fig. 1i), consistent with value memory being forgotten. The discrimination index, defined as the fraction of the cue discrimination retained across the ITI (see Methods), decreased with ITI duration (Fig. 1j). We also tested the effect of a long break (1 to 8 days) without any training (see Methods). On the first presentation of each cue after a break, discrimination remained high after up to 8 days in the stable task but was close to zero after 1 day in the dynamic task (Fig. 1k; EDF 1d). When dynamic-task experts were retrained on the stable task, discrimination on the first presentation of each cue in a session rose again, suggesting that this forgetting is reversible (EDF 1e).

As an alternative measure of value, we recorded the dopamine cue response in the ventral striatum (VS) (Fig. 1l; EDF 1i), which sped up with training as licking did (EDF 1f-h). After a long ITI, the CS- response of dynamic-task mice rose from near zero to a level comparable to the CS+ response, whereas in the stable task it stayed near zero (Fig. 1m; EDF 1j); the dopamine discrimination index, like the lick index, decreased with ITI duration in the dynamic task (Fig. 1n). Overall, as mice meta-learned, their value memory became labile. In both the behavioral and the dopamine readouts it decayed within minutes in the dynamic task, whereas in the stable task it survived long ITIs and, behaviorally, breaks of several days. This suggests that the two tasks engage distinct neural mechanisms of value computation.

### RNN model with continual plasticity

To understand the neural mechanism of meta-learning displayed by mice, we trained value RNNs to predict upcoming value during the same tasks, as in previous studies^15,27–29^. Most previous models update the weights only outside the task and then fix them after convergence, which assumes that plasticity plays no role in incremental improvement in performance. We therefore modified the learning algorithm to update the weights online, that is, continuously during the task rather than in a separate offline training phase, using truncated backpropagation through time (TBPTT) over a horizon of W timesteps (Fig. 2a; see Methods). Thus, the weights were never frozen, allowing a continual interaction between plasticity and dynamics. The window size, W, determines how much previous experience each learning update has access to. Unless stated otherwise we used a long window (W = 720 timesteps, about two blocks of the dynamic task), which we compare with a short one (W = 45, about six trials).

**Fig 2.**
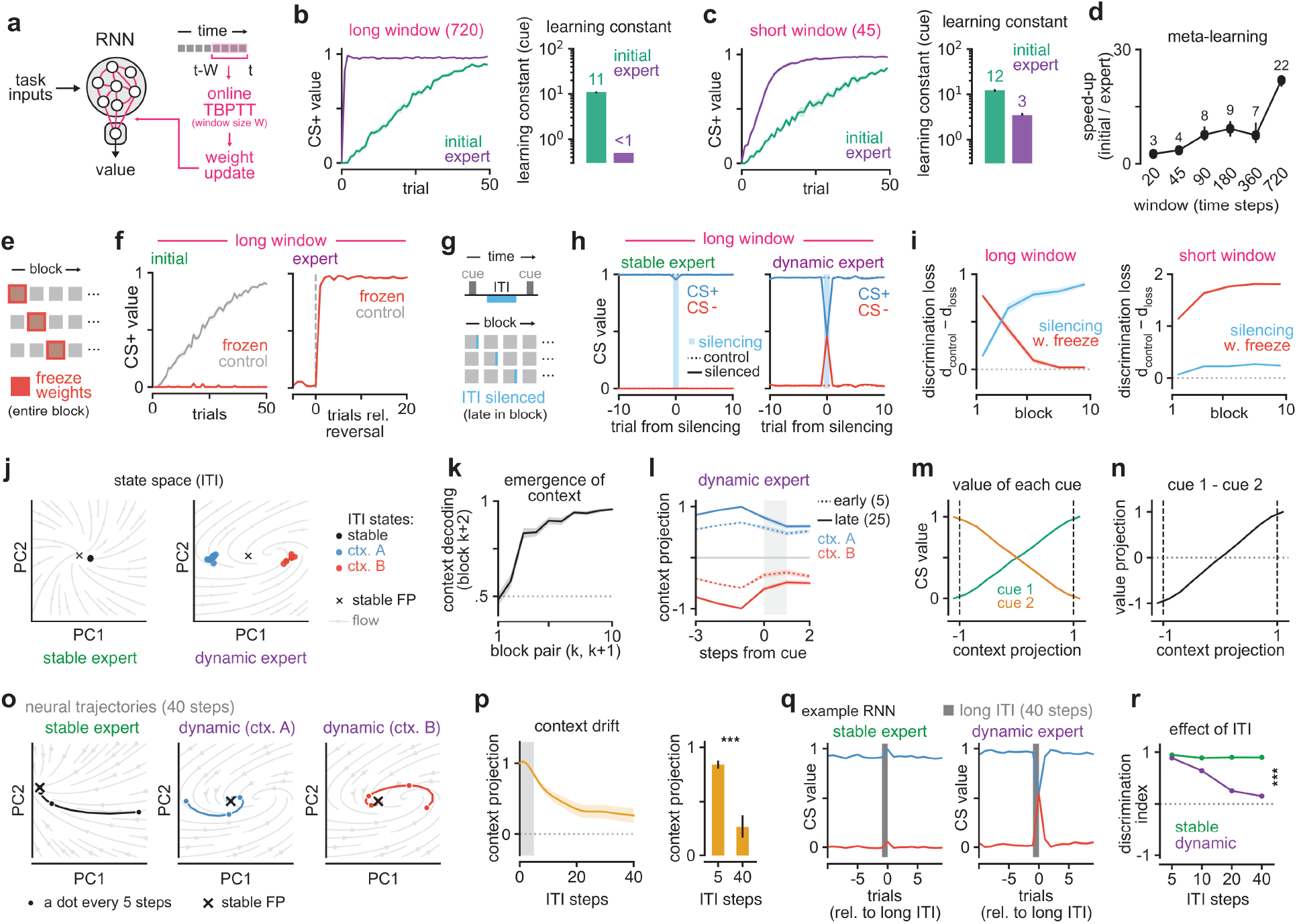
RNNs with continual plasticity recapitulate mouse behavior with shift from plasticity- to activity-based value updating. **a**, Recurrent neural network (RNN) updated online by truncated backpropagation through time (TBPTT) over the past W timesteps. **b, c**, CS+ value (trial, trial of the block) and median learning constant, in presentations of the cue; n=50 networks (initial and expert). **d**, Meta-learning speed-up (n=50 networks for W=20-720). **e, f**, Plasticity blocking (f, dashed line, reversal; n=50 networks; Extended Data Fig. 4m). **g, h**, ITI silencing: all units silenced for one ITI step (h, n=50 networks per set; Extended Data Fig. 4n). **i**, Discrimination loss (d=V(CS+) − V(CS-); n=50 networks per window and manipulation; Extended Data Fig. 3c). **j**, ITI state space of example experts (FP, fixed point; ctx. A/B, cue 1/2 rewarded). **k**, Context decoding of block k+2 (decoder trained on blocks k and k+1; n=100 networks). **l**, Context projection around the cue (grey band, cue and outcome; early/late, first 5/last 25 trials; n=100 networks). **m, n**, CS value as a function of context projection (dashed lines, context means; mean ± s.e.m. over trials; n=100 networks). **o**, Silent-ITI trajectories (PC plane of j). **p**, Context drift during silent ITIs (grey band, minimum training ITI; 1, context just experienced; 0, midway; n=50 networks; two-tailed Wilcoxon signed-rank test). **q, CS+ and CS- values of an example network around a long ITI (trial 0, first trial after it). r**, Discrimination index after a long ITI (see Methods; stable, n=50; dynamic, n=100 networks; two-tailed Mann-Whitney U test at 40 steps). Dotted lines, 0 or chance (k). Error bars in b-d, 95% bootstrap confidence interval. ***, *P*<0.001. All data shown are mean ± s.e.m. over networks unless stated otherwise.

RNNs learned to discriminate the CS+ from the CS- in either task regardless of window length (Fig. 2b, c; EDF 2). For the long window RNNs (long-RNNs), after repeated reversals the new CS+ value switched within a single presentation, close to expert mice (Fig. 2b; EDF 2d, e). For short window RNNs (short-RNNs), learning also sped up, with a smaller speed-up ratio (Fig. 2c; EDF 2d, e). Overall, even the shortest window size we tested resulted in meta-learning, with the learning speed-up generally increasing for longer window sizes (Fig. 2d).

To better understand the mechanism driving value updates and meta-learning in the RNNs, we either froze the weights for an entire block, or silenced ITI activity on single trials towards the end of each block at different points along training. This allowed us to test whether value updates were mediated by plasticity, and whether, at the end of the block, value computation depended on activity carried through the ITI. In long-RNNs, blocking plasticity (Fig. 2e; see Methods) abolished value acquisition in the initial block but left the expert reversal intact (Fig. 2f). Conversely, silencing all hidden units for one ITI timestep (ITI silencing; Fig. 2g) did not affect expert RNNs trained in the stable task, whereas cue discrimination collapsed on the silenced trial in expert RNNs trained in the dynamic task (Fig. 2h). Applying both manipulations in every block across learning revealed a transition from plasticity- to dynamics-based value updating (Fig. 2i; EDF 3, 4d; see Methods). Interestingly, this transition was not observed in the short window RNNs, with plasticity being necessary throughout training and ITI activity being mostly dispensable (Fig. 2i, right; EDF 3a, b). Intermediate windows fell in between (EDF 3c). The same online learning rule thus offered two routes to faster value updates, one that depends on plasticity (short window) and one that does not (long window).

What could be the mechanism driving value updating without plasticity for long-RNNs? Projected onto their first two principal components (Fig. 2j; see Methods), the ITI states of the stable expert sat next to a single stable fixed point, whereas those of the dynamic expert formed two clusters by context (which cue was rewarded). A fixed-point analysis using criteria similar to those of previous studies^30,31^ revealed that most RNNs had a single stable fixed point, whose position on the context axis varied across RNNs but peaked half-way between the two contexts. No RNN met our criteria for a line attractor, with a slow mode along the axis joining the two contexts (EDF 4a-c). A context code that generalized across blocks emerged with training (Fig. 2k). Projected onto this context axis, hidden-unit activity separated the two contexts, more so late in a block than in its first trials (Fig. 2l), consistent with the network updating its context estimate after each reversal. The position of the ITI state on this axis predicted the value of each cue, suggesting that the context held in the ITI state sets the value of the upcoming cue (Fig. 2m, n). Overall, these results suggest that in the dynamic task, RNNs gradually develop a representation of context in their persistent activity, with ITI states forming a cluster for each context. Because activity can move between these clusters without plasticity, the RNNs can rapidly update the value of each cue across blocks through neural dynamics alone.

Next, we simulated the effect of long ITIs as in the behavioral experiments (Fig. 1h-j). Given that context information encoded in the hidden units’ activity is important for computing value in the dynamic task, we reasoned that a long ITI might cause a drift in this activity, resulting in an ITI duration-dependent change in value discrimination. Consistent with this prediction, the ITI states of the dynamic expert drifted towards the fixed point between the two contexts (Fig. 2o). The context projection as well as upcoming CS value information decayed with the number of silent steps (Fig. 2p; example network in Fig. 2q), consistent with mice (Fig. 1j, n). The discrimination index fell with ITI duration only in dynamic-task RNNs (Fig. 2r), with most RNNs ending halfway between the two contingencies (EDF 4e). This effect was reproduced with other RNN variants (e.g. vanilla RNNs), although it was weaker when our Gated Recurrent Unit (GRU) networks were instead trained offline with full backpropagation through time, suggesting that online TBPTT yields dynamics that better match the forgetting of mice (EDF 4f, g). Overall, these results suggest that context information is held on a slow, leaky manifold, a region of state space along which population activity, without input, drifts slowly but steadily toward the fixed point, causing a time-dependent degradation of value memory in the dynamic task.

On the other hand, short-RNNs developed a context code, but their values depended less steeply on it, largely survived long ITIs and ITI silencing, and still required plasticity to reverse CS value (EDF 4h-n). Analysis revealed that short-RNNs meta-learned a strategy to make plasticity more efficient by increasing how much value is changed per unit of weight movement (EDF 5e) and reducing weight change not relevant for value reversal (EDF 5a–d, f–h). Both short- RNNs and long-RNNs relied on recurrent weights to store meta-learning (EDF 5i-l) but only the long-RNNs reproduced forgetting seen in mice (Fig. 2r; EDF 4l). Overall, these results suggest that fast value updates can be achieved via two distinct mechanisms (i.e. efficient plasticity vs neural dynamics). Interestingly, these two mechanisms predict qualitatively similar activity patterns (e.g. context coding), but might be dissociated causally by manipulating plasticity or activity during the ITI.

### Plasticity and persistent activity manipulations

RNN modelling revealed two distinct mechanisms of fast value updates, one requiring efficient plasticity and one relying on neural dynamics independent of plasticity. Thus, blocking synaptic plasticity in mice that are expert in the dynamic task could differentiate the two mechanisms. Global manipulation of plasticity in the whole brain can create aberrant conditions that might complicate the interpretation of the results. Thus, our approach was to first define a local plasticity manipulation that blocks initial value updates and see if the same manipulation affects later fast value updates. Phasic dopamine activity has been shown to be important for plasticity mediating value learning^5,13,32^. We thus hypothesized that inhibiting phasic activity around the outcome might block synaptic plasticity and value updates. We inhibited dopamine neurons in the ventral tegmental area at CS+ outcomes in mice expressing the light-gated anion channel gtACR1 in dopamine neurons (DAT-cre × gtACR1 mice), with light that abolished the dopamine response to unexpected reward in the VS (Fig. 3a-c; EDF 6a, b; see Methods). Mice were first habituated to the experimental setup for 3 days with unexpected water delivery, to minimize any learning unrelated to value learning (e.g. motor learning) on the subsequent test day. On stable day 1, brief inhibition of dopamine activity impaired value updates to the CS+ (CS+ licks), compared to controls, suggesting phasic dopamine activity is necessary for initial value updates (Fig. 3d, f, h). The same manipulation was applied after the reversal in mice that were expert in the dynamic task (Fig. 3b, c, e). Interestingly, although dopamine inhibition decreased the plateau level of CS+ licking, mice still updated value by increasing their CS+ licking above the pre-reversal level (Fig. 3e, g). The decrease in cue discrimination was stronger in the stable task than in the expert dynamic task (Fig. 3h), suggesting that although phasic dopamine activity is necessary for the initial value update, it becomes largely dispensable in the dynamic task. Similar results were obtained for CS dopamine activity (EDF 6d-f). Importantly, mice still consumed the reward on CS+ trials in both tasks (licks after the unconditioned stimulus, US; EDF 6c), suggesting that the impairment cannot be attributed to a general motor or motivational deficit.

**Fig 3.**
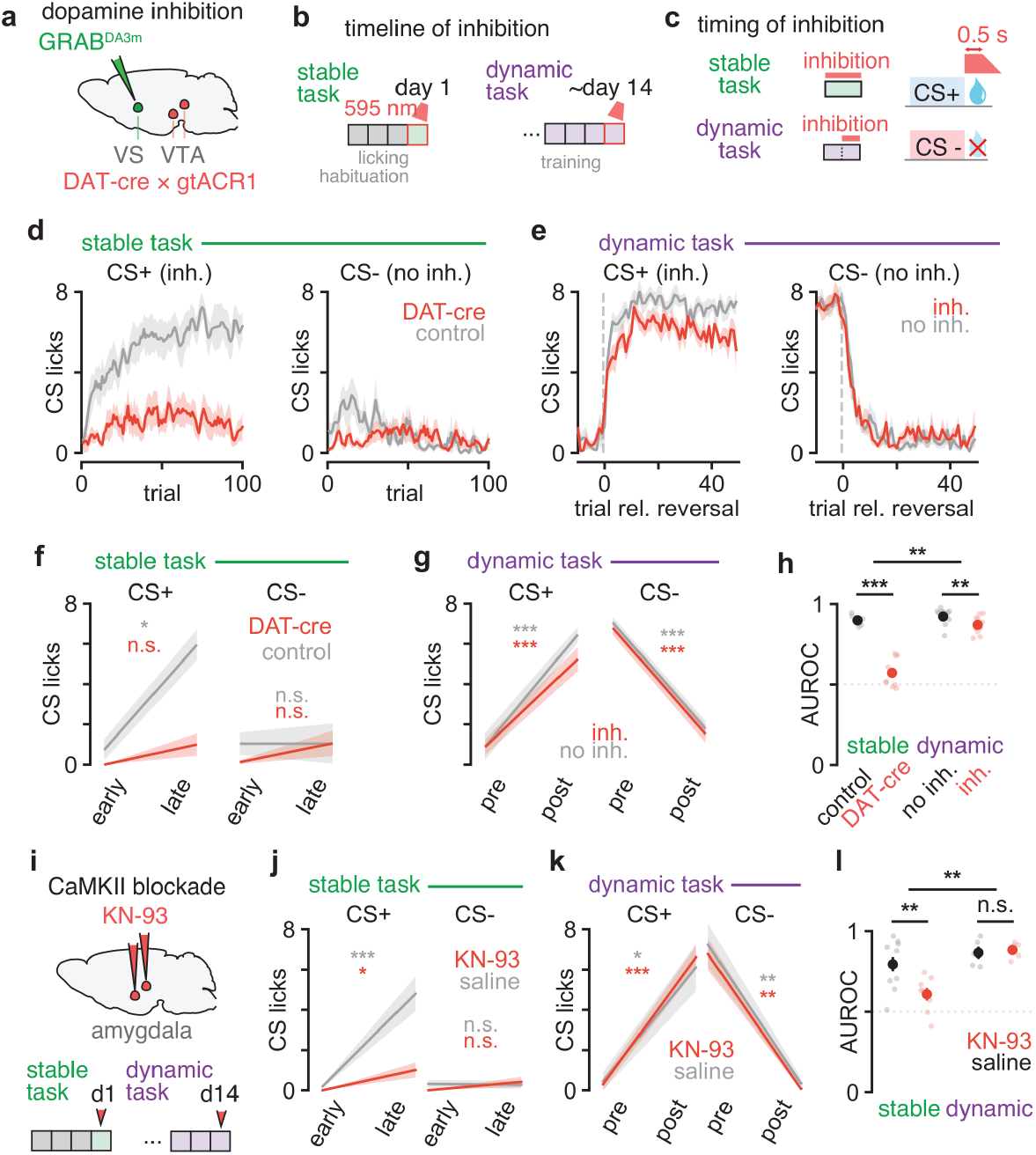
Two plasticity manipulations impair stable task learning but not value updating in the dynamic task. **a**, Inhibition of dopamine neurons in the ventral tegmental area (VTA); GRAB^DA3m^ recorded in the VS (see Methods). **b**, Timeline: stable day 1 or after day 14 dynamic-task session (red outline). **c**, Light at CS+ outcome on all trials (stable) or after reversal (dynamic). **d**, CS licks (licks during the 1.5-s cue; trial, the n-th presentation of that cue); control, mice without the opsin. **e, Same as d, in light (inh.)** and no-light (no inh.) sessions of the same mice; dashed line, reversal; CS+, the cue rewarded after it. **f**, CS licks early (session trials 1-5) and late (last 5). **g, Same as f, before (pre, 5 trials)** and after (post, trials 16-20) the reversal (see Methods). **h**, AUROC between CS+ and CS- licks over day 1 (stable) or 100 post-reversal trials (dynamic); pale dots, mice; dotted line, chance. **i**, Infusion of the CaMKII (Ca^2+^/calmodulin-dependent protein kinase II) inhibitor KN-93 or saline into the amygdala before stable day 1 (d1) or an expert dynamic session (d14). **j–l**, similar to f–h after saline or KN-93; k, post, last 5 trials of the post-reversal block. d–h: stable, n=7 control and 9 DAT-cre mice; dynamic, n=13 (no inh.) and 11 (inh.) mice, 31 and 26 sessions. j–l: stable, n=11 saline and 8 KN-93 mice; dynamic, n=5 mice (9 saline, 7 KN-93 sessions). Two-tailed tests: paired t-test (f, g, j, k, stars colored by group; dynamic h, l), Student’s t-test (stable h, l) and label-permutation test (top brackets, task difference of the effect). *, *P*<0.05; **, *P*<0.01; ***, *P*<0.001; n.s., *P*>0.05. All data shown are mean ± s.e.m. over mice.

Removing phasic activation of dopamine can block plasticity of D1-receptor-expressing striatal neurons^13^, but inhibition beyond baseline can also activate the protein kinase A pathway of D2-receptor-expressing striatal neurons, causing off-target plasticity^33,34^. Furthermore, dopamine inhibition could also change cell-intrinsic excitability as well as encoding properties^35–37^. Thus, to more cleanly manipulate plasticity while minimizing off-target effects, we pharmacologically inhibited plasticity that depends on calcium/calmodulin-dependent protein kinase II (CaMKII) in the amygdala, a brain region downstream of dopamine neurons that has been previously shown to undergo synaptic plasticity in both appetitive and aversive tasks^11,38–41^. We injected a CaMKII blocker (KN-93) into the amygdala before stable day 1 or an expert dynamic-task session (Fig. 3i; EDF 6g-i). KN-93 has been shown to effectively block synaptic plasticity in slices, and local infusion in vivo has been shown to cause behavioral effects consistent with an impairment of synaptic plasticity^13,42–45^. KN-93 infusion in the amygdala decreased the difference in anticipatory licking between CS+ and CS- trials in the stable task compared to saline infusion (Fig. 3j, l; EDF 6j, m–o). In stark contrast, the same manipulation had no effect in the dynamic task (Fig. 3k, l; EDF 6k). Mice still consumed the reward (US licks; EDF 6l), indicating that they remained engaged in the task. Overall, these results suggest that dopamine at the outcome and CaMKII in the amygdala are necessary to initially update value in the stable task, but become largely dispensable once mice are expert in the dynamic task, consistent with plasticity-independent fast value updates (model 1, EDF 7a).

One alternative explanation is that the amygdala becomes redundant in the dynamic task, with another region updating value through fast plasticity (model 2, EDF 7a). However, optogenetically silencing the basolateral amygdala (BLA) during the cue impaired discrimination in both tasks (EDF 7b-d; Emx1-cre × gtACR1 mice; see Methods), suggesting BLA activity is still necessary for the dynamic task, despite plasticity in the amygdala becoming dispensable. A third possibility is that the amygdala is still necessary in the dynamic task but inherits value or context information from upstream regions that compute these variables via fast plasticity (model 3, EDF 7a). To test this, we infused saline, muscimol or KN-93 into the orbitofrontal cortex (OFC) or the ventral hippocampus (vHip), two brain regions upstream of the amygdala and previously implicated in encoding context for reversal learning^18,46,47^ (EDF 7g, i; see Methods). Muscimol infusion in either the OFC or vHip impaired performance in the dynamic task, yet KN-93 infusion in either region did not affect performance (EDF 7h, j-l). These results suggest that CaMKII-dependent plasticity in either the OFC or vHip alone is not required for the fast value update.

The dynamics model further predicts that, in the dynamic task, cue value depends on activity carried through the ITI (Fig. 2g-i), so that disrupting it should impair the dynamic but not the stable task. We used Emx1-cre × gtACR1 mice to specifically silence excitatory neurons in either the basolateral amygdala (BLA), vHip, or anterior lateral motor cortex (ALM) during the ITI. ITI silencing in either the BLA or vHip impaired discrimination in the dynamic task but not in the stable task (EDF 7e, f). Interestingly, ITI silencing in the ALM impaired discrimination in both tasks (EDF 7f), consistent with its general role in motor preparation^48,49^. These results suggest that, in the dynamic task, activity in the BLA and vHip during the ITI is necessary for computing the value of the upcoming cue, consistent with context being held in persistent activity.

### Value and context coding in the amygdala and hippocampus

The RNN analysis suggested that context information is held and updated during the ITI, allowing value to be read out in the dynamic task (Fig. 2j-n). To test whether mice used a similar mechanism, we performed acute Neuropixels recordings in the amygdala and the hippocampus in mice performing the dynamic task with occasional long ITIs (Fig. 4a, b, g; EDF 8a-c; see Methods). Single units in both regions coded odor identity, outcome, value (responding to whichever odor was the CS+) or context (EDF 8d-i). Value units were the most common class in most subregions of both regions, and odor-identity units were more common in the amygdala (EDF 8f-h). Most amygdala and hippocampal value units fired more to the CS- than to the CS+ (EDF 8j), and context units made up about a fifth of the units in both regions (EDF 8i). Our recording sessions targeting the amygdala also allowed us to sample basal ganglia structures, including the posterior striatum and the external globus pallidus (GPe). Analysis of these regions revealed similar fractions of value and context units (EDF 8g, i), although, unlike in the amygdala and hippocampus, slightly more than half of the value units fired more to the CS+ than to the CS- (EDF 8j).

**Fig 4.**
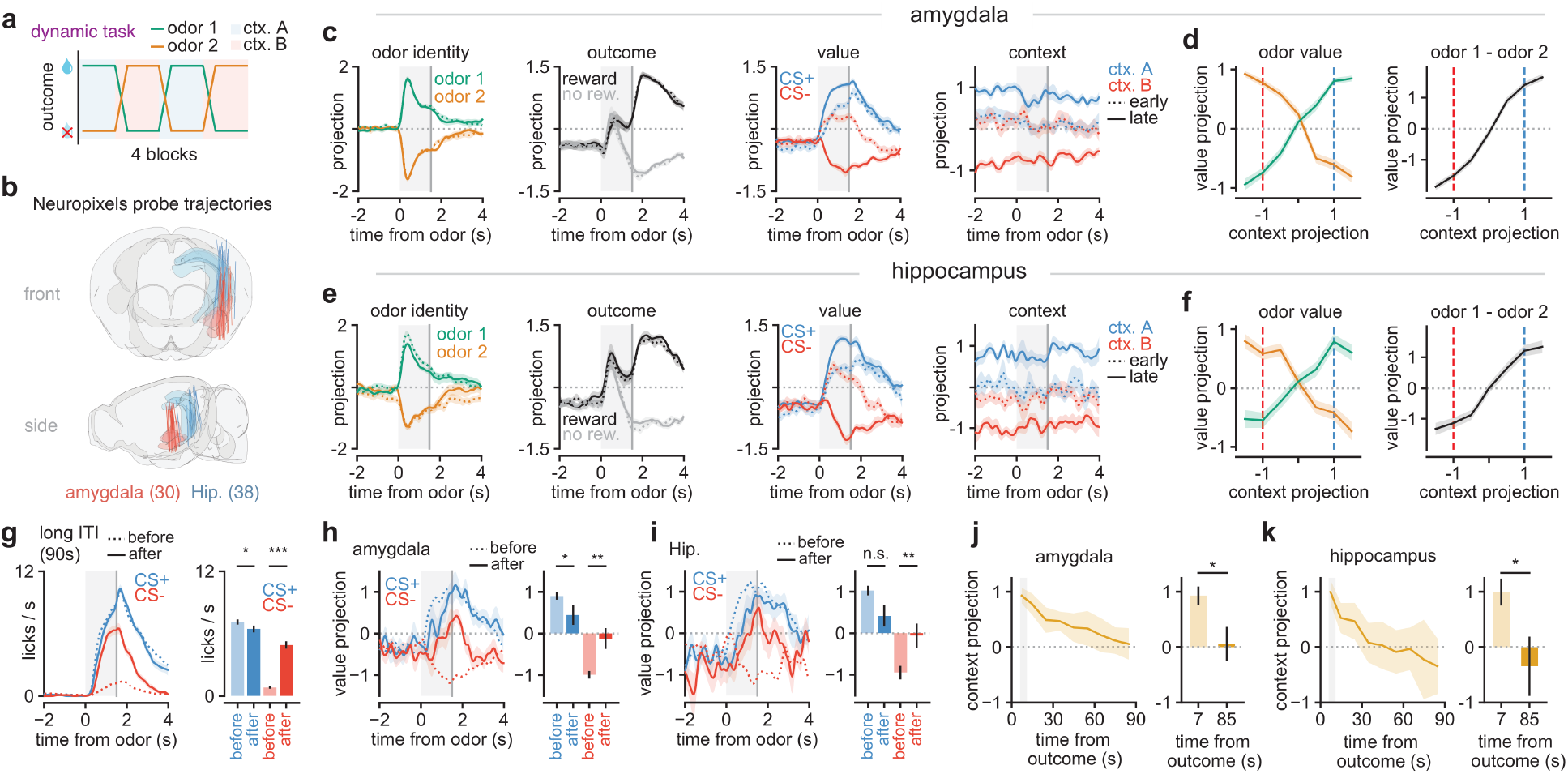
Value and context coding in the amygdala and hippocampus during the dynamic task. **a**, Recording task; ctx. A/B, blocks with odor 1/2 rewarded. **b**, Probe trajectories (amygdala, n=30 sessions; Hip., hippocampus, n=38 sessions; 7 mice; Extended Data Fig. 8a-c). **c**, Amygdala population projections (see Methods); solid/dotted are CS+/CS- (odor identity panel), the two odors (outcome panel) and late/early trials (value, context panels; last 10 of a block/first 10 after a reversal) (n=29 sessions from 7 mice; context, n=14 sessions from 5 mice). **d**, Value projection (0.5-1.5 s after odor onset) binned by context projection (2 s before odor onset); dashed lines, late-block levels of ctx. A (blue) and B (red) (n=3,176 trials, 14 sessions). **e, f, Same as c, d for the hippocampus** (e, n=33 sessions from 7 mice; context, n=11 sessions from 7 mice; f, n=2,458 trials, 11 sessions). **g,** Licking after the long ITI (see Methods); *left*, before (last 5 same-odor trials) and after (first trial); *right*, anticipatory lick rate (0.5-1.5 s after odor onset) (n=65/63 long ITIs for CS+/CS-; 64 sessions, 7 mice). **h, i, Same as g for the value projection** (as in d, f) (n=26/28 and 34/28 long ITIs; 27 and 31 sessions; 7 mice each). **j, k**, Context projection during the long ITI, normalized to the block’s level (1; see Methods); grey shading, normal ITI; *right*, normal-trial time (7 s) versus ITI end (85 s) (j, n=19 long ITIs, 12 sessions, 5 mice; k, n=16 long ITIs, 10 sessions, 7 mice). **c, e, g-i**, grey shading, odor; vertical line, outcome. Grey dotted lines, zero. **g-k**, two-tailed Wilcoxon signed-rank tests. *, *P*<0.05; **, *P*<0.01; ***, *P*<0.001; n.s., *P*>0.05. All data shown are mean ± s.e.m.

We then projected every trial onto odor-identity, outcome, value and context population axes, each fitted on simultaneously recorded units within a session (Fig. 4c, e; see Methods). In the amygdala, the value axis separated CS+ from CS- late in a block but only partially early after a reversal, when the projection of the former CS+ was still elevated. The context projection separated the block types throughout late trials, including before odor onset, but lay, on average, near the midpoint early after a reversal (Fig. 4c). The hippocampus showed the same organization with smaller odor-identity and outcome projections (Fig. 4e), as did the basal ganglia (EDF 8k).

Accordingly, CS+/CS- was decoded below chance on the first trial of a block in both regions, as was context in the amygdala, reflecting the previous contingency, whereas odor identity was decoded correctly; CS+/CS- and context decoding rose to a plateau within about 10 trials (EDF 8l, m). Importantly, the pre-odor context projection predicted the value the population assigned to the next odor. As it moved from one block type’s level to the other’s, the value projection of odor 1 rose and that of odor 2 fell, their difference crossing zero near the midpoint (Fig. 4d, f). This suggests that the context held before odor onset sets the value assigned to the upcoming odor, as in the RNNs (Fig. 2m, n).

The RNNs also predict that, without input, the context representation drifts toward the midpoint, and value discrimination decays (Fig. 2p-r), as lick discrimination does in dynamic-task mice after a long ITI (Fig. 1i, j). To test this, we analyzed the trials that followed a long ITI inserted at the end of a block (Fig. 4g; see Methods). After the long ITI, CS- licking increased and CS+ licking decreased only slightly (Fig. 4g), and the value projections changed in parallel (Fig. 4h, i). In long ITIs, the context projection decayed toward the midpoint in both regions, remaining near the block’s level early in the ITI and reaching the midpoint by its end (Fig. 4j, k). Overall, these results suggest that the amygdala and the hippocampus maintain a representation of the current contingency that covaries with the value of the next odor and relaxes over long ITIs in parallel with the animal’s value memory, as the long-RNN model predicts, consistent with the slow, leaky manifold of the RNN (Fig. 2o, p).

### Cortical and hippocampal CaMKIIα knockout impairs meta-learning

Acute blockade of plasticity in mice (Fig. 3) or in RNNs (Fig. 2e, f) largely spared the expert reversal. The RNN simulations predict, however, that plasticity is still needed during training to build the dynamics that carry value; blocking it throughout training should thus prevent the transition to dynamics-based updating but might spare the stable task. To test this, we generated conditional knockout mice lacking CaMKIIα in excitatory neurons of forebrain structures, including the cortex and hippocampus, thought to be important for meta-learning^14,15^ (Emx1-cre × CaMKIIα^fl/fl^; cKO; Fig. 5a; EDF 9a, b; see Methods). Previous studies have demonstrated the usefulness of this line for studying learning and memory^50^.

**Fig 5.**
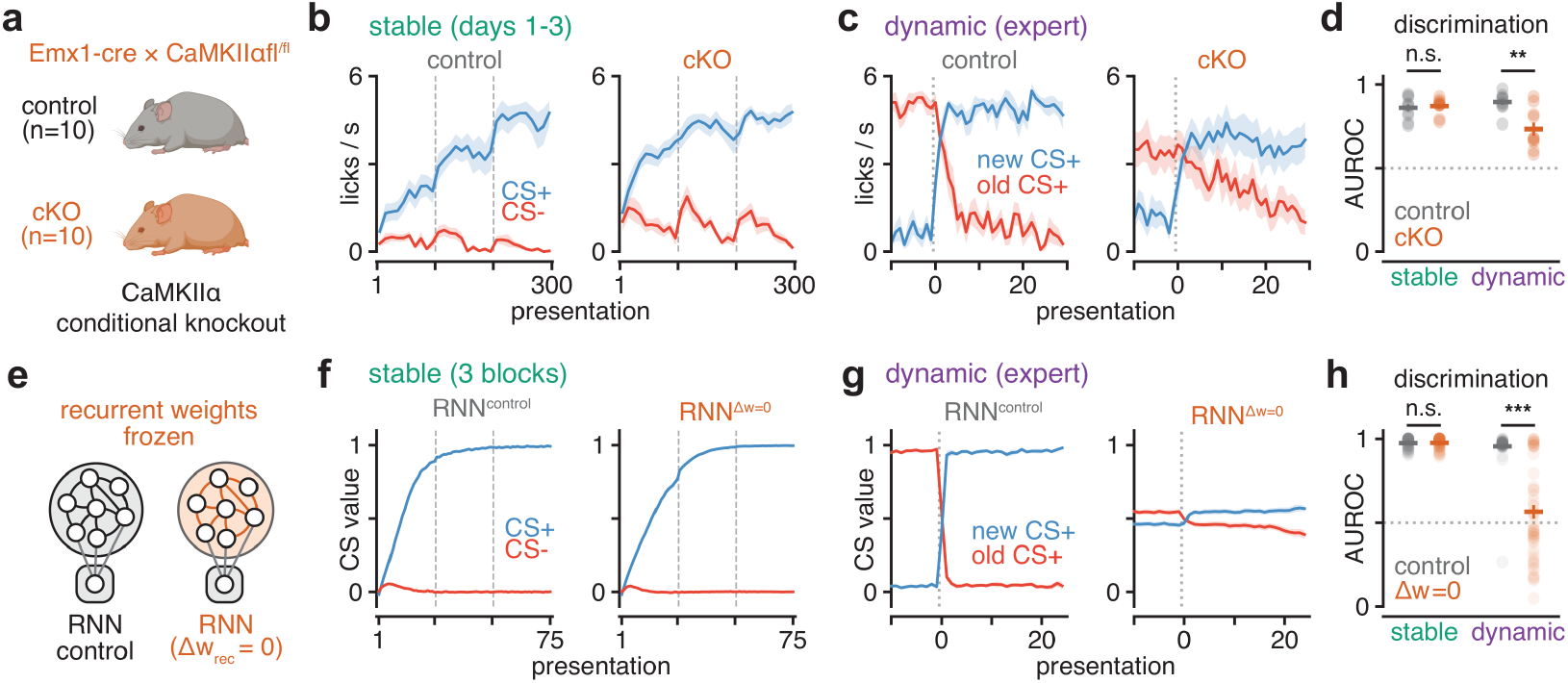
Conditional knockout of CaMKIIα impairs meta-learning. **a**, CaMKIIα conditional knockout. Emx1-cre × CaMKIIα^fl/fl^ mice (cKO) and Cre-negative littermate controls (see Methods). **b**, Stable task, days 1-3 (sta1 in Extended Data Fig. 9b). Lick rate (licks/s during the 1.5-s odor) for CS+ and CS- by presentation of each odor, the first 100 per session concatenated (dashed lines, session boundaries; see Methods). **c**, Dynamic task, expert session (last day of dyn2, Extended Data Fig. 9b). Lick rate for the new CS+ (rewarded after the reversal) and the old CS+ (rewarded before it) around the reversal, averaged over the session’s two reversals (see Methods). **d**, Discrimination. AUROC of the lick rate between CS+ and CS- trials of a session (stable, mean of days 1-3; dynamic, expert session; see Methods); dots, mice. **e**, RNN^control^, all weights plastic; RNN^Δw=0^, recurrent weights frozen at their initial values (see Methods; input or readout weights frozen in Extended Data Fig. 9i-m). **f**, Stable task, 3 blocks of 50 trials. Value at the cue for CS+ and CS- by presentation of each cue (25 per block, concatenated; dashed lines, block boundaries). **g**, Dynamic task, expert reversal (block 19, as in Fig. 2f). Value at the cue for the new and the old CS+ around the reversal (see Methods). **h**, Discrimination. AUROC of the value at the cue within a block (stable, mean of 3 blocks; dynamic, block 19), as in d; dots, networks. n=10 mice per group (a-d) and n=50 networks per group (f-h). Vertical dotted lines, reversal; horizontal dotted lines, chance. **d, h**, horizontal bar, mean; vertical line, s.e.m.; cKO or RNN^Δw=0^ versus control, two-tailed Mann-Whitney U test. **, *P*<0.01; ***, *P*<0.001; n.s., *P*>0.05. All data shown are mean ± s.e.m.

Over 3 days of the stable task, cKO mice learned as fast as their littermate controls (Fig. 5b, d; EDF 9c, f). cKO mice licked more overall (EDF 9e, g, h). In stark contrast, cKO mice were impaired at learning the dynamic task. After an expert reversal, control mice switched their licking within a few presentations, whereas in cKO mice licking to the old CS+ declined gradually and licking to the new CS+ rose less, lowering discrimination (Fig. 5c, d; EDF 9d). To model the knockout, we trained long-RNNs whose recurrent weights were frozen at their initial values from the start of training on either the stable or the dynamic task, while the input and readout weights remained plastic. Freezing the recurrent weights throughout training had the same effect in RNNs as in mice, with intact stable task learning but impaired dynamic task learning (RNN^Δw=0^; Fig. 5e-h; EDF 9i; see Methods). Freezing the input or readout weights instead left both tasks nearly intact (EDF 9j-m). In the RNNs, recurrent plasticity is thus needed to build the dynamics in order to compute value (Fig. 5g, h; EDF 5l, 9l, m). Overall, these results suggest that CaMKIIα-dependent plasticity in Emx1-cre-expressing neurons is dispensable for initial value learning, but necessary for meta-learning.

## Discussion

Through a combination of behavioral, circuit-level, and computational approaches, we show that value computation can transition from a synaptic plasticity-based to a neural dynamics-based mechanism. Although value updates using neural dynamics were fast, value memory decayed over intervals longer than the ITIs on which the animals were trained, suggesting a trade-off between stability and flexibility. Overall, our work provides a mechanistic framework to understand how the brain optimizes the trade-off between stability and flexibility for value computation.

Computationally, our results go beyond previous models, which relied solely on fast dynamics for value updating^14,20,26,27^. Here we show that a single online learning rule can produce two types of meta-learning, one through recurrent dynamics and one through more efficient weight updates. The latter route is consistent with previous experimental studies showing that prior learning, increases in excitability, or earlier induction of long-term potentiation can make subsequent plasticity more efficient^51–55^. Our online learning rule allowed us to study two distinct mechanisms of meta-learning without prior assumptions about how it occurs. Although we found that our work is consistent with a transition to dynamics-based value updates, it is possible that other task regimes might display meta-learning via more efficient plasticity. In our model, the route was set by how far back in time credit was assigned (the TBPTT window; Fig. 2i; EDF 3). Short windows favored more efficient plasticity and long windows neural dynamics, suggesting that tasks or circuits with shorter credit-assignment horizons might meta-learn through more efficient plasticity. Further work is needed to understand whether and when these meta-learning regimes might dominate. Although we used TBPTT to model the interaction of learning and dynamics, TBPTT is biologically implausible, and various alternatives that can mimic BPTT while preserving biological plausibility have been proposed^56–58^. The brain might approximate TBPTT using various cellular and circuit mechanisms, such as eligibility traces via synaptic tagging^8,59^, replay^60,61^ or dendritic plateau potentials^22,24^. Future work will be necessary to determine to what extent and how the brain approximates the TBPTT used here.

Although dopamine has been implicated in value learning, previous studies obtained mixed results on its role in fast value updates^62–64^. Our findings are consistent with the idea that dopamine becomes largely dispensable for driving value updates in dynamic tasks. This suggests that, contrary to previous models, the outcome signal that drives dynamics-based value updates comes largely from a source other than dopamine neurons. In our RNNs, this signal is an observation of the outcome itself, provided as an input, rather than a reward prediction error. Future work should identify where this outcome signal enters the regions that maintain value and context information. Our results are consistent with the finding that CaMKII-dependent plasticity in the OFC is needed for meta-learning but not for trial-by-trial value updating in experts^15^. In line with this, KN-93 infused into the OFC of expert mice had no effect (EDF 7h, k), whereas deleting CaMKIIα from Emx1-cre-expressing neurons selectively impaired meta-learning, suggesting a division of labor between the plasticity that supports initial learning and the plasticity that builds the dynamics for fast updating. Emx1-cre is also expressed in BLA excitatory neurons, yet performance in the stable task was spared in cKO mice but impaired by KN-93 infused into the amygdala. One explanation for the discrepancy is that infusion in the amygdala might also have targeted neighboring nuclei largely devoid of Emx1-cre-expressing neurons, such as the central or basomedial amygdala, which both our results (EDF 8f-h) and previous findings implicate in storing value^65,66^. Thus, blocking plasticity in the Emx1-positive cells in the BLA alone might not be sufficient to impair initial learning. A second explanation is that developmental knock out of CaMKIIα in BLA might be compensated by other regions taking over BLA’s role in initial value update. Future work should address this by targeting specific amygdala subregions, both acutely and genetically.

Although we found a persistent context representation in both the amygdala and the hippocampus, this does not by itself establish where context information is computed. ITI silencing of the BLA and vHip impaired performance in the dynamic task, but the effects were modest, suggesting that several brain regions, including regions not recorded here such as the OFC and the medial prefrontal cortex, might maintain context information together^16,67^. Comparing and contrasting the specific contributions of each brain region will be important for future studies.

Our work does not exclude the possibility that other forms of synaptic plasticity might contribute to value and context update using dynamics. For instance, recent findings and modelling suggest that short-term synaptic plasticity (STP) plays a role in maintaining information in the persistent activity^68–71^. However, STP decays within tens of milliseconds to seconds (facilitation, depression and augmentation) or, for post-tetanic potentiation, within minutes^72,73^. Thus, STP cannot support long-term memory storage and would be expected to degrade over the minutes-long intervals across which value memory decayed in our study. The exact contribution of other forms of plasticity that might contribute to the maintenance of information during the ITI needs to be investigated more carefully in the future.

Our work has parallels with previous modelling approaches contrasting weight-based (in-weight learning) versus activity-based approaches (in-context learning)^74–80^. Computationally, meta-learning can be achieved by optimizing gradient descent similar to our fast-plasticity approach^81^ or through context held in the network’s activity (in-context learning)^74,82^. Overall, both biological and artificial systems might optimize the learning algorithm by optimizing the weight update process or using a weight-independent process. A deeper understanding of these trade-offs might help us gain insight into intelligent and adaptive learning algorithms.

Finally, our results reveal a fundamental trade-off between stability and flexibility in value computation. Dynamics-based value representations enable rapid value updates at the expense of degradation over time, whereas plasticity-based representations provide stable long-term storage at the cost of being slow and inflexible. Given the further cost of having to maintain information in persistent activity, dynamics-based value updates might only emerge when the benefits of faster value updates (e.g. when the environment is dynamic) are high. Our work suggests that the mode of value computation depends on the stability of the environment.

## Methods

### Experimental procedures

#### Animals

A total of 75 male and female wild-type C57BL/6J mice (The Jackson Laboratory) were used for the behavioral and fiber-photometry experiments (Fig. 1, Extended Data Fig. 1), the KN-93 infusion experiments in the amygdala (stable task: n = 11 saline and n = 8 KN-93 mice; dynamic task: n = 5 mice; Fig. 3i-l, Extended Data Fig. 6g-o), the muscimol and KN-93 infusion experiments in the orbitofrontal cortex (OFC) and ventral hippocampus (vHip) (n = 8 mice; Extended Data Fig. 7g-l) and the Neuropixels recordings (n = 7 mice; Fig. 4, Extended Data Fig. 8). Additional genetically modified mice were used as follows. For optogenetic inhibition of dopamine neurons, DAT-IRES-Cre mice (JAX stock no. 006660) were crossed with mice carrying a Cre-dependent GtACR1 reporter allele (R26-LNL-GtACR1-Fred-Kv2.1; JAX stock no. 033089); experimental mice were heterozygous for both alleles (n = 13 mice, of which 9 were tested in the stable task and 13 in the dynamic task; Fig. 3a-h, Extended Data Fig. 6a-f). Control mice for the dopamine inhibition experiments did not express the opsin but received the same implants and illumination (n = 7 mice; heterozygous for the GtACR1 reporter allele and negative for DAT-IRES-Cre). For optogenetic inhibition of Emx1-cre-expressing excitatory neurons, Emx1-IRES-Cre mice (JAX stock no. 005628) were crossed with the same GtACR1 reporter line; experimental mice were heterozygous for both alleles (17 mice in total: n = 6 mice for cue and ITI silencing of the BLA (5 of them for cue silencing), n = 7 mice for ITI silencing of the vHip and n = 4 mice for ITI silencing of the anterior lateral motor cortex (ALM); Extended Data Fig. 7b-f). To conditionally delete CaMKIIα in Emx1-cre-expressing cells, Emx1-IRES-Cre mice were crossed with mice carrying a floxed Camk2a allele (JAX stock no. 006575). Conditional knockout mice were homozygous for the floxed Camk2a allele and heterozygous for the Emx1-IRES-Cre allele (Camk2a fl/fl; Emx1-Cre/+; n = 10 mice). Cre-negative littermates homozygous for the floxed Camk2a allele (Camk2a fl/fl; Emx1+/+; n = 10 mice) served as controls. The knockout and control mice were run in two cohorts of 5 mice per group (Fig. 5a-d, Extended Data Fig. 9a-h).

Animals were housed on a 12-h dark–12-h light cycle and performed the task at the same time each day (±1 h), during the dark period. Ambient temperature was kept at 75 ± 5 °F, and humidity was kept below 50%. Animals were group-housed (2–5 animals per cage) until surgery, then individually housed throughout the experiment. All procedures were performed in accordance with the US National Institutes of Health Guide for the Care and Use of Laboratory Animals and approved by the Harvard Institutional Animal Care and Use Committee.

#### Surgeries

All surgical procedures were conducted under aseptic conditions. Mice (older than 8 weeks) were anesthetized with isoflurane (3.5% for induction, followed by 1–2% for maintenance at 1 L min⁻¹). A local anesthetic (2% lidocaine) was administered subcutaneously at the incision site. Analgesia was provided with buprenorphine (0.1 mg kg⁻¹, i.p.) pre-operatively and ketoprofen (5 mg kg⁻¹, i.p.) for two days post-operatively. After leveling, cleaning, and drying the skull, a custom-made titanium head plate was attached using adhesive cement (C&B Metabond, Parkell). For all injections, the solution (virus, KN-93 or muscimol) was loaded into a pulled glass pipette (5-000-1001-X, Drummond) backfilled with mineral oil and fitted with a plunger. A small craniotomy (<1 mm diameter) was made using a dental drill, and the pipette assembly was mounted on a stereotaxic holder, lowered to the target coordinates, and injected slowly (∼100 nL min⁻¹) to minimize tissue damage (MO-10, Narishige). After each injection, the pipette was left in place for at least 5 min to allow diffusion before being raised to the next site or withdrawn from the brain. Target coordinates for the brain regions were: ventral striatum (VS): 1.0/1.5/3.8 mm, amygdala: −1.2/3.3/4.2 mm, OFC: 2.6/1.9/1.8 mm, vHip: −3.4/3.2/4.2 mm (anterior-posterior/medial-lateral/dorsal-ventral; coordinates are relative to bregma, and DV relative to the surface of the brain). For fiber photometry, an optical fiber (200 μm, 0.66 NA, Doric) was implanted above the VS injection site in the left hemisphere. For optogenetic inhibition of dopamine neurons, optical fibers (400 μm, 0.66 NA, Doric) were implanted bilaterally above the ventral tegmental area (VTA) at AP −2.9 mm and ML ±1.75 mm, with an insertion depth of 4.0 mm from the brain surface along a trajectory angled 15° from vertical toward the midline (Extended Data Fig. 6a); in these mice a second fiber (200 μm, 0.66 NA, Doric) was implanted above the VS for photometry (Extended Data Fig. 6a). For optogenetic inhibition of Emx1-cre-expressing neurons, optical fibers (400 μm, 0.66 NA, Doric) were implanted bilaterally above the BLA (−1.2/3.3/4.0 mm), the vHip (−3.4/3.2/3.3 mm) or the ALM (2.5/1.27 mm, just above the brain surface). For the KN-93 infusions in the amygdala, the craniotomies were made the day before the first infusion and the drug was injected through a glass pipette on the day of testing (see KN-93 infusion into the amygdala). For the muscimol and KN-93 infusions in the OFC and vHip, craniotomies above both regions were made during the head-plate surgery and sealed with silicone elastomer (Kwik-Cast) until the infusion days, on which the drug was injected through a glass pipette (see Muscimol and KN-93 infusions into the OFC and ventral hippocampus). For the Neuropixels recordings, craniotomies were made above the amygdala and above the hippocampus the day before the first recording session (at the amygdala and vHip coordinates given above; the registered probe tracks were centred at AP −1.6 mm and AP −3.4 mm, respectively; Extended Data Fig. 8a, b) and sealed with silicone elastomer until the recording day (see Neuropixels recordings). Post surgery, mice were allowed to recover for at least 2 weeks before beginning the experiment.

#### Viruses

To record dopamine using fiber photometry, we expressed AAVs encoding the green dopamine sensor GRAB-DA3m (AAV-hsyn-DA3m(h-D05), WZ Biosciences, 300 nL, concentration 5×10¹² gc/mL) in the left hemisphere VS of wild-type mice (Fig. 1l, Extended Data Fig. 1i) and of the DAT-cre × gtACR1 and control mice used for the calibration of dopamine inhibition and for the cue dopamine responses (Extended Data Fig. 6a, b, d-f).

#### Histology

Mice were deeply anesthetized with an overdose of ketamine/medetomidine, exsanguinated with 0.9% phosphate-buffered saline (PBS) and transcardially perfused with cold 4% paraformaldehyde (PFA) in PBS. The brain was extracted from the skull and stored in 4% PFA for 24–48 h at 4 °C, after which it was rinsed and stored in PBS and cut into 100-μm sections on a vibratome (VT1000S, Leica). Sections were mounted on slides and imaged with a slide scanner (Zeiss Axioscan 7).

#### Behavioral setup

All behavioral experiments took place inside a custom-built enclosed behavioral box in which head-fixed mice could stand on a fixed running wheel. The frame of the behavioral box was built using aluminum frames (McMaster) and the walls were made using black hardboard (Thorlabs, TB4). Mice had access to a water spout which delivered artificially sweetened water (Acesulfame Potassium powder dissolved in water at 6 g/L; Prescribed For Life) for water reward, and an odor spout which delivered the odor cues for classical conditioning. Behavioral events were controlled (and licking was monitored) using custom-written software in MATLAB (Mathworks) and the Bpod library (Sanworks) interfacing with the Bpod state machine (1024 and 1027, Sanworks), valve module (1015, Sanworks) and port interface board (1020, Sanworks)/water valve (LHDA1233115H, Lee Company) assembly. Odors were delivered using a custom olfactometer, which directed air through one of eight solenoid valves (LHDA1221111H, Lee Company) mounted on a manifold (LFMX0510528B, Lee Company). Each odor was dissolved in mineral oil at 10% dilution, and 30 μL of diluted odor solution was applied to a syringe filter (2.7 μm pore, 13 mm diameter; 6823-1327, Whatman). Wall air was passed through a hydrocarbon filter (HT200-4, Agilent Technologies) and split into a 100 mL min⁻¹ odor stream and 900 mL min⁻¹ carrier stream using analogue flowmeters (MFLX32460-40 and MFLX32460-42, Cole-Parmer), which were recombined at the odor manifold before being delivered to the nose of the mouse. During photometry experiments, licking was monitored using a closed-loop circuit similar to a previously described method^83^. During in vivo electrophysiology, licking was monitored with an infrared emitter–photodiode pair, which is less prone to electrical artifacts and thus more suitable for in vivo electrophysiology.

#### Behavioral tasks

We used a classical conditioning paradigm in which odor cues predict specific outcomes (Fig. 1a). An odor was chosen pseudo-randomly (out of two odors, 50% each) and was delivered for 1.5 s, after which an outcome (1.5 μL of artificially sweetened water or no reward) was delivered via the water spout. An inter-trial interval (ITI) separated each trial. The duration of the ITI, measured from the outcome to the next cue, followed a truncated exponential distribution with a mean of 8 s, a minimum duration of 5 s and a maximum duration of 12 s. Each session consisted of 200 trials, which lasted about 35 min on average. In the stable task, odor A ((S)-(−)- limonene) was always rewarded and odor B (1-heptanol) was never rewarded (Fig. 1b, c). In the dynamic task, the reward contingency changed once every session, with the reversal happening at the 101st trial (the middle of the session), and every session started with the contingency that the mouse had last seen at the end of the previous session, to preserve the continuity of the reward contingency across days (Fig. 1b, c). The CaMKIIα conditional knockout mice and their controls were trained on two versions of the dynamic task (Extended Data Fig. 9b): a task with one reversal per session (dyn1; 2 blocks of 100 trials) and a task with two reversals per session (dyn2; 3 blocks of 70 trials, 210 trials per session). For the Neuropixels recordings we used a version of the dynamic task with four blocks per session, the reward contingency of the two odors reversing at every block transition (Fig. 4a). Two protocols were used: 240 trials (4 blocks of 60 trials) and 300 trials (4 blocks of 50 trials with the two familiar odors, followed by 2 blocks of 50 trials with two novel odors). In these sessions the ITI (from the outcome to the next cue) followed an exponential distribution truncated between 5 and 11 s, and in 64 of the 68 recording sessions the last trial of blocks 2 and 3 was preceded by a long ITI of 90 s from the previous outcome to the cue (the 2.5- s post-outcome period followed by an 87.5-s ITI state; see Task during the recordings) (Fig. 4g-k). To test the effect of long ITIs on value memory (Fig. 1h-j, m, n; Extended Data Fig. 1j), two long ITIs of 37.5, 75, 150 or 300 s (measured from the previous outcome to the next cue) replaced the normal ITIs preceding trials 50 and 150 of an otherwise normal 200-trial session of the stable or dynamic task; each session contained either the 75-s and 150-s or the 37.5-s and 300-s long ITIs, in either order, and the first cue after the long ITI was the CS- (52 of the 54 long ITIs). To test the effect of breaks between sessions (Fig. 1k), normal sessions were separated by breaks of 1, 2, 4 or 8 days in the stable task and of 1 day in the dynamic task. Extended Data Fig. 1d shows the last 10 trials of a session and the first 10 trials of the next session across the regular 1-day break between daily sessions (stable task, n = 6 mice; dynamic task, n = 5 mice; one session pair per mouse). To test whether forgetting after a break was reversible (Extended Data Fig. 1e), one cohort of mice (n = 6) was trained in sequence on the stable task (12 sessions), the dynamic task (12 sessions) and again the stable task with the contingency of their last stable sessions (retraining; 6 sessions).

#### Odors

We used (S)-(−)-limonene (odor A) and 1-heptanol (odor B) for the stable and dynamic tasks. All odors were diluted in mineral oil at 10%.

#### Behavioral training

Mice were first handled while undergoing water deprivation. This lasted for at least a week until mice reached around 85% of their baseline weight. We confirmed that mice were comfortable licking to a syringe that delivered water while handling, indicating a reduction in overall stress and a willingness to seek water. After handling, mice were habituated on the rig for 2–3 days (3 days for the mice of the infusion and dopamine-inhibition experiments). We head-fixed the mice and immediately dispensed water reward. The 1st habituation session lasted for 15 min, and the subsequent habituation sessions lasted for 35 min (similar to the length of a behavioral session). We confirmed on the last habituation session that mice were comfortably licking to the spout. If mice never licked to the spout on that day, we continued the habituation until mice became comfortable licking.

For training in the stable task, we trained mice for at least 5 days. Mice were able to discriminate the cues on the first day and steadily improved (Fig. 1d, Extended Data Fig. 1a, c). Value memory for the stable task was tested after at least 5 days of training. For training on the dynamic task, mice were first trained on the stable task for 2–3 days and then on the stable task with the reversed contingency (odor A = no reward, odor B = reward) for 2–3 days. After this, mice were trained on the dynamic task, in which the reversal happened every session at trial 101; the number of dynamic-task sessions before each manipulation is given below. Sessions from the 9th dynamic-task session onward were considered expert sessions (Fig. 1f, g). The lick rates and AUROCs of Extended Data Fig. 1a-c come from 10 mice trained on the stable task for 3 sessions and then on the dynamic task as described above. The mice of Fig. 1e experienced their first reversal within the first dynamic-task session, without the intervening stage of reversed stable training. For the CaMKIIα conditional knockout mice and their controls (Fig. 5a-d, Extended Data Fig. 9a-h), all mice were trained, in order, on the stable task (sta1; 3 sessions), the stable task with the reversed contingency (sta2; 3 sessions), the dynamic task with one reversal per session (dyn1; 6 sessions) and the dynamic task with two reversals per session (dyn2; 6 sessions); in dyn1 and dyn2 the starting contingency alternated across days, and the last dyn2 session was taken as the expert session (Fig. 5c, d). For dopamine inhibition in the stable task (Fig. 3b), the light was delivered from the first stable-task session, which followed the licking habituation and the calibration session (see Dopamine optogenetic inhibition); the light was also delivered in the following 2–3 stable-task sessions. For the dynamic task, mice were trained for 10–16 sessions (about 2 weeks) and then ran light and no-light sessions on alternate days. For the KN-93 infusion experiments in the amygdala (Fig. 3i), KN-93 or saline was infused 0.5–1 h before each of the first three stable-task sessions, which immediately followed the habituation (day 1, Fig. 3j, l; days 1–3, Extended Data Fig. 6m-o). In the dynamic task, the infusions were made in expert mice after at least 2 weeks of the dynamic task, KN-93 and saline sessions alternating on consecutive days (1–2 sessions per drug and mouse). For the Emx1-cre × gtACR1 mice, mice were first trained on the stable task for 4–5 sessions and then underwent 1–2 silencing sessions of each type (cue or ITI; only the BLA mice underwent cue silencing), interleaved with sessions without light; the same mice were then trained on the dynamic task for ∼15 sessions and underwent 1–2 further silencing sessions of each type, again interleaved with sessions without light. For the OFC and vHip infusion experiments, mice were trained on the dynamic task for about 2 weeks and then received one infusion session per drug and region (saline, muscimol and KN-93; Extended Data Fig. 7h, j-l) on separate days, 1–2 days apart, while continuing the dynamic task. For the Neuropixels recordings, mice were trained on the four-block dynamic task for 11–12 sessions (after 2–3 sessions of the stable task, 2 sessions of the reversed stable task and 5–6 sessions of the one-reversal dynamic task) before the first recording; recordings were then performed on consecutive days, in runs of 1–7 sessions targeting one region and then the other (68 sessions from 7 mice: 30 amygdala and 38 hippocampal sessions; Extended Data Fig. 8a-c).

#### Testing value memory

We tested the stability of the value memory in two ways. In the first way, after mice were fully trained on the stable or dynamic task, we introduced breaks between sessions (1, 2, 4 or 8 days for the stable task, n = 4 mice; 1 day for the dynamic task, n = 7 mice). The break duration was randomized across mice. In the second way, after mice were fully trained on the stable or dynamic task, we inserted two long ITIs (37.5, 75, 150 or 300 s) before trials 50 and 150 of a session (see Behavioral tasks). For the stable task, each mouse experienced one long ITI of each duration over two sessions (n = 6 mice); for the dynamic task, 7–8 long ITIs per duration were collected from n = 7 mice (15 sessions). To quantify the mice’s ability to remember previously learned values across a long ITI, we computed the discrimination index as the difference in CS licks (number of licks during the 1.5-s odor) between the first CS+ and the first CS- presented after the long ITI, divided by the mean difference in CS licks between the CS+ and the CS- over the last five presentations of each cue before the long ITI (Fig. 1j). An index of 1 indicates that the discrimination was fully retained, 0 that it was lost and −1 that it was reversed. For the breaks between sessions, the discrimination index was the difference in CS licks between the first CS+ and the first CS- of the session following the break, divided by the mean difference in CS licks between the CS+ and the CS- over that same (post-break) session (Fig. 1k). For the stable– dynamic–retraining sequence (Extended Data Fig. 1e), the index was computed on the last session of each stage, i.e. after the overnight break, as the difference in CS licks between the first CS+ and the first CS- of the session, normalized by the maximal CS+ lick count in that session. A similar metric was used for the dopamine cue response, replacing the CS licks by the mean dF/F during the odor period (0–1.5 s from odor onset, 1-s pre-cue baseline subtracted; Fig. 1n; see Dopamine after long ITIs), and for the cue values of the RNNs (Fig. 2r; see Long ITIs in the RNNs).

### Behavioral analysis

#### Lick detection

Licks were recorded as spout-contact onsets registered by the digital input of the behavioral control system in the photometry, optogenetic, infusion and knockout experiments, and as infrared beam breaks during the Neuropixels recordings (see Behavioral setup). For the contact sensor, an onset that followed the previous one by less than 55 ms was treated as a double detection of the same lick and discarded (70 ms for the learning curves and learning constants of Fig. 1d–g). The only exception was the lick traces by presentation of each cue of the dopamine-inhibition experiment (Fig. 3d,e), for which every onset event was counted. For the beam-break sensor, a break of duration d was converted into max(1, round(8d)) licks spaced 125 ms apart from its onset, so that a bout during which the mouse held the beam broken was counted as licking at 8 Hz (see Alignment of spikes and licks). CS licks were defined as the number of licks during the 1.5-s odor (odor onset to offset); where a lick rate is reported (Fig. 5b–d, Extended Data Figs 7h, j–l and 9c, d, f, g) it is this count divided by 1.5 s, except in Fig. 4g, where the rate was taken 0.5–1.5 s after odor onset. US (consumption) licks were the licks in the 1 s after the outcome (Extended Data Figs 6c, l and 9h; see Reward consumption), and ITI licks the lick rate in the 2 s before odor onset (Extended Data Fig. 9e). The learning curves of Fig. 1d–f were binned by trial (5-trial bins, each cue averaged over its trials in the bin); all other lick curves were plotted against the presentation number of each cue, i.e. the trials on which that cue was shown. Where the axis of such a per-cue curve of licks or dopamine is nevertheless labelled "trial" (Fig. 1i; Fig. 3d, e; Extended Data Figs 1d, j and 6d, e, j, k), trial denotes the n-th presentation of that cue, counted separately for each cue, and not the trial number of the session. When a mouse contributed several sessions, they were averaged within the mouse before averaging across mice, unless the legend states that sessions are the unit. In the dynamic task the two cues were followed across the reversal: "CS+" denotes the cue rewarded after the reversal (the CS- before it) and "CS-" the cue rewarded before it. The reversal trial was taken from the block transition recorded by the behavioral control software, which marks the last trial of the first block, so the new contingency began on the following trial.

#### Cue discrimination

Discrimination between the cues was quantified as the area under the receiver operating characteristic curve (AUROC) of the single-trial CS licks (or lick rate) of CS+ against CS- trials, i.e. the probability that a randomly chosen CS+ trial had more licks than a randomly chosen CS- trial, ties counted as one half (0.5, no discrimination; 1, complete discrimination). The AUROC was computed over all 200 trials of day 1 of the stable task (Fig. 3h,l) or of each of days 1-3 (Extended Data Fig. 6n), over the 100 trials after the reversal in the dynamic task of the dopamine-inhibition and KN-93 experiments (Fig. 3h,l), and over all trials of a session with the CS+ defined by the current contingency (Fig. 5d, Extended Data Figs 1c and 9f). The difference between CS+ and CS- licks was computed over the same trials (Extended Data Fig. 6o), or, where stated, excluding the first 10 presentations of each cue in each block (Extended Data Figs 7k,l and 9g). For the optogenetic silencing experiments (Extended Data Fig. 7d,f), the difference between CS+ and CS- licks was computed separately for the light trials and for the interleaved trials without light of the same session, each session being one observation; in the dynamic task the first 10 trials after the reversal were left out. For cue silencing (Extended Data Fig. 7d) the light fell on the scored trial itself; for ITI silencing (Extended Data Fig. 7f) the scored trial was the one whose ITI was silenced. The same AUROC was computed from the single-trial cue dopamine responses of the dopamine-inhibition experiment (Extended Data Fig. 6f; see Dopamine cue response in the dopamine-inhibition experiment).

#### Learning constants of mice

The learning speed of a cue was estimated by fitting its CS licks with an exponential, y(j) = A∞ + (A0 − A∞)·exp(−j/τ), where j = 0, 1, 2, … counts the presentations of that cue (trials of the other cue do not advance the clock), A0 and A∞ are the initial and asymptotic lick counts and τ is the learning constant in cue presentations. τ was found by a grid search over log-spaced values, with A0 and A∞ solved by least squares at each τ; only solutions in the expected direction (rising for acquisition, falling for extinction) were accepted and the τ with the smallest squared error was taken. The grid spanned 1–1,000 presentations (100 values) for initial acquisition and the first reversal and 1–200 presentations (80 values) for the expert reversals. Initial acquisition (Fig. 1d,g) was fitted per mouse on the CS+ licks of days 1 and 2 of the stable task concatenated (400 trials). The first reversal (Fig. 1e,g) was fitted per mouse in mice that had never experienced a reversal (first dynamic-task session without a prior reversed stable stage), on the post-reversal trials of that session concatenated with the first block of the next session, which continued the same contingency (about 200 trials), for the acquisition of the new CS+ and the extinction of the old CS+. Expert reversals (Fig. 1f,g) were fitted per session on the 100 trials after the reversal of dynamic-task sessions 9 and later. Learning constants are reported as medians with 95% bootstrap confidence intervals of the median (2,000 resamples), on a logarithmic axis. The same fit was applied to the per-trial cue dopamine response (Extended Data Fig. 1h; see Dopamine learning curves and learning constants) and, with the grid starting at 0.5 presentations, to the value readout of the RNNs (Fig. 2b, c; Extended Data Fig. 2d; see Learning constants for RNNs).

#### Meta-learning speed-up in mice

The speed-up of learning with experience in mice, the reference for the RNN speed-up of Fig. 2d, was defined as the ratio of the median initial learning constant to the median expert learning constant of the new CS+. For mice this was 73/1.7 = 43. Its confidence interval was obtained by resampling mice (20,000 resamples), each resampled mouse contributing its initial fit and all of its expert sessions. For the RNNs, whose initial and expert constants come from different networks, the two sets of constants were resampled independently (20,000 resamples; see Meta-learning speed-up in RNNs).

#### Discrimination index after long ITIs and breaks

Sessions of the long-ITI test contained two long ITIs, preceding trials 50 and 150, whose durations, including the 2.5 s that followed the outcome, were 37.5, 75, 150 or 300 s. For each long ITI, the discrimination index (Fig. 1j) was the difference in CS licks between the first CS+ and the first CS- presentation after the ITI, divided by the difference between the mean CS licks of the last five CS+ and the last five CS- presentations before it, so that 1 means that the discrimination was fully retained, 0 that it was lost and −1 that it was reversed (by design, the first cue after a long ITI was almost always the CS-, 52 of 54 long ITIs). The stable-task mice had one long ITI of each duration (two sessions per mouse), the dynamic-task mice 7–8 per duration; the tasks were compared at each duration. Fig. 1i shows the CS licks on the successive presentations of each cue around the long ITIs of the sessions whose first long ITI lasted 150 or 300 s. The same index was computed from the mean cue dopamine response (mean dF/F 0–1.5 s from odor onset minus the 1-s pre-cue baseline) instead of CS licks (Fig. 1n; see Dopamine after long ITIs). After a break (Fig. 1k), the index was the difference in CS licks between the first CS+ and the first CS- trial of the session following the break, divided by the difference between the mean CS+ and mean CS- licks over that whole session; the stable task was tested after breaks of 1, 2, 4 and 8 days, the order of the break durations being randomized across mice, and the dynamic task after the 1-day break of the daily schedule. In Extended Data Fig. 1e the first-trial index was instead normalized by the maximal CS+ lick count of that session and computed on the last session of each stage (stable, dynamic and retraining), i.e. after the overnight break; CS licks were the licks during the odor (55-ms debouncing), and CS+ trials were the trials on which the reward was delivered. The stages were compared within mice with a two-tailed paired t-test (stable versus dynamic and dynamic versus retraining; n = 6 mice).

#### Early and late windows in the manipulation experiments

For the dopamine-inhibition and KN-93 experiments, licking to each cue was compared between two five-trial windows (Fig. 3f,g,j,k). In the stable task (day 1), "early" was trials 1–5 and "late" the last five trials of the session; in the dynamic task, "pre" was the five trials before the reversal and "post" trials 16–20 after it (dopamine inhibition, Fig. 3g) or the last five trials of the 100-trial post-reversal block (KN-93, Fig. 3k). A cue’s value in a window was the mean CS licks on the trials of the window on which that cue was presented; a mouse without a presentation of that cue in a window was left out of that point, and sessions were averaged per mouse. The cue dopamine response was not compared between windows; it was followed by presentation of each cue and quantified by its AUROC (Extended Data Fig. 6d-f; see Dopamine cue response in the dopamine-inhibition experiment).

#### Statistics

All tests were two-tailed and no correction for multiple comparisons was applied, except in Extended Data Fig. 4f,g, where the pairwise comparisons were Holm-corrected. Comparisons between cohorts of different mice used Student’s t-test (Fig. 1j,k,n; stable task of Fig. 3h,l and Extended Data Fig. 6f, n, o) or the Mann–Whitney U test (Fig. 5d, Extended Data Fig. 9e–h, and all comparisons between groups of networks), and comparisons within mice (or within sessions, or within long ITIs) used the paired t-test (Fig. 3f,g,j,k; dynamic task of Fig. 3h,l and Extended Data Fig. 6f; Extended Data Figs 1e, 6b,c,l and 7d,f,k,l) or the Wilcoxon signed-rank test (Fig. 4g– k and the paired network comparisons of Fig. 2p and Extended Data Figs 4m and 5f–j). Whether the effect of a manipulation differed between the two tasks (Fig. 3h,l; Extended Data Fig. 6f) was tested with a label-permutation test: the statistic was the effect (control minus manipulated) in the stable task minus the mean within-mouse effect in the dynamic task; its null distribution was generated by shuffling the group labels of the mice in the stable task and, independently, swapping the two conditions within each mouse in the dynamic task, 20,000 times, and P was (number of permuted statistics at least as extreme as the observed one + 1)/(20,000 + 1). Confidence intervals of medians and ratios are percentile bootstrap intervals (2,000 resamples for the learning constants, 20,000 for the meta-learning speed-up; 500 resamples for the bootstrap s.e.m. of the AUROC in Extended Data Fig. 3). Further tests, named in the legends, were a two-tailed Mann–Whitney U test between the initial and expert dopamine learning constants (Extended Data Fig. 1h), Student’s t-test between the per-session silencing effects of the two tasks (Extended Data Fig. 7f), chi-square tests of the fractions of coding units across subregions and regions and a binomial test of the fraction of value units excited by the CS+ against 0.5 (Extended Data Fig. 8f–j), and the Wilcoxon signed-rank test for decoding accuracies (Extended Data Fig. 8l, m) and for the within-network comparisons of Extended Data Figs 4n and 5. Unless stated otherwise, data are mean ± s.e.m., and *, ** and *** denote P < 0.05, 0.01 and 0.001. The unit of n is stated in each legend: mice for comparisons between cohorts and within-mouse comparisons (sessions averaged per mouse), sessions or long ITIs where each is a separate observation, and networks (independent random initializations) for the RNNs. Analyses were performed in MATLAB and Python (NumPy, SciPy, scikit-learn).

### Fiber photometry

#### Fiber photometry

We used a commercially available bundle fiber photometry system (BFMC, Doric) to record photometry signals from multiple animals simultaneously. A low-autofluorescence Branching Bundle Patchcord (400 μm, 0.57 NA) was connected to the photometry system. Each end of the fiber went to a single behavioral rig, allowing us to independently track photometry signals from up to 3 animals simultaneously. A blue excitation LED (470 nm, 10 μW power at the tip of the 400-μm patchcord ferrule) was used to collect the GRABDA3m signal. A purple excitation LED (415 nm, 9 μW power) was used to collect the control signal for correcting movement artifacts. The following parameters were used for imaging in the Doric Neuroscience Studio V6 software: power = 10%, framerate = 30 Hz, exposure = 0.012 s, gain = 9.9 dB. Dopamine was recorded in wild-type mice expressing GRABDA3m in the left VS (see Viruses) through an optical fiber implanted above the injection site (200 μm, 0.66 NA, Doric; fiber tip locations in Extended Data Fig. 1i). In the dopamine-inhibition experiments the same sensor was recorded through a 200-μm fiber (see Dopamine optogenetic inhibition). Trial start times were registered on the photometry system through a TTL pulse from the Bpod at the start of every trial, and cue onset within each trial was taken from the Bpod state transitions.

#### Photometry preprocessing

The fluorescence of each channel (470 nm and 415 nm; sampled at 30 Hz) was converted to dF/F = (F − F0)/F0, where F0 was the 10th percentile of the raw signal in a 30-s rolling window (evaluated every 1 s and linearly interpolated). Samples outside the 0.01st–99.99th percentiles of each channel were treated as outliers and replaced by linear interpolation. Both channels were then linearly interpolated onto a common 1-kHz time base, the 470-nm dF/F was regressed linearly on the 415-nm dF/F, and the fitted prediction was subtracted from the 470-nm dF/F. This isosbestic-corrected dF/F was used in all analyses without further smoothing or normalization, except that the example sessions of Extended Data Fig. 1j show the corrected dF/F z-scored over the session.

#### Dopamine cue response

For the long-ITI experiment (Fig. 1m, n) and the learning curves (Extended Data Fig. 1f–h), the corrected dF/F was aligned to odor onset in 10-ms bins, the mean over the 1 s preceding odor onset was subtracted trial by trial, and the cue response was defined as the mean of the baseline-subtracted dF/F over the 1.5-s odor period (0–1.5 s from odor onset). For the dopamine-inhibition experiment (Extended Data Fig. 6d-f), the cue response was the mean of the corrected dF/F over the 1.5 s from cue onset without baseline subtraction; because the light was delivered at the outcome (Fig. 3c), this window preceded the light on every trial.

#### Dopamine learning curves and learning constants

Sessions were selected from the training log. Initial acquisition (Extended Data Fig. 1f) used the first two odor sessions (day 1 and day 2 of the stable task) of wild-type sensor mice that never received an optogenetic manipulation (n = 5 mice). Expert reversals (Extended Data Fig. 1g) used dynamic-task sessions from training day 10 or later (counting all dynamic-task sessions), excluding days with a drug infusion or any other manipulation, up to the first five recorded sessions per mouse. Responses were averaged in 5-trial bins for display (Extended Data Fig. 1f, g; the trial axis counts the trials of the session, both cues, each cue averaged over its trials in the bin), and expert sessions were averaged within each mouse before averaging across mice. Learning constants (Extended Data Fig. 1h) were obtained by fitting the per-trial cue response of one cue with y(t) = a − b·exp(−t/τ) for acquisition and y(t) = a + b·exp(−t/τ) for extinction, where t = 0, 1, 2, … counts the presentations of that cue (the other cue’s trials do not advance the clock). τ was found by grid search over a logarithmic grid (1–1,000 presentations, 100 values, for initial acquisition; 1–200 presentations, 80 values, for expert sessions), with a and b fitted by least squares for each τ and b constrained to be positive. Initial acquisition was fitted per mouse on the concatenated day 1 and day 2 (CS+ only); expert acquisition (new CS+) and extinction (old CS+) were fitted per session on the post-reversal trials. We report medians with 95% bootstrap confidence intervals of the median (2,000 resamples).

#### Dopamine after long ITIs

Dopamine was recorded during long-ITI sessions of expert mice on the stable (n = 6 mice, two sessions each) or dynamic (n = 7 mice, two or three sessions each) task. Each session comprised 200 trials with a long ITI (37.5, 75, 150 or 300 s) before trials 50 and 150 (see Testing value memory), and each mouse received every duration. By design, the first cue after a long ITI was the CS− (52 of 54 long ITIs). For each long ITI and each cue, we computed the mean cue response over the last five presentations before the ITI (before) and the response on the first presentation after it (after). Fig. 1m shows the cue-aligned traces of the 150- and 300-s ITIs pooled. The dopamine discrimination index (Fig. 1n) was defined as for licking (see Testing value memory): the difference between the CS+ and CS− responses on their first presentations after the ITI divided by the corresponding difference over the last five presentations before it (1, discrimination fully retained; 0, lost; −1, reversed). Because ratios of single-trial responses are heavy-tailed, each index was clipped to [−1, 1] before averaging (mean ± s.e.m. over the long ITIs of each duration; stable: 6 per duration; dynamic: 7–8 per duration), and the two tasks were compared at each duration with a two-tailed unpaired t-test, as in Fig. 1j.

#### Calibration of dopamine inhibition

To calibrate the light power for each DAT-cre × gtACR1 mouse (Extended Data Fig. 6b), we recorded dopamine in the VS during a session of 100 free water rewards delivered at intervals of about 15 s (10–24 s; 1.5 μL), every 10th reward was paired with the 595-nm light at the outcome (10 light rewards per session), delivered as in the task (0.5 s of constant light followed by a 1-s ramp; see Dopamine optogenetic inhibition). For this analysis the 470-nm and 415-nm dF/F were used without the isosbestic regression described above and aligned to reward delivery. Because the orange light produced an artifact in both channels, on light trials the 415-nm dF/F, scaled by the ratio of the baseline fluorescence of the two channels (F0,415/F0,470), was subtracted from the 470-nm dF/F; no-light trials were used without subtraction. Traces were baseline-subtracted (mean over 2.5–1 s before the reward, averaged over the mice of each condition), and the reward response was the mean dF/F over the 1.5 s after reward delivery, averaged over the light and over the no-light rewards of each mouse and compared with a two-tailed paired t-test (n = 13 mice). The light power was adjusted for each mouse so that the phasic response to the unexpected reward was inhibited while minimizing the suppression beyond baseline (see Dopamine optogenetic inhibition).

#### Dopamine cue response in the dopamine-inhibition experiment

The cue response on each trial (mean dF/F over the 1.5-s odor; see Dopamine cue response) was followed by presentation of each cue (Extended Data Fig. 6d, e). In the stable task (day 1; control, n = 5 mice with a GRAB-DA3m fiber; DAT-cre, n = 9 mice), presentations 1–100 of each cue were smoothed per mouse with a centred running mean of 3 presentations. In the dynamic task, the responses to each odor were aligned to the reversal (the last 10 presentations before it and the first 50 after it, within the 100 trials after the reversal), CS+ denoting the odor rewarded after the reversal (paired with the light in the light sessions) and CS- the odor rewarded before it. Not every behavioral session had a photometry recording; 23 light sessions from 10 mice and 20 no-light sessions from 12 mice were analysed, and sessions were averaged within each mouse. Discrimination by the cue dopamine response (Extended Data Fig. 6f) was the AUROC between the single-trial cue responses of the CS+ and CS- trials of a session (stable: all trials of day 1; dynamic: the 100 trials after the reversal, CS+ being the odor rewarded in that block), averaged within each mouse. The stable task was compared between mice (two-tailed Student’s t-test), the dynamic task within mice (two-tailed paired t-test on the 10 mice with both session types), and the difference of the light effect between the tasks with the label-permutation test (20,000 permutations; see Statistics).

### Causal manipulations

#### Dopamine optogenetic inhibition

We performed optogenetic experiments to test the role of phasic dopamine during the outcome period in the stable and the dynamic task. Optical fibers (400 μm, 0.66 NA, Doric) were implanted bilaterally above the VTA in DAT-cre × gtACR1 mice. The VTA was targeted at AP −2.9 mm and ML ±1.75 mm relative to bregma, with an insertion depth (from the brain surface) of 4.0 mm along a trajectory angled 15° from vertical toward the midline. To confirm the inhibition of the dopamine response, we expressed GRAB-DA3m in the ventral striatum (1.0/1.5/3.8 mm; see Surgeries and Viruses) and implanted an optical fiber (200 μm, 0.66 NA, Doric) above the injection site (DV 3.6 mm) in the same surgery. Control mice (n = 7, five of them with a GRAB-DA3m fiber; heterozygous for the GtACR1 reporter allele and negative for DAT-IRES-Cre) did not carry the opsin and received the same light through the same fiber implants (Extended Data Fig. 6a). The light (595 nm; fiber-coupled LED, Thorlabs) was delivered at the outcome as a 1.5-s waveform: 500 ms at constant amplitude, after which the amplitude was ramped down linearly to zero over 1 s (Fig. 3c). We first calibrated the light power for each mouse so that the phasic dopamine response to an unexpected reward was abolished while minimizing the suppression of dopamine below baseline, because suppression below baseline has been shown to elevate protein kinase A signaling in D2-receptor-expressing cells^33,34^, which would add a negative prediction error to the removal of the phasic response. The calibration session was run on the day before the first task session: 100 unexpected rewards (no odor) were delivered at intervals of about 15 s (10-24 s), the light was paired with every tenth reward (rewards 10, 20, …, 100). The dopamine response (mean dF/F over the 1.5 s after the reward, baseline subtracted) was compared between the rewards with and without light (Extended Data Fig. 6b; n = 13 mice). The light power at the fiber tip was 1–3 mW. The light artifact in the photometry signal of the light trials was removed by subtracting the isosbestic (415 nm) channel scaled by the ratio of the baseline fluorescence of the two channels (F0,415/F0,470) of each session, a factor that needs no fitting; in the opsin-negative control mice this correction made the reward responses of light and no-light trials coincide. In the stable task, dopamine was inhibited on day 1, the first session after licking habituation (Fig. 3b): from the first trial of the session, the light was delivered at the outcome of the CS+ trials and never on CS- trials (Fig. 3c). One session per mouse (day 1) was analysed (control, n = 7; DAT-cre, n = 9 mice). In the dynamic task, the DAT-cre mice (n = 13) were trained for 10-16 sessions of the dynamic task (about two weeks, after the stable and the reversed stable sessions) and then ran light and no-light sessions on alternating days (26 light sessions from 11 mice and 31 no-light sessions from 13 mice). In a light session the light was delivered at every reward of the block after the reversal, that is at every reward of the new CS+, and never before the reversal or on unrewarded trials, because the light was triggered by the reward-delivery state of the trial (Fig. 3c). Licks were counted during the 1.5-s odor (CS licks) and during the 1 s after the outcome (US licks; Extended Data Fig. 6c; see Reward consumption); lick events closer than 55 ms to the previous one were discarded. Discrimination was quantified as the AUROC between the CS licks of CS+ and CS- trials, over all 200 trials of the day-1 session in the stable task and over the 100 trials after the reversal in the dynamic task (CS+, the cue rewarded in that block), with sessions averaged within each mouse. Early and late windows were the first and the last five trials of the session (stable) or the five trials before the reversal and trials 16-20 after it (dynamic). The stable task was compared between mice (two-tailed Student’s t-test) and the dynamic task within mice (two-tailed paired t-test on the 11 mice with both session types); the difference of the light effect between the two tasks was tested with a label-permutation test (20,000 permutations). The dopamine response to the cue was the mean dF/F over the 1.5-s odor (Extended Data Fig. 6d-f; see Dopamine cue response in the dopamine-inhibition experiment). CS licks were also plotted against the presentation number of each cue (Fig. 3d, e): in the stable task, presentations 1–100 of each cue on day 1, smoothed per mouse with a centred running mean of 3 presentations; in the dynamic task, the last 10 presentations of each cue before the reversal and the first 50 after it (within the 100 trials after the reversal), without smoothing; sessions were averaged within each mouse before averaging across mice.

#### KN-93 infusion into the amygdala

For inhibiting synaptic plasticity in the amygdala, we used KN-93, a CaMKII inhibitor known to disrupt synaptic plasticity^13,84^. Once mice were fully habituated to the rig (for testing in the stable task) or fully trained on the dynamic task (for testing in the dynamic task), the day before the infusion, two small craniotomies were made above the amygdala on each hemisphere (−1.2/3.3/4.2 mm) and sealed with silicone elastomer (Kwik-Cast, World Precision Instruments). On the day of infusion, mice were lightly anaesthetized with isoflurane (1%). KN-93 (water-soluble KN-93 dissolved in saline at 100 μM; 422711-1MG, Millipore Sigma) was injected bilaterally into the amygdala (300 nl per site). Behavioral testing started 0.5–1 h after the infusion. To verify the injection site, KN-93 was mixed with a lipophilic tracer DiI (less than 5% of the total volume; V22889, ThermoFisher Scientific). Mice were perfused (see Histology) and the slices were imaged under a wide-field microscope (Extended Data Fig. 6g-i). For saline infusion, the same procedure was performed with saline + DiI. In the stable task, different mice received saline (n = 11) or KN-93 (n = 8) before the day-1 session (Fig. 3j, l; Extended Data Fig. 6j, l). To follow the recovery of learning, mice were infused again, with the same drug and dose, before each of the sessions of days 2 and 3 (saline, n = 8; KN-93, n = 5 mice; Extended Data Fig. 6m-o). In the dynamic task, a separate group of mice (n = 5) was trained for at least 2 weeks on the dynamic task and then received KN-93 and saline before sessions on alternating days (7 KN-93 and 9 saline sessions in total), each session with one reversal at trial 101 (Fig. 3k, l; Extended Data Fig. 6k, l). Licks were counted and discrimination was quantified as for the dopamine inhibition experiments: AUROC over all 200 trials of day 1 (stable) or over the 100 trials after the reversal (dynamic; sessions averaged within each mouse); early and late windows were the first and last five trials of day 1, and the five trials before the reversal and the last five trials of the post-reversal block. The stable task was compared between mice (two-tailed Student’s t-test, also for days 1-3 in Extended Data Fig. 6n, o) and the dynamic task within mice (two-tailed paired t-test); the difference of the KN-93 effect between the tasks was tested with a label-permutation test (20,000 permutations). The CS licks were also plotted against the presentation number of each cue: the first 85 presentations of each cue on day 1 of the stable task (Extended Data Fig. 6j) and, in the dynamic task, the last 10 presentations before and the first 49 after the reversal (Extended Data Fig. 6k; mean ± s.e.m. over sessions). Over days 1–3 (Extended Data Fig. 6m), the CS licks of the first 100 presentations of each cue on each day were averaged in bins of 10 presentations in the mice that completed all three days, and the AUROC and the difference between CS+ and CS- licks (Extended Data Fig. 6n, o) were computed over all 200 trials of each day’s session.

#### Reward consumption

To check that the manipulations did not prevent the mice from drinking, we counted the US (consumption) licks, the licks in the 1 s after the outcome (from the start of the post-outcome state, which began when the reward valve closed or, on unrewarded trials, at the time of the omitted outcome; 55-ms debouncing), on CS+ (rewarded) and CS- trials. In the dopamine-inhibition experiment (Extended Data Fig. 6c) they were averaged over all trials of day 1 of the stable task for each mouse (control, n = 7; DAT-cre, n = 9 mice) and, in the dynamic task, over the 100 trials after the reversal (the block in which the light was given), sessions averaged within each mouse, for the light and the no-light sessions of the same DAT-cre mice (26 light sessions from 11 mice and 31 no-light sessions from 13 mice). In the KN-93 experiment (Extended Data Fig. 6l) they were averaged over all trials of day 1 for each mouse in the stable task (saline, n = 11; KN-93, n = 8 mice) and over all trials of each session in the dynamic task, each session being one observation (9 saline and 7 KN-93 sessions from 5 mice). CS+ and CS- trials were compared within each group with a two-tailed paired t-test.

#### Muscimol and KN-93 infusions into the OFC and ventral hippocampus

To test whether the OFC or the ventral hippocampus (vHip), which project to the amygdala, supports the reversal in the dynamic task, wild-type mice (n = 8) were trained on the dynamic task (two stable sessions, two reversed stable sessions and about 2 weeks of the dynamic task) and then received an infusion of saline, muscimol or KN-93 before a dynamic-task session, one session per mouse, region and drug (48 sessions), each with a single reversal at trial 101. Small craniotomies above the OFC and the vHip were made at the head-plate surgery and sealed with silicone elastomer. Infusions were made under light isoflurane anesthesia as for the amygdala: 200–300 nl per site of saline, muscimol (1 ng nl⁻¹) or KN-93 (100 μM) was injected into the OFC (2.6/1.9/1.8 mm AP/ML/DV) or the vHip (−3.4/3.2 mm AP/ML; DV 4.2 mm from the brain surface), and the session started ∼1h later. Muscimol was infused bilaterally into the vHip and unilaterally into the OFC in six of the eight mice (bilaterally in the other two); saline and KN-93 were always bilateral. In some sessions a lipophilic tracer (DiI, DiD or DiO) was added to the infused solution and the injection sites were identified from the histology (Extended Data Fig. 7g, i). Anticipatory licking was quantified as the lick rate during the 1.5-s odor (licks per s; 55-ms debouncing as above) as a function of the presentation number of each odor relative to the reversal (Extended Data Fig. 7h, j). Discrimination was the difference in lick rate between the CS+ and the CS- under the current contingency, before and after the reversal, excluding the first ten presentations of each odor in each block (Extended Data Fig. 7k, l), and each drug was compared with saline within mice (two-tailed paired t-test, n = 8). In Extended Data Fig. 7h, j, presentation 0 is the first presentation of each odor after the reversal, and the last 10 presentations before and the first 30 after the reversal are shown (mean ± s.e.m. over mice).

#### Optogenetic silencing of the basolateral amygdala, ventral hippocampus and ALM

We performed optogenetic experiments to test the role of the basolateral amygdala (BLA), the vHip and the anterior lateral motor cortex (ALM) in the stable and the dynamic task. Optical fibers were implanted bilaterally above the BLA (400 μm, 0.66 NA, Doric; coordinates −1.2/3.3/4.0; n = 6 mice), the vHip (400 μm, 0.66 NA, Doric; coordinates −3.4/3.2/3.3; n = 7 mice, 6 of which were also tested in the dynamic task) or the ALM (400 μm, 0.66 NA, Doric; coordinates 2.5/1.27 mm, just above the brain surface; n = 4 mice) in Emx1-cre × gtACR1 mice (Extended Data Fig. 7b, c, e). Mice were allowed to recover for at least 2 weeks before starting the handling procedure. Mice were first trained on the stable task for 4-5 sessions and then underwent 1-2 silencing sessions of each type (cue or ITI; only the BLA mice underwent cue silencing); the same mice were then trained on the dynamic task for about 15 sessions and underwent 1-2 further silencing sessions of each type (session numbers in the legend of Extended Data Fig. 7d, f). A 470-nm LED was used to activate gtACR1. The light was delivered on about 15% of the trials (30 trial numbers drawn at random between trials 10 and 190, giving 25-30 light trials per session), randomly interleaved with trials without light. For testing the role of the ITI activity, the light was on for 3 s at the end of the ITI and ended 1 s before odor onset (0.5–1 mW at the fiber tip). For testing the role of the cue- period activity of the BLA, the light was on from odor onset for 3 s at 0.5–1 mW. Licks were counted during the 1.5-s odor as above. For each session, the effect of silencing was quantified as the difference between CS+ and CS- licks (and as the AUROC between CS+ and CS- licks) on the light trials against the interleaved trials without light; in the dynamic task the CS+ was the cue rewarded in the current block and the first 10 trials after the reversal were excluded. Light and control trials were compared across sessions with a two-tailed paired t-test (Extended Data Fig. 7d, f).

### CaMKIIα conditional knockout

**CaMKIIα conditional knockout mice. To conditionally delete CaMKIIα in Emx1-cre-expressing cells, Emx1-IRES-Cre mice (JAX stock** no. 005628) were crossed with mice carrying a floxed Camk2a allele (JAX stock no. 006575) (see Animals). Conditional knockout mice (cKO) were homozygous for the floxed Camk2a allele and heterozygous for the Emx1-IRES-Cre allele (Camk2a^fl/fl; Emx1-Cre/+); Cre-negative littermates homozygous for the floxed Camk2a allele (Camk2a^fl/fl; Emx1+/+) served as controls (Fig. 5a). Genotypes were determined by Transnetyx. Mice were trained in two cohorts of 5 control and 5 cKO mice each (control: n=10 mice; cKO: n=10 mice in total; males and females; 2–4 months old at the start of training). Water deprivation, handling and habituation to the rig followed the procedure described in Behavioral training.

#### Training of the CaMKIIα cKO mice

All mice went through the same four stages in a fixed order, one session per day on consecutive days (Extended Data Fig. 9b). (1) Stable task (sta1): 3 sessions of 200 trials in which odor A was always rewarded and odor B never rewarded, as in the stable task (see Behavioral tasks); this stage is the stable task of Fig. 5b (days 1–3). (2) Stable task with the reversed contingency (sta2): 3 sessions of 200 trials in which odor B was rewarded and odor A was not. (3) Dynamic task with one reversal per session (dyn1): 6 sessions of 200 trials with a reversal of the contingency at trial 101 (2 blocks of 100 trials). (4) Dynamic task with two reversals per session (dyn2): 6 sessions of 210 trials with reversals at trials 71 and 141 (3 blocks of 70 trials). In the dynamic stages the rewarded odor of the first block alternated between consecutive sessions, so that the first block of each dyn1 session continued the contingency of the last block of the previous session. The last dyn2 session (day 6) is the "expert" dynamic session of Fig. 5c, d and Extended Data Fig. 9d. Odors were drawn pseudo-randomly with equal probability on every trial, the outcome was delivered at odor offset (1.5-s odor) and the ITI followed the truncated exponential distribution of the other behavioral experiments (see Behavioral tasks); the reward volume was 1.5 μL.

#### Behavioral analysis of the cKO mice

Licks were detected as the rising edges of the lick sensor signal and were debounced with a 55-ms refractory period. For every trial we computed the anticipatory lick rate (licks/s during the 1.5-s odor), the ITI lick rate (licks/s in the 2 s before odor onset) and the outcome lick rate (licks/s in the 1 s after the outcome, that is, after the end of the odor; reward consumption on CS+ trials and licking after omission on CS- trials). Discrimination between CS+ and CS- was quantified per session as the area under the receiver operating characteristic curve (AUROC) of the anticipatory lick rate between all rewarded and all unrewarded trials of the session (Fig. 5d, Extended Data Fig. 9f); for the stable task of Fig. 5d the AUROC was averaged over the sta1 sessions of each mouse (days 1–3), and for the dynamic task it was taken from the last dyn2 session. Lick rates as a function of presentation number (Fig. 5b) were computed per mouse by binning the first 100 presentations of each odor in each sta1 session into bins of 10 presentations, and then averaged over mice. Lick rates around the reversal (Fig. 5c) were aligned to the last 10 presentations of each odor before and the first 30 presentations after each reversal of the last dyn2 session, averaged over the two reversals of the session for each mouse and then over mice. For the per-session CS+ and CS- lick rates of Extended Data Fig. 9g, the first 10 presentations of each odor in each block were excluded so that the rates reflect the asymptotic, rather than the transitional, licking of each block. The ITI lick rate (Extended Data Fig. 9e) was averaged over all trials of a session and then over the sessions of each stage for each mouse; the outcome lick rate (Extended Data Fig. 9h) was averaged separately over the rewarded and unrewarded trials of each session. For the example sessions (Extended Data Fig. 9c, d) the single-trial anticipatory lick rates of each odor were plotted without smoothing. Control and cKO mice were compared for each session, stage or task with a two-tailed Mann–Whitney U test (n=10 mice per group unless stated otherwise), without correction for multiple comparisons; all data are mean ± s.e.m. over mice.

#### CaMKIIα immunohistochemistry

At the end of behavioral testing, mice were perfused and the brains were sectioned as described in Histology. To verify the loss of CaMKIIα in the cKO mice, free-floating sections were immunostained for CaMKIIα (mouse monoclonal CaMKII-α (6G9) antibody, #50049, Cell Signaling Technology; 1:500; overnight at 5 °C), followed by an Alexa Fluor 647-conjugated secondary antibody (1:500; room temperature), and mounted in VECTASHIELD mounting medium with DAPI. Sections were imaged with the slide scanner described in Histology (Zeiss Axioscan 7), and one coronal section at AP −1.6 mm from bregma is shown for one control and one cKO mouse (Extended Data Fig. 9a; DAPI shown in green and CaMKIIα in magenta).

### Neuropixels recordings and analysis

#### Neuropixels recordings

Acute Neuropixels recordings were made in 7 mice (wild-type C57BL/6J) trained on the dynamic task (see Behavioral training). A head plate was attached as described above (see Surgeries). The day before the first recording session, small craniotomies were made above the amygdala and the hippocampus (−1.2/3.3 mm AP/ML for the amygdala and −3.4/3.2 mm AP/ML for the hippocampus; left hemisphere only) and sealed with silicone elastomer (Kwik-Cast, World Precision Instruments). Each mouse was recorded once per day for up to 12 days (7-12 sessions per mouse, 68 sessions in total), with one single-shank Neuropixels 1.0 probe per session targeting either the amygdala (30 sessions) or the hippocampus (38 sessions); consecutive sessions in the same mouse alternated between the two targets in runs of one to seven days. On each recording day the silicone was gently removed to expose the brain, the probe was soaked in DiO, DiI or DiD solution (Vybrant Multicolor Cell-Labeling Kit, ThermoFisher Scientific) to track its position and mounted on a vertical manipulator (KDC101 and Z825B, Thorlabs). The probe was slowly lowered until it touched the brain surface; saline was applied around the craniotomy to prevent drying and to help the insertion through the dura. The probe was inserted through the dura at 0.05 mm s⁻¹ and, once it had penetrated the brain, at 0.01 mm s⁻¹, to up to 5.3 mm below the brain surface for amygdala sessions and up to 4.4 mm for hippocampal sessions, and left to settle for 10 min before the session. After the session the probe was retracted at 0.01 mm s⁻¹ and silicone was applied on the craniotomy. All recordings were performed in SpikeGLX (https://github.com/billkarsh/SpikeGLX) with a sampling rate of 30 kHz, local field potential gain of 250 and action potential gain of 500, and we analyzed only the action potential channel (high-pass filtered in hardware with a cut-off frequency of 300 Hz). Behavioral and neural recordings were synchronized using a transistor-transistor logic (TTL) pulse sent from the Bpod to the PXIe acquisition module SMA input at the start of every trial (high during the inter-trial-interval state; see Alignment of spikes and licks).

#### Task during the recordings

The recording sessions used the dynamic task with the trial structure described above (see Behavioral tasks) and four blocks per session (Fig. 4a). A trial began with an inter-trial interval drawn from a truncated exponential distribution (5–11 s from the previous outcome to the next odor, including the 2.5-s post-outcome period), followed by one of two familiar odors ((S)-(−)-limonene and 1-heptanol, as above; 1.5 s, 50% each), the outcome at odor offset (water reward, 1.5–2 μL, or no reward) and the 2.5-s post-outcome period. Within a block one odor was rewarded on every presentation and the other never, and the contingency reversed at every block transition; the odor rewarded in the first block varied across sessions (odor 1 in 37 and odor 2 in 31 sessions). Sessions had either 240 trials (four blocks of 60 trials; 41 sessions) or 300 trials (four blocks of 50 trials followed by two blocks of 50 trials with a novel pair of odors; 27 sessions). In a subset of sessions, the inter-trial interval preceding the last trial of blocks 2 and 3 (trials 120 and 180, or 100 and 150) was lengthened to 87.5 s (90 s from the previous outcome to the odor; Fig. 4a), so that the trial after each long inter-trial interval was the last trial of its block, one trial before the reversal. Licking was detected with an infrared emitter-photodiode pair across the water spout (see Behavioral setup).

#### Histology and probe registration

Mice were perfused and the brains sectioned and imaged as described in Histology. The sections with the dye tracks were registered to the Allen Mouse Brain Atlas with AP_histology (https://github.com/petersaj/AP_histology), the coordinates of each probe track were converted to the format of the IBL Ephys Atlas alignment tool (https://github.com/int-brain-lab/iblapps/tree/master/atlaselectrophysiology), and the track was aligned to the electrophysiological features of the recording to assign atlas coordinates and a brain region to each of the 383 located channels. Each single unit was assigned the location of its peak channel (units on channel 383, which has no registered location, took the location of channel 382). Units were grouped into the amygdala (basolateral, BLA; lateral, LA; basomedial, BMA; central, CEA; piriform-amygdalar, PAA; cortical, COA; and posterior, PA, amygdalar nuclei, pooled over their subdivisions), the hippocampus (CA1, CA3 with CA2, dentate gyrus, DG; prosubiculum, ProS; subiculum, SUB, with presubiculum; and entorhinal cortex, ENT) and the basal ganglia (caudoputamen, CP, 1,786 units; unassigned striatum, STR, 351 units; external globus pallidus, GPe, 51 units; and pallidum, PAL, 4 units). A session was classified as an amygdala or a hippocampal session according to whether more of its units were located in the amygdala or in the hippocampus. All basal ganglia units were recorded in amygdala sessions (29 of 30 sessions), in the posterior striatum and adjacent pallidum (median AP −1.6 mm). Probe tracks (the 383 located channels of each session) were rendered on the 25-µm Allen atlas (Fig. 4b and Extended Data Fig. 8a, b); the coronal slices of Extended Data Fig. 8b are at the median anterior-posterior position of the tracks of each target (AP −1.6 mm and −3.4 mm).

#### Spike sorting and curation

Neuropixels recordings were spike sorted offline with Kilosort4 (https://github.com/MouseLand/Kilosort) with default parameters, followed by manual curation of individual units in phy (https://github.com/cortex-lab/phy; version 2.1.0). Clusters labelled ’good’ by Kilosort were inspected in descending order of amplitude, labelled as single units or noise, and merged where one unit had been split. All clusters labelled as single units after curation (34-659 per session) entered the analyses; no firing-rate threshold was applied except in the selection of the dPCA regularization parameter (units with a mean rate below 0.1 Hz excluded) and of the example units (rate ≥2 Hz; Extended Data Fig. 8d, e).

#### Alignment of spikes and licks

Spike times were converted from the probe clock to the behavioral clock by piecewise-linear interpolation between the synchronization pulses that mark the start of every trial. Licks were recorded as the onset and offset of each beam break; because mice sometimes held the beam broken for the duration of a lick bout, every break of duration d was counted as max(1, round(8d)) licks spaced at 8 Hz from its onset, in all sessions. Anticipatory licks are the licks 0.5-1.5 s after odor onset.

#### Firing-rate matrices and drift test

For every unit, spikes were binned in 50-ms bins from 2 s before to 4 s after odor onset and smoothed with a Gaussian kernel (s.d. 100 ms). Only trials with the familiar odors were analyzed, and the two trials following a long inter-trial interval were excluded from all fits and trial averages (they are analyzed separately below). For the population analyses the rate of each unit was divided by the range of its condition-averaged responses (four conditions, odor × context, over the whole trial window) plus 5 Hz (soft normalization). Each unit was tested for drift by comparing its mean rate per trial (−1 to 4 s from odor onset) between blocks 1 and 3, which share a context: a unit was discarded as drifting when this difference was significant (two-tailed Welch’s t-test, *P*<0.05) and had the same sign as the difference between blocks 4 and 2. Population analyses used the sessions with at least 20 non-drifting units of the region (amygdala, 29 sessions from 7 mice; hippocampus, 33 sessions from 7 mice; basal ganglia, 21 sessions); unit counts before and after the drift test are given for the amygdala and hippocampal subregions in Extended Data Fig. 8c.

#### Single-unit coding

Per session, a Rescorla-Wagner value V was computed for each odor from the trial sequence, V ← V + α(r − V) after every presentation (r = 1 for reward, 0 otherwise; initial value 0.5), with α chosen for each session among 41 logarithmically spaced values between 0.01 and 1 to maximize the correlation between V and the anticipatory licks. For every unit, the trial-by-trial mean rate in the odor window (0-1.5 s) was regressed (ordinary least squares) on odor identity (±1), the centred Rescorla-Wagner value and context (±1), and the mean rate in the outcome window (1.5-4 s) on the same regressors plus reward (±1). A unit coded odor identity or value when the corresponding coefficient of the odor-window regression, and outcome when the reward coefficient of the outcome-window regression, was significant (t-test, *P*<0.01). Context coding was assessed on the non-drifting units only, by regressing the mean rate in the 2 s before odor onset, after the first 10 trials of every block, on context (*P*<0.01). A value unit was classified as excited by the CS+ when its value coefficient was positive. Fractions of coding units per subregion are shown with 95% Wilson confidence intervals (Extended Data Fig. 8f-j). Example units (Extended Data Fig. 8d, e) were chosen among the non-drifting units with a firing rate of at least 2 Hz and a coefficient significant at *P*<10⁻⁴ (context, *P*<10⁻³; outcome examples in addition without a CS+/CS− difference in the odor window, *P*>0.05), and are drawn as rasters and peri-stimulus time histograms (50-ms bins, Gaussian smoothing with s.d. 50 ms). For the example units, the value raster and PSTH and the context PSTH use the trials after the first 10 of each block (Extended Data Fig. 8d, e).

#### Population axes for odor identity, outcome and value

The odor-identity, value and outcome axes (Fig. 4c, e and Extended Data Fig. 8k) were obtained by dPCA (see Context axis and good-context sessions) in every session with at least 20 non-drifting units of the region, fitted on the condition-averaged rates (odor × context) of the trials after the first 10 of every block: odor identity, the first component of the odor marginalization fitted on the odor window (0-1.5 s); value, the first component of the odor × context marginalization fitted on the odor window; and outcome, the first component of the same marginalization fitted on the window after the outcome (1.5-4 s), made orthogonal to the odor-identity and value axes (Gram-Schmidt), because in this task the odor × context marginalization also carries the CS+/CS− difference during the odor. Every trial was projected on each axis, and the projection was normalized so that the two class means of the fitted trials in the axis’s window were at +1 and −1 (odor identity, 0-1.5 s, odor 1 as labelled in Fig. 4c at +1; value, 0.5-1.5 s, CS+ at +1; outcome, 1.5-4 s, reward at +1). Curves are trial averages per session (odor-identity and outcome axes, split by odor and CS+/CS−; value axis, split by CS+/CS− and by early trials, the first 10 trials after a reversal in blocks 2-4, versus late trials, the last 10 trials of each block), then mean ± s.e.m. over sessions. The context axis was fitted as described next.

#### Context axis and good-context sessions

The context axis was obtained by demixed principal component analysis (dPCA^85^; Python package dPCA 1.0.5) on the non-drifting units of a session, with the marginalizations condition-independent, odor, context (block type) and odor × context (equivalent to outcome, since an odor was rewarded if and only if it was the rewarded odor of the block). The regularization parameter was selected for each session and region by the cross-validation routine of the package and reused in all subsequent fits of that session. The context axis was the first component of the context marginalization, fitted on the condition-averaged rates (odor × context) of the trials after the first 10 of every block over −2 to 4 s from odor onset; the value axis used in Fig. 4d, f, h, i was the first component of the odor × context marginalization fitted on the odor window (0-1.5 s). Every trial (including the first 10 trials of each block) was projected on each axis, and the projections were normalized so that the two class means of the fitted trials in the axis’s own window were at +1 and −1 (context, 2 s before odor onset: context A, +1; context B, −1, as labelled in Fig. 4c; value, 0.5-1.5 s: CS+, +1; CS−, −1). Early and late trials are defined as above. A context axis was considered reliable, and the session included in the context analyses (good-context sessions; Fig. 4c-f, j, k and Extended Data Fig. 8c, k-m), when (i) the session had at least 5 non-drifting units whose rate differed between the contexts (two-tailed Welch’s t-test between the whole-trial rates, −1 to 4 s, of context A and context B trials after the first 10 trials of each block, *P*<0.05, with the three block-to-block changes of the block means alternating in sign); (ii) the mean context projection of the trials after the first 20 of each of blocks 2-4 lay on the side of its own context; and (iii) a held-out test of context coding exceeded a margin of 0.05. For the latter, a plain coding direction (difference between the mean pre-odor population vectors of the two contexts, after the first 10 trials of each block) was fitted on the even-numbered trials and used to classify the odd-numbered trials and vice versa, and the fraction of correctly classified trials was compared with the same procedure applied to block labels shifted by 0.3-0.7 of a block length (mean over shifts); the margin is the accuracy with the true labels minus the mean accuracy with the shifted labels. 14 of 29 amygdala sessions (5 mice) and 11 of 33 hippocampal sessions (7 mice) met all three criteria. The context curves of Fig. 4c, e are trial averages per session, split by context and by early versus late trials, then mean ± s.e.m. over the good-context sessions.

#### Value against context

For every trial of the good-context sessions, the context projection (mean over the 2 s before odor onset) and the value projection (mean over 0.5-1.5 s after odor onset) were computed as above. Trials were pooled over sessions (their projections are already in normalized units), binned by their context projection in bins of 0.5 between −1.75 and 1.75, and the mean ± s.e.m. of the value projection over the trials of each bin was drawn separately for odor 1 and odor 2 and as their difference, odor 1 minus odor 2 (Fig. 4d, f); bins with fewer than 15 trials were omitted (Fig. 4d, 3,176 trials from the 14 good-context amygdala sessions; Fig. 4f, 2,458 trials from the 11 good-context hippocampal sessions).

#### Decoding

Odor identity (odor 1 versus odor 2; 0-1.5 s), CS+ versus CS− (0.5-1.5 s), outcome (reward versus no reward; 1.5-4 s) and context (context A versus context B; the 2 s before odor onset) were decoded from the window-averaged rates of the non-drifting units of a session with L2-regularized logistic regression (regularization parameter C = 1, class weights balanced; scikit-learn) on z-scored rates. Decoders were trained on the trials after the first 10 of all four blocks with 5-fold cross-validation stratified by block, so that every block was represented in every training set; held-out late trials were scored by the model of their fold, and the early trials (the first 10 trials after each reversal, blocks 2-4) by the mean predicted probability of the five fold models. Accuracy is the balanced accuracy (chance 0.5), per session and per trial position in the block (Extended Data Fig. 8l, m). Odor identity, CS+/CS− and outcome were decoded in every session with at least 20 non-drifting units of the region and context in the good-context sessions.

#### Long inter-trial intervals

Licking around the long inter-trial interval (Fig. 4g) was quantified in all 64 sessions with long inter-trial intervals as the lick rate (all licks, 50-ms bins, Gaussian smoothing with s.d. 100 ms) on the trial after the long interval and on the last five trials with the same odor in the same block before it (averaged), with the bars giving the mean rate 0.5-1.5 s after odor onset; the trial after the long interval was compared with the trials before it with a two-tailed Wilcoxon signed-rank test over long intervals, separately for CS+ and CS− probe trials. The value projection around the long interval (Fig. 4h, i) was computed in the same way in every session with at least 20 non-drifting units of the region, using the dPCA value axis fitted on the normal trials of the session (the trial after the long interval was not part of the fit) and projecting the rates of the probe trial with the same unit scaling, mean and axis; the value projection is the mean over 0.5-1.5 s after odor onset in the normalized units defined above. The context projection during the long interval (Fig. 4j, k) was computed in the good-context sessions: spikes from the previous outcome to the odor onset were binned in 50-ms bins, smoothed (s.d. 100 ms), soft-normalized and projected on the context axis of the session, and the projection was divided by the block’s own level (the mean pre-odor context projection of the trials after the first 10 of that block, with its magnitude floored at 0.5), so that 1 is the level of the block on normal trials and 0 the midpoint between the contexts. Time was measured from the last outcome; the normal-trial time was the mean interval from the outcome to the centre of the 2-s pre-odor window on the normal trials of the block (7.2 s on average). Long intervals whose mean projection within ±2.5 s of the normal-trial time was not on the side of the current context (that is, in which the context was not decoded correctly at the time at which the next odor would normally have arrived) were excluded; the remaining traces (amygdala, 19 long intervals from 12 sessions; hippocampus, 16 from 10 sessions) were averaged in that ±2.5-s window and in 10-s bins thereafter, and the projection at the normal-trial time (first bin, ∼7 s) was compared with that at the end of the long interval (last bin, ∼85 s) with a two-tailed Wilcoxon signed-rank test over long intervals. Traces and bars in Fig. 4g-k are mean ± s.e.m. over long inter-trial intervals.

### RNN modelling

#### Recurrent neural network implementation

We implemented RNNs as previously described^28,29^. Briefly, we trained recurrent neural networks composed of GRU units (N=20) to estimate value at each timestep. The hidden unit activity is given by the following equation:

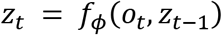

*z_t_* is hidden units’ activity at time t

*φ* is the parameter vector of the RNN

*o_t_* is the observation produced by environment at time t

*z_t_*_−1_ is the hidden units’ activity at time t-1

*o_t_* is a one-hot vector defined as follows:

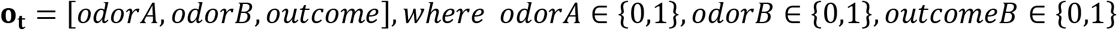

Thus, at each timestep, the RNN had access to a vector indicating whether odor A (cue 1) was present, odor B (cue 2) was present, and whether reward was delivered or not. Each trial began with an inter-trial interval (ITI) of 5 + k silent timesteps, with k drawn from a geometric distribution with parameter p = 0.8 (see Task implementation). After the ITI, a cue was presented (**o_t_** = [1,0,0] *or* **o_t_** = [0,1,0] for odor A or B respectively) followed by the outcome (**o_t_** = [0,0,1] *or* [0,0,0]] for reward or no reward respectively). V*_t_* = *w*^T^*z_t_* + *w*_0_ for *z_t_*, *w* ∈ ℝ*^H^* (where H=20 is the number of hidden units), *w*_0_, *V_t_* ∈ ℝ . The full parameter vector *θ* = [***φ***, ***w***, *w*_0_] was learned using TD learning. This involved backpropagating the gradient of the squared error loss *δ*^2^ = (*r_t_*_+1_ + *γV_t_*_+1_ − *V_t_*)^2^ with respect to *V_t_*. The discount factor *γ* was set to *γ*=0.2.

#### Task implementation

Each trial consisted of an ITI of silent timesteps (zero input), one timestep on which the cue was presented, and the outcome one timestep later (reward, 1, or no reward, 0). The ITI lasted 5 + k timesteps, with k drawn from a geometric distribution with parameter p=0.8 (P(k)=0.8×0.2^k; minimum ITI 5 timesteps, mean 5.25 timesteps), so that one trial lasted 7.25 timesteps on average. On every trial one of the two cues was chosen at random with equal probability. In the stable task, one cue was always rewarded and the other never. Depending on the analysis, the stable task consisted of 200 trials of this fixed contingency (Fig. 2h, j, o, Extended Data Fig. 4n), of 3 blocks of 50 trials (initial-acquisition traces of Fig. 2b, c and Extended Data Fig. 2d; Fig. 5f, h; Extended Data Fig. 9j, k) or of one 200-trial block trained from scratch (initial learning constants, see below); "after one block of the stable task" (Fig. 2q, r, Extended Data Fig. 4l) denotes a network at the end of the first 50-trial block of its training, when it had experienced only one contingency. In the dynamic task the reward contingency of the two cues was reversed at every block boundary, for 19 blocks of 50 trials; the contingency of the first block was chosen at random for each network, and no input signaled the block boundaries. The value at the cue is the value readout on the timestep on which the cue was presented, before the outcome.

#### Training

Each network was initialized with a different random seed: the recurrent weights of each GRU gate were orthogonal (gain 1), the input weights Glorot-uniform, the GRU biases and the output bias zero, and the readout weights were drawn from a uniform distribution between −1/√H and 1/√H (about ±0.22; the PyTorch default for a linear layer). Networks learned online, in a single pass over the trial stream, using truncated backpropagation through time (TBPTT)^86,87^ with a sliding window and stride 1: at every timestep t the network was run over the past W timesteps (from t−W to t; over all timesteps since the start of the stream while t<W), the loss was computed over that window and a single gradient step was taken, so that the weights changed at every timestep, as plasticity would during continuous learning. The hidden state at the start of each window was the state the network had reached at that timestep on the previous step, so that the state was carried through the stream rather than recomputed with the updated weights. The loss was the mean over the window of the squared TD error, (r_t+1_ + γV_t+1_ − V_t_)^2^, with the bootstrapped target r_t+1_ + γV_t+1_ treated as a constant (semi-gradient TD(0)), plus an L1 cost on the hidden-unit activity, λ×mean|z_t_| over the hidden states of the window, with λ=0.03. The activity cost penalizes activity that is not needed for the value prediction; it improved convergence on the dynamic task, and networks trained without it are compared in Extended Data Fig. 4a, f, g. Weights were updated with the Adam optimizer (learning rate 0.0005, AMSGrad variant, amsgrad=True in PyTorch, which enforces a non-decreasing second-moment estimate for more stable updates). For the majority of the analyses we fixed the window size to W=720 timesteps (∼100 trials, roughly the length of two blocks of the dynamic task); the effect of W is shown in Fig. 2b-d, i and Extended Data Figs 2-4, with W=45 as the short window of Fig. 2 and Extended Data Figs 2, 4h-n and 5. Given that the network is continuously learning in this setting, it is possible that learning gets stuck in non-optimal local minima. To exclude networks that had not learned the task, we computed the mean squared reward prediction error (RPE^2^) over the last 20 trials of each of four late blocks of the dynamic task (blocks 16-19, or blocks 14-17 in the runs that recorded the training traces, the plasticity-blocking clones and the ITI-silencing probes) and defined a network as converged when this value was below 0.005 (Extended Data Fig. 2a). Unless stated otherwise, only converged networks were analysed; the networks with a frozen weight compartment (Fig. 5e-h, Extended Data Fig. 9i-m) were all kept. The fixed-contingency runs that give the initial-acquisition traces (Fig. 2b, c, Extended Data Fig. 2d) and the networks of the weight-update analyses (Extended Data Fig. 5) were not selected either. The number of networks behind each panel is given in the legends.

#### Learning constants for RNNs

The value at the cue was recorded on every trial of the training pass with the weights as they were at that moment. Initial acquisition (Fig. 2b, c; Extended Data Fig. 2d, initial acquisition) was taken from networks trained with the standard recipe on three blocks of 50 trials of one fixed contingency (cue 2 rewarded throughout; the control networks of the frozen-compartment experiment, trained at every window; 50 networks per window, no convergence criterion): the value at the cue on the CS+ trials over the first 50 trials (Fig. 2b, c) or on the CS+ and CS- trials over the first 100 trials (Extended Data Fig. 2d). The first reversal (Extended Data Fig. 2d, 1st reversal) was the reversal from block 1 to block 2 of the serial-reversal training, and the expert reversals (Fig. 2b, c; Extended Data Fig. 2d, expert reversal) were the reversals into blocks 11–19 of the same runs, averaged per network; the first 50 converged networks of the 60 trained at each window were used (see Window sweep), and trial 0 is the first trial of the new block. As for the mice, we fitted an exponential, y(t) = a ∓ b·exp(−t/τ), to the value of one cue as a function of the number of presentations of that cue (t = 0, 1, 2, …, counting only the trials on which that cue was shown), with a rising fit for acquisition of the CS+ or the new CS+ and a falling fit for extinction of the old CS+ (the sign of b was fixed by the expected direction of the change). τ was found on a grid of 200 values spaced logarithmically between 0.5 and 500 presentations, with a and b solved in closed form for each τ; a time constant of 0.5 presentations (the lower limit of the grid) means the value switched within a single presentation and is reported as an upper bound (<1). Because the 50-trial blocks of the dynamic task are too short to fit the slow constants of naive networks, the initial and first-reversal constants were estimated on 200-trial blocks of fresh networks trained with the standard recipe at each window: for the initial constant, a fresh network trained online on 200 trials of one fixed contingency (both cues shown); for the first reversal, a fresh network trained on 50 trials of one contingency followed by 200 trials of the reversed contingency, which gave the acquisition constant of the new CS+ and the extinction constant of the old CS+ (n=50 networks per window; Fig. 2b, c; Extended Data Fig. 2d, learning constant). The expert constants were fitted per network on the training reversals themselves, as for the mice (Fig. 2b, c; Extended Data Fig. 2d, learning constant): the value of the new CS+ (or the old CS+) after each of the reversals into blocks 11–19 was taken by presentation of that cue, the reversals were averaged by presentation index and the first 25 presentations were fitted. Time constants are reported as the median over networks with a 95% percentile bootstrap confidence interval of the median (2,000 resamples). The value traces of Fig. 2b, c are plotted against the trial of the block (both cues counted, trial 0 being the first trial of training or of the new block), and each trial is the mean over the networks that were shown the CS+ on that trial.

#### Plasticity blocking

To test what the network could do without any weight change (Fig. 2e, f, i; Extended Data Figs 3b, c and 4m), the network was cloned at the start of a block: the clone’s weights were frozen for the entire block (no gradient step), and it was run over the same trial sequence from the same hidden state as the plastic network, so that the two differed only in whether the weights could change. The training pass of the plastic network itself was the control and was never perturbed. The value at the cue was recorded for both networks on every trial of the block. The clone was made at the start of every block (Fig. 2i; Extended Data Figs 3b, c and 4m; the first 50 converged networks of each window in Fig. 2i and Extended Data Figs 3 and 4m) or at the start of blocks 1, 2, 3 and 19 (Fig. 2f, initial, block 1, and expert, block 19; the first 50 converged networks). To quantify cue discrimination we computed, per network, the area under the receiver operating characteristic curve (AUROC) between the values of the CS+ and the CS- trials over the last 25 trials of the block (the probability that the value on a randomly chosen CS+ trial exceeds that on a randomly chosen CS- trial, ties counted as one half), and the value difference V(CS+) − V(CS-) over the same trials (Extended Data Figs 3b and 4m; for the AUROC of Extended Data Fig. 3b, see below). The effect of freezing was defined per network as control minus frozen and compared between block 1 (initial acquisition) and block 19 (expert reversal) with a two-tailed Wilcoxon signed-rank test (Extended Data Fig. 4m), since the same networks contributed to both blocks. In Extended Data Fig. 3b, as for the ITI-silencing probes of Extended Data Fig. 3a, the AUROC was computed across networks at each block: the mean CS+ values of the networks over the last 25 trials of the block were ranked against their mean CS- values, with a bootstrap s.e.m. over networks (500 resamples).

#### ITI silencing

To test whether the value depended on the activity carried through the ITI, all hidden units were set to zero (silenced) for one timestep of the ITI. For the traces of Fig. 2h, networks with frozen weights (initial: after 200 trials of one contingency; expert: end of block 19 of the dynamic task; the first 50 converged networks of each set) were first settled for 20 trials in one context and then run for 10 recorded trials before and 10 after the silenced trial. On the silenced trial all units were set to zero at the penultimate timestep of the ITI, so that the state evolved freely for one timestep before the cue; the silenced state was carried on, so that the following trials show the recovery. At every trial the values of both cues were read by forking the episode (the same state was presented with cue A or with cue B), so that a CS+ and a CS- value exist on every trial. The control was the identical trial sequence without silencing. Each network was probed 4 times per context, and the two contexts of the dynamic task were averaged so that any cue bias cancelled. For the AUROC of Extended Data Fig. 4n, networks of the same two kinds (stable expert, 200 trials of one contingency, and dynamic expert, block 19; n=50 networks each at W=720) were settled for 20 trials in a context and then given 40 probe trials per context; on each probe trial a copy of the state ran that trial’s ITI with all units set to zero at the first ITI timestep (immediately after the previous outcome), the remaining ITI timesteps (at least 4) ran freely, and the trial’s own cue was presented; the control was the same state and trial without silencing, and the trajectory carrying on from trial to trial was never silenced. The networks trained with W=45 (Extended Data Fig. 4n, top row) were probed only in the context of their own last training block (see Long ITIs in the RNNs). The AUROC between CS+ and CS- values was computed per network over its probe trials, and the effect of silencing (control minus silenced) was compared between stable and dynamic experts with a two-tailed Mann-Whitney U test, since different networks contributed to the two stages. To follow the effect of ITI silencing over training (Fig. 2i; Extended Data Fig. 3a), the same probe was applied during learning as a read-only clone: on two trials per block (one CS+ and one CS-, drawn from the last 10 trials of the block and at least 5 trials apart), the hidden state at the start of the trial was copied and, in the copy only, all units were set to zero at the first ITI timestep; the rest of the ITI ran freely, the value at the cue was read from the copy, and a copy of the same trial without silencing served as control, so that training was never perturbed. These probes were run at every window (the first 50 converged networks of each window; W=720 and W=45 in Fig. 2i). At each block, the CS+ probe values of all networks were ranked against their CS- probe values to give one AUROC per block, with a bootstrap s.e.m. over networks (500 resamples; Extended Data Fig. 3a).

### Discrimination loss

The contribution of each mechanism to cue discrimination (Fig. 2i, Extended Data Fig. 3c) was defined per network and block as d_control − d_manipulation, where d = V(CS+) − V(CS-). For plasticity, d_control was computed over the last 25 trials of the block in the plastic network and d_manipulation over the same trials in its clone with frozen weights (see Plasticity blocking). For ITI activity, d_control and d_manipulation were computed over the intact and the silenced read-only copies of the two probe trials of the block (see ITI silencing). Fig. 2i shows blocks 1-10 of training for the long (W=720) and the short (W=45) window, each on its own axis (the first 50 converged networks of each window and manipulation), and Extended Data Fig. 3c all blocks for every window. In both, consecutive blocks were averaged in pairs, so that the two block types (each cue rewarded) contribute equally and a preference for one cue cancels (the silenced readout of the clone favours one cue and would otherwise alternate in sign with the contingency of the block). A loss above 1 means that the frozen clone kept the contingency of the previous block.

#### Frozen weight compartments

As a model of a compartment-specific block of plasticity (Fig. 5e-h, Extended Data Fig. 9i-m), networks were trained with the standard recipe while one set of weights was held at its random initial value for the whole run by excluding it from the optimizer: the recurrent weights and their biases (RNN^Δw=0), the input weights and their biases, or the readout weights and the output bias. Control networks had all weights plastic. Separate sets of networks were trained on the stable task (3 blocks of 50 trials of one fixed contingency) and on the dynamic task (the standard 19-block run), 50 networks per condition and task; conditions were compared with a two-tailed Mann-Whitney U test versus control. Discrimination was quantified as the AUROC between the values at the cue on CS+ and CS- trials over all trials of a block (stable task: the mean over the 3 blocks; dynamic task: block 19), and the value at the cue was plotted by presentation of each cue (stable: the first 25 presentations of each cue in each block; dynamic: the last 10 presentations before and the first 25 after the last reversal). The control networks of the stable task were also trained at the other TBPTT windows (50 networks per window) and provided the initial-acquisition traces of Fig. 2b, c and Extended Data Fig. 2d. For Extended Data Fig. 5k, l, control and recurrent-frozen networks were also trained on the dynamic task with W=45 (50 networks per condition; at W=720, the dynamic-task networks of Fig. 5e-h were used). The learning constant was fitted as described in Learning constants for RNNs, to the value at the cue of the CS+ over its first 25 presentations in block 1 (initial acquisition) and of the new CS+ over its first 25 presentations after each reversal; the expert constant of a network is the median over the reversals into blocks 16-19. Because the blocks of these runs are 50 trials long, these initial constants are not comparable with the 200-trial estimates of Fig. 2 and Extended Data Fig. 2. The value traces show the last 10 presentations before and the first 25 after the first reversal and the expert reversals (averaged per network). Initial and expert constants were compared within a condition with a two-tailed Wilcoxon signed-rank test, and between conditions with a two-tailed Mann-Whitney U test.

#### Window sweep

To vary how far back in time the gradient could reach, networks were trained with W = 20, 45, 90, 180, 360 or 720 timesteps, everything else unchanged. For each window 100 networks were trained on the dynamic task (Extended Data Fig. 2a-c, all networks shown). Separate runs of the dynamic task at each window recorded the plasticity-blocking clones and the ITI-silencing probes; for these manipulations (Fig. 2i, Extended Data Fig. 3) and for the learning curves of Extended Data Fig. 2e the first 50 converged networks of each window were analysed. For the learning constants and the meta-learning speed-up (Fig. 2b-d; Extended Data Fig. 2d), the initial-acquisition traces came from 50 networks per window trained on one fixed contingency, the initial and first-reversal constants from 50 fresh networks per window trained on the 200-trial blocks described above, and the first-reversal and expert traces and the expert constants from the first 50 converged networks of the separate runs of that window (see Learning constants for RNNs). W = 45 is the short window of Fig. 2 (c, i) and Extended Data Fig. 2 (d shows W = 720, 180 and 45); it is also compared with W = 720 in Extended Data Figs 4h-n and 5.

#### Weight-update size

To measure how much the weights changed over training (Extended Data Fig. 4d), the size of every gradient step, |Δθ|, was recorded as the Euclidean norm of the change of the parameters produced by one optimizer step, for all parameters together and separately for the recurrent weights, the input weights, the GRU biases and the readout weights (including the output bias), and averaged over the steps of each block (the first 100 converged of 106 networks).

#### Meta-learning speed-up in RNNs

The speed-up of learning with training (Fig. 2d) was defined for each window as the median initial learning constant divided by the median expert learning constant of the new CS+ (n=50 networks per window for each constant: the first 50 fresh networks of the 200-trial blocks and the first 50 converged networks of the separate runs of the window sweep). Its 95% confidence interval was obtained by percentile bootstrap (20,000 resamples), resampling the initial and the expert constants independently, as they come from different networks. All data are shown as mean ± s.e.m. over networks unless stated otherwise.

### RNN state-space analyses

#### State-space analysis of the RNNs

To visualize the hidden-state dynamics of a trained network, we froze its weights at the checkpoint of interest and ran it on a probe stream of the task (50 trials per block, cues chosen at random, ITI of 5 + geometric(0.8) − 1 steps as in training), taking the hidden state at the last ITI step before each cue as the ITI state of that trial. Principal component analysis (PCA) was fitted on all ITI states of the probe block or blocks shown in each panel, and the ITI states of the last 5 trials (stable expert) or of the last 10 trials of each context (dynamic expert) were plotted in the PC1–PC2 plane (Fig. 2j; Extended Data Fig. 4c). The autonomous flow field was obtained by embedding every point of a 24 × 24 grid of the PC plane back into the 20-dimensional state space (remaining components at their mean), stepping it once through the GRU with zero input and the weights frozen, and projecting the displacement back onto the plane, drawn as streamlines. Silent-ITI trajectories (Fig. 2o) were produced by the same zero-input map, started at the last plotted ITI state of a context and iterated for 40 steps, with a marker every 5 steps so that the spacing of the markers indicates the speed of the state.

#### Fixed points and slow points

Fixed points of the autonomous dynamics x = F(x), where F is one GRU step under zero input, were found by minimizing the kinetic energy q(x) = ½‖x − F(x)‖²^88^. Because every unit of a GRU is confined to (−1, 1), we optimized over u with x = tanh(u), which prevents the spurious large-norm solutions that an unconstrained search produces for this architecture. Starting points were noisy copies (Gaussian noise, s.d. 0.05) of the ITI states of the probe stream: 600 starts for Fig. 2j and Extended Data Fig. 4b,c, and 1,500 starts plus 400 points sampled along the segment between the two context means (position t drawn uniformly from −0.3 to 1.3, same noise) for the classification in Extended Data Fig. 4a. All starts were optimized together with Adam (learning rate 0.05, reduced by 10% whenever the mean q over the starts had not decreased for 20 iterations) for 3,000 iterations, and a start was counted as converged when q < 10⁻⁶. Converged points were merged greedily, keeping the slowest of any two points closer than a merge radius: fixed points at a radius of 0.3 (Fig. 2j; Extended Data Fig. 4a,c) or 0.03 (Extended Data Fig. 4b), and slow points always at 0.03, one within-context s.d. of the ITI states (0.032). A converged point was called a fixed point when q < 10⁻¹⁰ (Extended Data Fig. 4a, b) or q < 10⁻⁸ (Fig. 2j; Extended Data Fig. 4c) and a slow point when q lay between that bound and 10⁻⁶. Stability was read from the eigenvalues of the Jacobian of F at the fixed point, obtained by automatic differentiation: a fixed point was stable when every eigenvalue λ of the Jacobian satisfied |λ| < 1 and a saddle or unstable point otherwise. Because the slow points lie on a ridge rather than being isolated, their number is a sampling density that depends on the merge radius (2.8 per network at 0.03 against 0.8 at 0.3); the number of stable fixed points does not (1.1 per network at either radius).

#### Attractor classification and training rules compared

Extended Data Fig. 4a compares five training rules on the dynamic task (20 hidden units, deterministic contingencies, blocks of 50 trials, ITI 5 + geometric(0.8) − 1 steps): full BPTT with a tanh RNN or a GRU, trained on episodes of three blocks of 50 ± 2 trials (200 episodes per epoch, 400 epochs, Adam, batch size 16, learning rate 3 × 10⁻³); online TBPTT (W = 720, one gradient step per timestep, Adam with AMSGrad, learning rate 5 × 10⁻⁴) with a tanh RNN or a GRU on a single episode of 19 blocks without an activity cost, in which the first weight update was taken once the window was full (after 720 timesteps); and a separate set of online TBPTT GRUs trained with the recipe of all other figures (L1 activity cost, λ = 0.03; updates from the first timestep). For each rule the first 50 networks that passed the convergence gate were analysed; for the online rules the gate was the mean RPE² over the last 20 trials of four late blocks of the training pass (< 0.005; see Training), and for the full-BPTT networks the same windows of a six-block probe episode run with the weights frozen (all 55 networks of each full-BPTT cell passed, median RPE² 0.0018). For each network, a context axis was defined on a frozen-weight probe of 60 three-block episodes as the difference between the mean ITI states of the two block types, m₁ − m₀, and every converged point was placed on it as proj(x) = (x − m₀)·(m₁ − m₀)/‖m₁ − m₀‖² (0, the context in which cue 1 is rewarded; 1, the context in which cue 2 is rewarded). Networks were then classified by three criteria: (1) a marginal mode aligned with the context axis, i.e. a converged point (fixed or slow) whose basin attracted at least 5% of the 1,900 starts, whose Jacobian eigenvalue closest to 1 had |λ| ≥ 0.98 and whose eigenvector had |cos| ≥ 0.7 with the unit context axis; (2) a flat step-size profile, i.e. the step size ‖F(x) − x‖ of the zero-input map, sampled at 300 points along the segment from t = −0.3 to 1.3, varied by less than a factor of 2 over the interior of the segment (0 ≤ t ≤ 1); and (3) interior points, i.e. at least one additional converged point resolved at the fine merge radius (0.03) that was more than 0.3 from every point found at the coarse radius and projected between 0.1 and 0.9 on the context axis. A network meeting all three criteria was called a line attractor, a network meeting none discrete attractors, and any other network mixed. The number of stable fixed points per network (Extended Data Fig. 4a, middle) was counted at the coarse radius with the bound q < 10⁻¹⁰, and the number of slow points (right) was the number of converged points at the fine radius that were not fixed points.

#### Context axis of the RNNs

The context axis was always the direction between the mean ITI states of the two contexts; its estimation and scaling differed with the panel. For the projections around the cue (Fig. 2l; Extended Data Fig. 4i), the expert network with frozen weights was run on four alternating blocks of 50 trials (contexts A, B, A, B). The axis was the difference between the mean pre-cue states (last ITI step) of the late trials (last 25 of each block) of the two contexts, estimated on a random half of the late trials; the hidden state at every timestep was projected onto it after subtracting the midpoint of the two means and scaled so that the two means lay at +1 (context A, cue 1 rewarded) and −1 (context B, cue 2 rewarded). Only the other half of the late trials and the early trials (first 5 of each block) were plotted, so the late curves did not enter their own axis. For the positions of the fixed points (Extended Data Fig. 4b) and the value maps (Fig. 2m, n; Extended Data Fig. 4j,k), the probe stream comprised the two contexts of blocks 18 and 19 (50 trials each), the context means m₀ and m₁ were the mean ITI states of the last 10 trials of each context, and states were placed as proj(x) = (x − m₀)·(m₁ − m₀)/‖m₁ − m₀‖², with 0 at the mean of the context in which cue 1 is rewarded and 1 at that in which cue 2 is rewarded; for display the axis was rescaled to run from −1 to 1 as 1 − 2·proj(x), so that context A lies at +1, as in Fig. 2l (Fig. 2m, n; Extended Data Fig. 4b, j, k); the axis was oriented by cue identity, not by block order, so that networks could be pooled. For the projections during a silent ITI (Fig. 2p; Extended Data Fig. 4g), the state was projected onto the unit vector from m₀ to m₁ relative to the midpoint and divided by the level of the context in which the network had been settled, (m_c − mid)·u, so that +1 is the level of the context just experienced, −1 the level of the other context and 0 the midpoint; the trace of a network is the mean over its two settle contexts, so that a network that rests at one context whatever it experienced scores 0 rather than +1 and a standing cue preference cannot appear as retention.

#### Context decoding across blocks

To follow the emergence of a context code during training (Fig. 2k; Extended Data Fig. 4h), we used the hidden state at the last ITI step of every trial of the training pass itself (online states, with the weights still updating). An L2-regularized logistic-regression decoder (unit activities standardized on the training set; C = 1) was trained to separate all trials of blocks k and k + 1, which have opposite contingencies, and tested on all trials of block k + 2, which repeats the contingency of block k; accuracy is the fraction of its trials labelled as block k, plotted against the block pair. Because the blocks alternate, the memory of the previous trial’s cue and outcome, which by itself identifies the block within a pair, appears with the opposite sign in block k + 2 and cannot support this generalization; only a context code that is reproduced from block to block does, so this curve starts at chance in naive networks and rises with training.

#### Value reported from each ITI state

To relate the position of the state on the context axis to the value the network would report (Fig. 2m, n; Extended Data Fig. 4j,k), the expert network with frozen weights was stepped once from every ITI state of the probe stream (100 trials) with the input of cue 1 and, from the same state, once with the input of cue 2 (no reward), and the value output at the cue step was read in each case. This yields for every state its context projection and the two values, V(cue 1) and V(cue 2), the network would report, the counterfactual the task never shows. Trials were pooled over networks and binned by context projection (12 bins of proj(x) from −0.1 to 1.1; bins with fewer than six trials not shown), and the value projection was defined as V(cue 1) − V(cue 2), the value of the cue rewarded in context A minus that of the other cue (Fig. 2n; Extended Data Fig. 4k).

### Long ITIs in the RNNs

To simulate the long-ITI experiment (Fig. 2q,r; Extended Data Fig. 4e,l), the network with frozen weights was settled for 30 trials in one context (random cues, normal ITIs), after which a pause of 5, 10, 20 or 40 silent steps (zero input) preceded a probe trial presenting the CS− of that context, as in the mouse experiment, followed by 10 normal trials. At every trial from 10 before to 9 after the pause, the value of both cues was read by forking the episode at that trial (same state before the cue, cue 1 or cue 2 presented), so that both value curves exist at every position. The discrimination index on the probe trial was (V(CS+) − V(CS−))/(|V(CS+)| + |V(CS−)| + 0.05) and was averaged per network over the two settle contexts and two repeats per context. Each network was classified, after averaging its two contexts, as kept (V(CS+) > 0.7 and V(CS−) < 0.3), flipped (V(CS+) < 0.3 and V(CS−) > 0.7) or halfway (Extended Data Fig. 4e); averaging the contexts first ensures that a network with a resting preference for one cue, which keeps the contingency in one context and flips it in the other, is counted as halfway rather than as retention. Stable-task networks were settled and probed in their single contingency. For the networks trained with W = 45 (Extended Data Fig. 4l,n), each dynamic network was settled only in the context of its own last training block. For the comparison of training rules (Extended Data Fig. 4f,g), each network was settled for 40 trials in one context (cues drawn at random) and the value of each cue was read at the cue of a probe trial after a silent ITI of 5, 10, 20 or 40 steps, using an identical settle sequence for the two cues (20 repeats per ITI and context); V(CS+) − V(CS−), with CS+ the cue rewarded in the settle context, was averaged over both settle contexts. The context projection during the silent ITI (Extended Data Fig. 4g) started from the state at the last step of the 40th settle trial, after its outcome, and followed the zero-input dynamics for 40 steps (5 repeats), normalized as described above with context means taken from the 60-episode probe of the fixed-point analysis. Fig. 2p used the same normalization on networks trained with the recipe of the main figures for 20 blocks (weights at the end of block 20): each network was settled with frozen weights for 40 trials in each context, the context means were the last 10 ITI states of each settle block, the trajectory started from the ITI state of the 40th settle trial and ran for 40 silent steps, and the projection after 5 silent steps, the minimum ITI of training, was compared with that after 40 steps in the same networks. In the example traces of Fig. 2q, trial 0 is the probe trial, the first trial after the pause.

#### Stable and dynamic expert networks

Unless stated otherwise, a dynamic expert is a network whose weights were taken at the end of the last block of the dynamic task (block 19; Fig. 2f,h,j,l-o,q,r; Extended Data Fig. 4b,c,e,i-n; block 20 of a 20-block run in Fig. 2p), and the last block itself served as the expert reversal. The expert reversals of Fig. 2b, c and Extended Data Fig. 2d are instead the reversals into blocks 11-19 of the training pass, with the weights as they were at that moment (see Learning constants for RNNs). Stable experts were obtained by training networks with the same initial weights (same seeds) and the same online recipe on a single contingency (cue 2 rewarded, cue 1 not) and taking the weights after 200 trials, i.e. four blocks of 50 trials (Fig. 2h,j,o; Extended Data Fig. 4n), or after the first block of 50 trials where the comparison was with initial acquisition (Fig. 2q,r; Extended Data Fig. 4l). Stable networks were always the first 50, in seed order, whose RPE² over the last 20 trials of each of their four blocks was below 0.005. Initial acquisition (block 1 of the dynamic task; Fig. 2f,i; Extended Data Fig. 3) is likewise the first 50 trials under one contingency; the initial-acquisition traces of Fig. 2b, c and Extended Data Fig. 2d come from the fixed-contingency runs of the frozen-compartment control (see Learning constants for RNNs), and the stable task of Fig. 5f comprised three blocks of 50 trials of one contingency.

#### Controlled networks for the weight-update analyses

The analyses of Extended Data Fig. 5d-j used a separate set of networks trained with the standard recipe (GRU, 20 hidden units, L1 activity cost λ = 0.03, Adam with AMSGrad, learning rate 5 × 10⁻⁴, discount factor 0.2, stride 1; see Training) at W = 45 and W = 720 on the dynamic task (19 blocks of 50 trials), n = 10 networks (seeds 0-9) per window. They differed from the other runs in two respects. First, the trial stream was generated from a seeded generator, so that a run could be reproduced exactly and every experiment below could be branched from it, and the first block rewarded cue 1 in every network, so that the reversals into the even-numbered blocks (2, 4, …, 18) were reversals from the context in which cue 1 is rewarded (context A) to that in which cue 2 is rewarded (context B). We call the reversal into block 2 the first reversal, that into block 4 early and those into blocks 16 and 18 late; late measures were averaged over the two blocks per network. Second, at the start of every block a checkpoint was saved (the weights, the state of the optimizer and the hidden state carried into the block), and the change of every parameter produced by every optimizer step was logged, so that the movement of the weights could be followed within a block; restoring a checkpoint and continuing the training reproduced the values of the training pass exactly. No convergence criterion was applied to these networks. Throughout Extended Data Fig. 5, d = V(new CS+) − V(old CS+) at the cue, i.e. V(CS+) − V(CS−) of the current block, with the value of each cue carried forward from its last presentation (undefined until both cues have been shown in the block), and the mean d of a block, the mean over its trials, is the area under the learning curve of the reversal (−1, never reversed; +1, reversed from the first trial). The parameters were grouped into the recurrent weights (1,200), the input weights (180), the readout weights (20) and the biases (121: the six GRU bias vectors and the readout bias), 1,521 parameters in all. Unless stated otherwise, conditions were compared within networks with the two-tailed Wilcoxon signed-rank test over the 10 networks, uncorrected.

#### Weight-space trajectories

For Extended Data Fig. 5a-c, networks were trained with the standard recipe on the dynamic task while the weight vector θ (1,521 parameters) was recorded at the end of every block: 10 networks per window for the direction and size of the weight change (Extended Data Fig. 5c), and one further network per window, from the same random seed at both windows, whose block-end weight vectors were saved for the example trajectories (Extended Data Fig. 5a, b). No convergence criterion was applied to these networks. The weight change of block k was dw_k = θ_k − θ_k−1_, where θ_k is the weight vector at the end of block k and θ_0 the initial weights. For the trajectory of the example network (Extended Data Fig. 5a, b), the block-end weight vectors, after subtraction of their mean, were projected onto two orthogonal unit vectors: the net-drift axis, the direction from θ_0 to θ_19, and the late-dw axis, the first principal direction (leading right-singular vector) of the weight changes of blocks 10-19 after removal of their component along the drift axis. The number printed with each axis is the share of the summed squared late weight changes, Σ ^19^ |dw_k|², that lies along that axis. The direction of the weight change (Extended Data Fig. 5c, top) was the cosine between the weight changes of consecutive blocks, cos(dw_k, dw_k−1_), for k = 2-19 (−1, the weights return along the previous block’s change; +1, they continue in the same direction), and its size (Extended Data Fig. 5c, bottom) the norm |dw_k|; both were averaged over the 10 networks of each window.

#### Where the reversal happens: mapping versus resting state

To ask whether a reversal changes where the network rests before the cue or what that state is read out as (Extended Data Fig. 5d), we compared, for each reversal block of the controlled networks, the weights at its start (θ_a, from the checkpoint) and at its end (θ_b). The resting state s of a network was its mean pre-cue state (last ITI step) over 30 probe trials of the contingency it had last been trained on, run with the weights frozen after 10 burn-in trials, starting from the hidden state the network had reached online at the block boundary. We used the pre-cue state rather than the fixed point of the autonomous dynamics (see Fixed points and slow points), because at W = 45 the ITI (5-10 steps) is shorter than the relaxation to the fixed point. The value the network reports when a cue arrives from a state was read after one GRU step with that cue’s input, for each cue from the same state (as in Value reported from each ITI state), and M(θ, s) denotes V(new CS+) − V(old CS+) obtained with weights θ from state s. The change across the block, M(θ_b, s_b) − M(θ_a, s_a), was split into three additive parts: the change of the mapping, ½[M(θ_b, s_a) − M(θ_a, s_a) + M(θ_b, s_b) − M(θ_a, s_b)], the same state read out with the new weights; the state moved by the outcomes, ½[M(θ_a, s_a′) − M(θ_a, s_a) + M(θ_b, s_a′) − M(θ_b, s_a)], where s_a′ is the resting state of the old network (θ_a, weights frozen) probed in the new contingency; and the state moved by the weights, ½[M(θ_a, s_b) − M(θ_a, s_a′) + M(θ_b, s_b) − M(θ_b, s_a′)], the remainder of the move of the resting state. This is a Shapley decomposition over state and weights in which the state path is split at s_a′. The parts were computed for the first reversal and for the two late reversals of each network, the latter averaged per network, and then averaged over networks.

### Value gained per unit of weight movement

For Extended Data Fig. 5e, the movement of the weights within a reversal block was followed from the logged optimizer steps of the controlled W = 45 networks: the cumulative movement before trial t, S(t), was the sum of |Δθ| (the Euclidean norm of the parameter change produced by one optimizer step; see Weight-update size) over all steps of the block taken before the value of trial t’s cue was read. Per network, d was expressed as a function of S by carrying its last value forward onto a common grid of S (0 to 8 in steps of 0.133), averaged over the blocks of a stage and then over networks; a curve is drawn from the first grid point at which at least half of the networks have a defined d to the largest block-end S among the networks of that stage. Whole-block measures were the path, S at the end of the block; the gain in d, the mean d over the last 10 trials of the block minus the first defined d; and the efficiency, the gain divided by the path. Stages were compared per measure with the paired Wilcoxon signed-rank test over networks.

#### One parameter group plastic

To ask which parameters can carry a reversal on the trained circuit (Extended Data Fig. 5f, g), training was continued from the checkpoint at the start of block 16 of each controlled W = 45 network (weights, optimizer state and carried hidden state restored, so that the continuation with all parameters plastic reproduced the training pass) with one group of parameters plastic and every other parameter frozen: after every optimizer step the parameters outside the plastic group were returned to their values before the step, so that they never changed although they still entered the gradient. The plastic group was all parameters, the recurrent weights, the input weights, the readout weights, all biases or none (Extended Data Fig. 5f), or one type of GRU bias (Extended Data Fig. 5g): the reset-gate biases (b_ir_ and b_hr_, 40 parameters), the update-gate biases (b_iz_ and b_hz_, 40), the input-side candidate biases (b_in_, 20) or the hidden-side candidate biases (b_hn_, 20, inside the reset-gated term of the candidate activation), in the notation of the GRU equations above. The outcome was the mean d of block 16, the first block of the continuation, compared with the continuation with all parameters plastic. The same continuation from the checkpoint at the start of block 2 (the first reversal) served as a control (not shown).

#### Matched bias change along the switch axis

To compare how much value the same bias change buys at the first and at a late reversal (Extended Data Fig. 5h), a switch axis u was estimated for each controlled W = 45 network as the mean of the per-block weight change over the blocks in which cue 2 was rewarded minus the mean over the blocks in which cue 1 was rewarded, using blocks 4-15 only, so that the blocks tested were excluded; +u points from the weights that solve context A toward those that solve context B. u_b is u restricted to the 121 biases and normalized to unit length. The weights at the start of the first reversal (block 2; first) and at the start of the late reversals (blocks 16 and 18, averaged per network; late) were perturbed to θ + εu_b with ε = 0, 0.1, 0.2, 0.3, 0.45, 0.6, 0.8 and 1 in parameter units (ε = 1.2 was also computed but is not shown; for scale, the net change of the biases over one reversal block is about 0.9 at the first and 0.55 at a late reversal), all weights were frozen, and d was read at the resting state on the old contingency as in the mapping analysis above (10 burn-in and 30 measured probe trials from the hidden state at the block boundary, the same probe trials at every ε). Δd is d(ε) − d(0). As a control for the sensitivity of the value to any bias change of that size, 10 random unit directions in the bias subspace were applied per network and stage in the same way, and their mean |Δd| is reported (the signed mean is zero by symmetry). The two stages were compared at ε = 0.6 with the paired Wilcoxon signed-rank test over networks.

#### Keeping or resetting one parameter group

To ask where the learned reversal speed is stored (Extended Data Fig. 5i, j), we took, for each controlled network, the weights at the start of block 18 (the block before the last reversal; the entry block) and the network’s initial weights θ_0, and built a hybrid weight vector in which either one group kept its trained values and every other parameter was reset to θ_0 (kept trained), or one group was reset to θ_0 and every other parameter kept its trained values (reset to initial); all parameters trained (all) and all reset (none) were the references. The hybrid network was then trained online with the trainer’s rule (the same window W, TD loss with the activity cost, Adam with AMSGrad, learning rate 5 × 10⁻⁴) from a fresh optimizer state, with the hidden state started from zero at the first window (which, as in training, reached back into the preceding block), through block 18, in which the old contingency could be re-learned or re-expressed, and block 19, the last reversal, and scored by the mean d over the first 25 trials of block 19 (trials on which d was undefined skipped), the 50-trial mean diluting the early difference between conditions. A kept group was compared with none and a reset group with all (paired Wilcoxon signed-rank test over networks). A condition whose mean d over the last 10 trials of the entry block, averaged over networks, was below 0.5 had not learned the old contingency, so that block 19 was an acquisition rather than a reversal for it; these conditions are drawn as open bars and were not tested (at W = 720, none and the input, readout and bias groups kept).

---

**Extended Data Fig 1.**
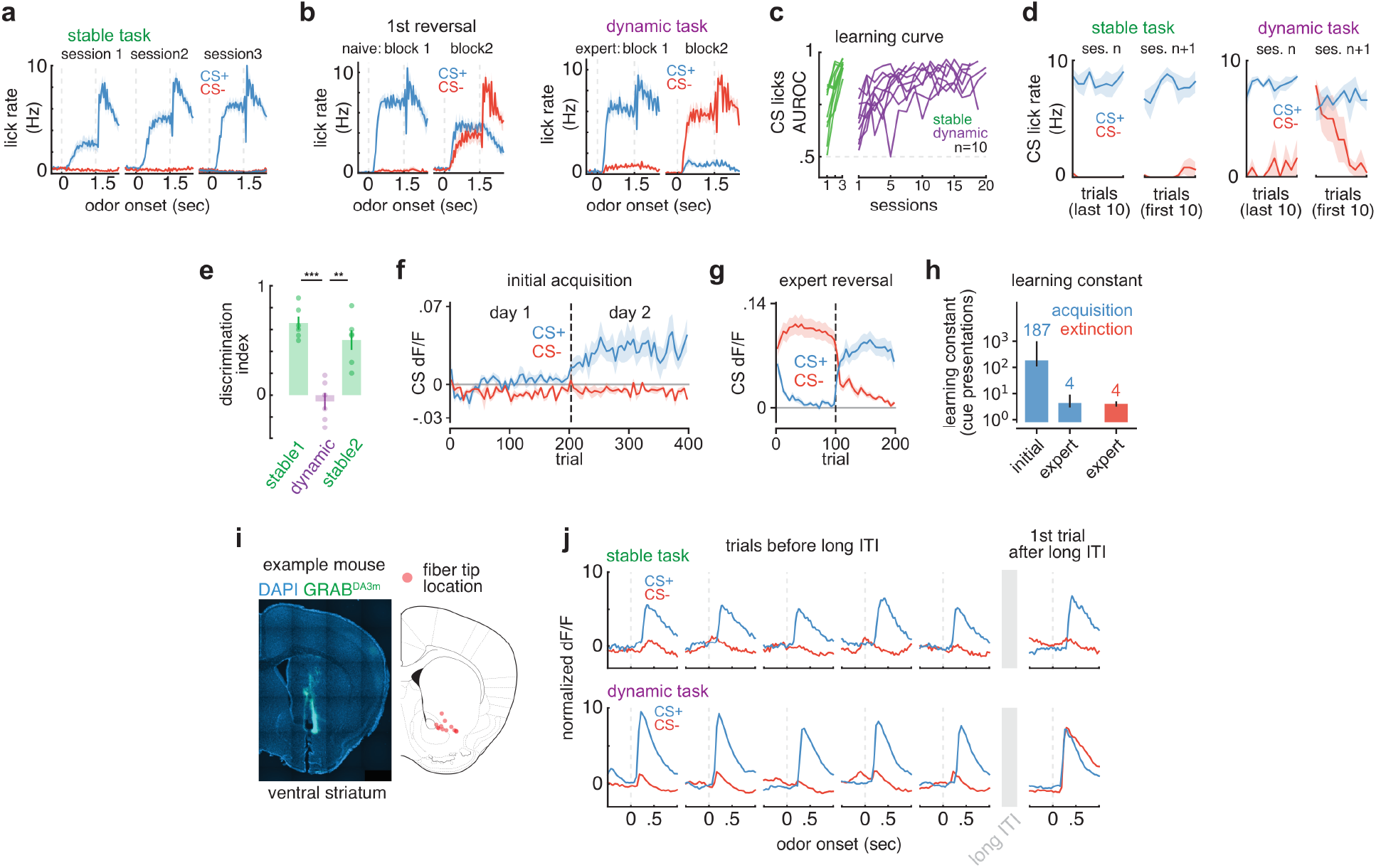
Further quantification of behavioral training, forgetting after breaks, and dopamine signaling related to forgetting. **a**, Lick rate to the CS+ and CS- aligned to odor onset in sessions 1-3 of the stable task (dashed lines, odor onset and offset; n=10 mice). **b**, Same as a in blocks 1 and 2 of the first reversal session of naïve mice (left) and of an expert session (right); CS+ (blue) and CS- (red) as in block 1 (n=10 mice). **c**, AUROC between CS+ and CS- lick rates across sessions of the stable and dynamic task (see Methods); each line, one mouse; dashed line, chance (n=10 mice for each task). **d**, CS lick rate over the last 10 trials of session n and the first 10 trials of session n+1 after a 1-day break, in the stable (left) and dynamic (right) task (trial, the n-th presentation of that cue; CS+ and CS-, the contingency at the end of session n) (stable, n=6 mice; dynamic, n=5 mice). **e**, Discrimination index on the first presentation of each cue in the last session of each stage, for mice trained in sequence on the stable (stable1), dynamic and again the stable task (stable2; see Methods); bars, mean; dots, mice (n=6 mice; two-tailed paired t-test). **f**, Dopamine cue response to the CS+ and CS- over days 1 and 2 of the stable task, against the trial number of the session (both cues counted; dashed line, day boundary; grey line, zero; see Methods) (n=5 mice). **g**, Same as f for expert reversals of the dynamic task (dashed line, reversal); CS+, the odor rewarded after the reversal; CS-, the odor rewarded before it (n=109 sessions from 25 mice). **h**, Learning constant of the dopamine cue response, in presentations of that cue (log scale), for acquisition (blue; initial, n=5 mice; expert, n=95 sessions from 25 mice) and extinction (red; expert, n=107 sessions from 25 mice) (see Methods); bars, median; error bars, 95% bootstrap confidence interval. **i**, *left*, GRAB^DA3m^ (green) and DAPI (blue) in the ventral striatum of an example mouse. *right*, Fiber tip locations (red) (n=12 mice). **j**, Example sessions: dopamine response to the CS+ and CS- on the last five trials of each cue before and the first trial of each cue after a 300-s long ITI (grey box; trials counted separately for each cue; cf. Fig. 1m), in the stable (top) and dynamic (bottom) task (dashed lines, odor onset). **, *P*<0.01; ***, *P*<0.001. All data shown are mean ± s.e.m. unless stated otherwise.

**Extended Data Fig 2.**
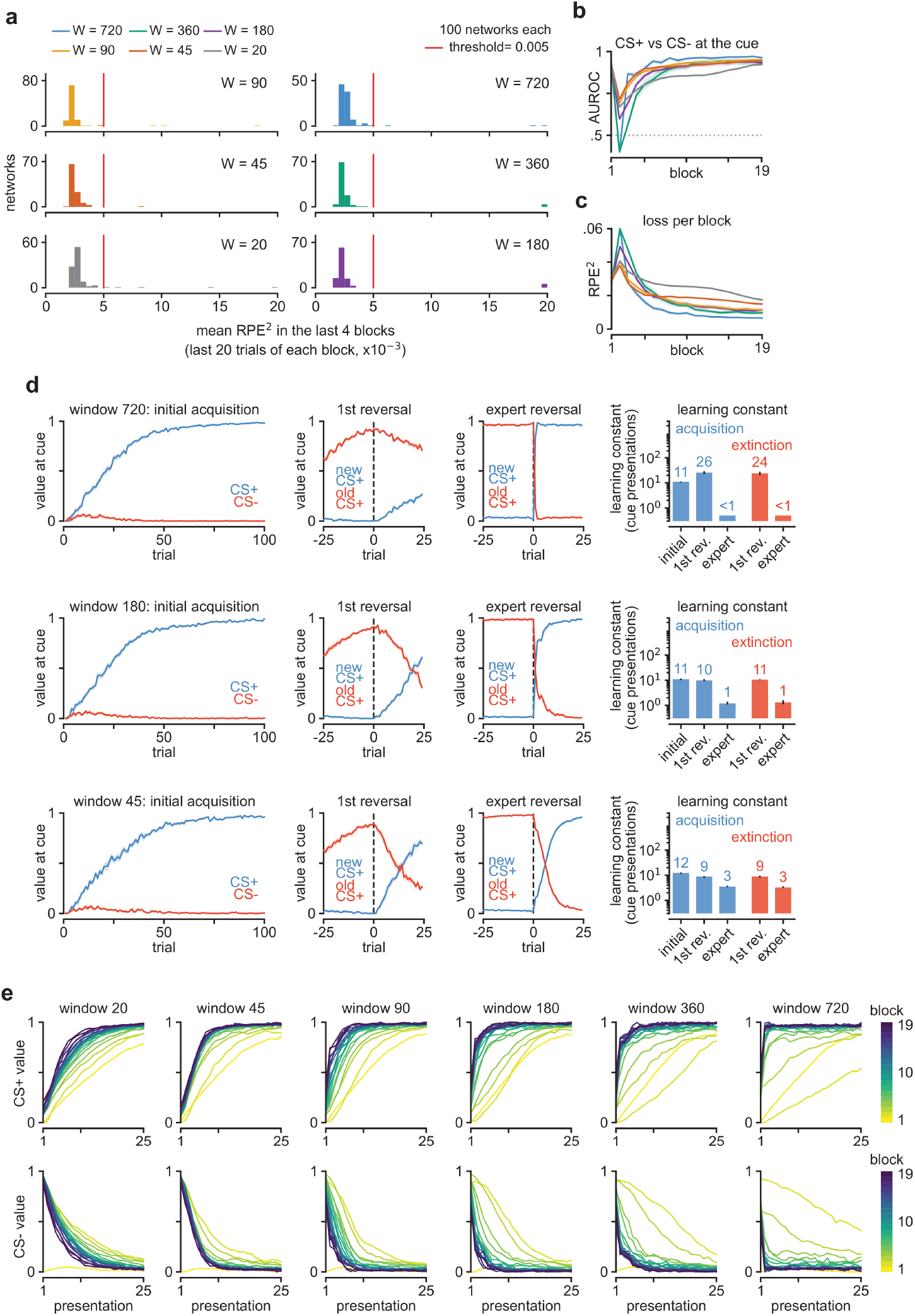
RNN modelling with different TBPTT windows. **a**, Convergence. Distribution of the mean loss (RPE^2^) over the last 20 trials of blocks 16-19 of the dynamic task for each window size W (the last bin includes all larger values); red line, threshold for convergence (see Methods) (n=100 networks per window). **b**, AUROC of the CS+ against the CS- value at the cue in each block of the dynamic task (values below the axis drawn at its limit); colors as in a; dotted line, chance (n=100 networks per window, converged or not). **c**, Mean RPE^2^ over all trials of each block of the dynamic task (values above the axis drawn at its limit); colors as in a (n=100 networks per window). **d**, Value at the cue (CS value) and learning constants of networks trained with W=720, 180 and 45 (rows, top to bottom; trial, the trial of the training stream, both cues counted). Initial acquisition, value of the CS+ (blue) and CS- (red) under one fixed contingency (n=50 networks per window). 1st reversal and expert reversal, value of the new CS+ (blue) and old CS+ (red) around the first reversal and the reversals into blocks 11-19 (dashed line, first trial of the new block; n=50 converged networks per window). Learning constant, in presentations of that cue (log scale), for acquisition (blue) and extinction (red) (see Methods); bars, median (<1, at the lower limit of the fit); error bars, 95% bootstrap confidence interval (n=50 networks per window). **e**, Learning curves of every block of the dynamic task for each window size: value of the CS+ (top) and CS- (bottom) of that block against the presentation of that cue within the block; one line per block, from block 1 (yellow) to block 19 (dark purple) (mean of the first 50 converged of 60 networks per window). All data shown are mean ± s.e.m. over networks unless stated otherwise.

**Extended Data Fig 3.**
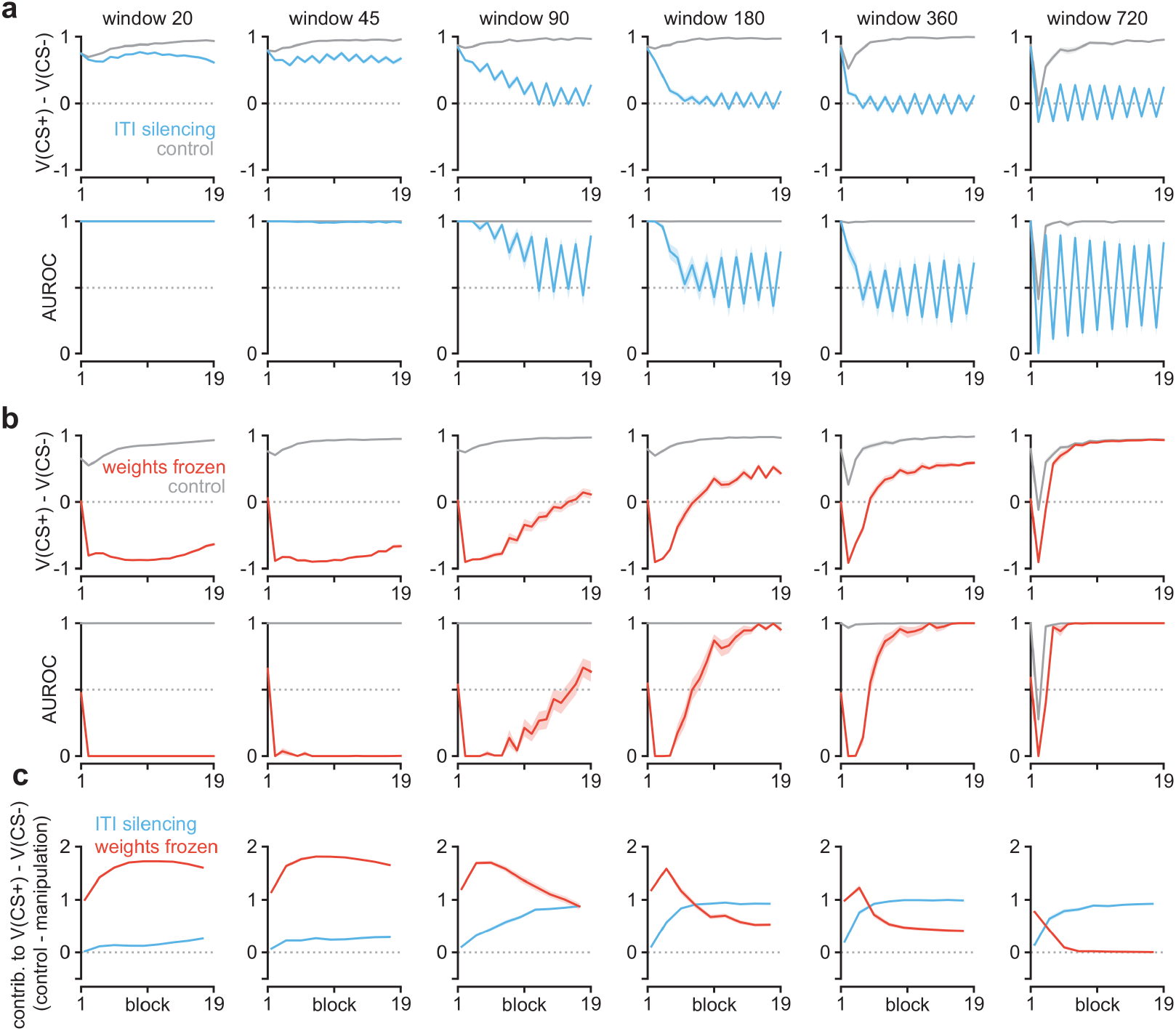
Effects of ITI silencing and weight freezing in RNNs trained with different TBPTT window sizes. Networks were trained online on the dynamic task (19 blocks) with TBPTT windows of 20-720 timesteps (columns), and each manipulation was tested in every block of training without perturbing the training itself (see Methods). **a**, ITI silencing (as in Fig. 2i). *top*, V(CS+) − V(CS-) on silenced (light blue) and control (grey) probe trials across blocks. *bottom*, AUROC between the CS+ and CS- probe values across networks in each block. **b**, Same as a for weights frozen (red; as in Fig. 2f) and control (grey), with the values averaged over the last 25 trials of each block. **c**, Contribution of each mechanism to V(CS+) − V(CS-), control − manipulation, for ITI silencing (light blue) and weights frozen (red); consecutive blocks averaged in pairs (see Methods). Dotted lines, zero (a, b, top; c) and chance (a, b, bottom). n=50 converged networks for each window and manipulation. All data shown are mean ± s.e.m. over networks (AUROC panels, bootstrap s.e.m. over networks).

**Extended Data Fig 4.**
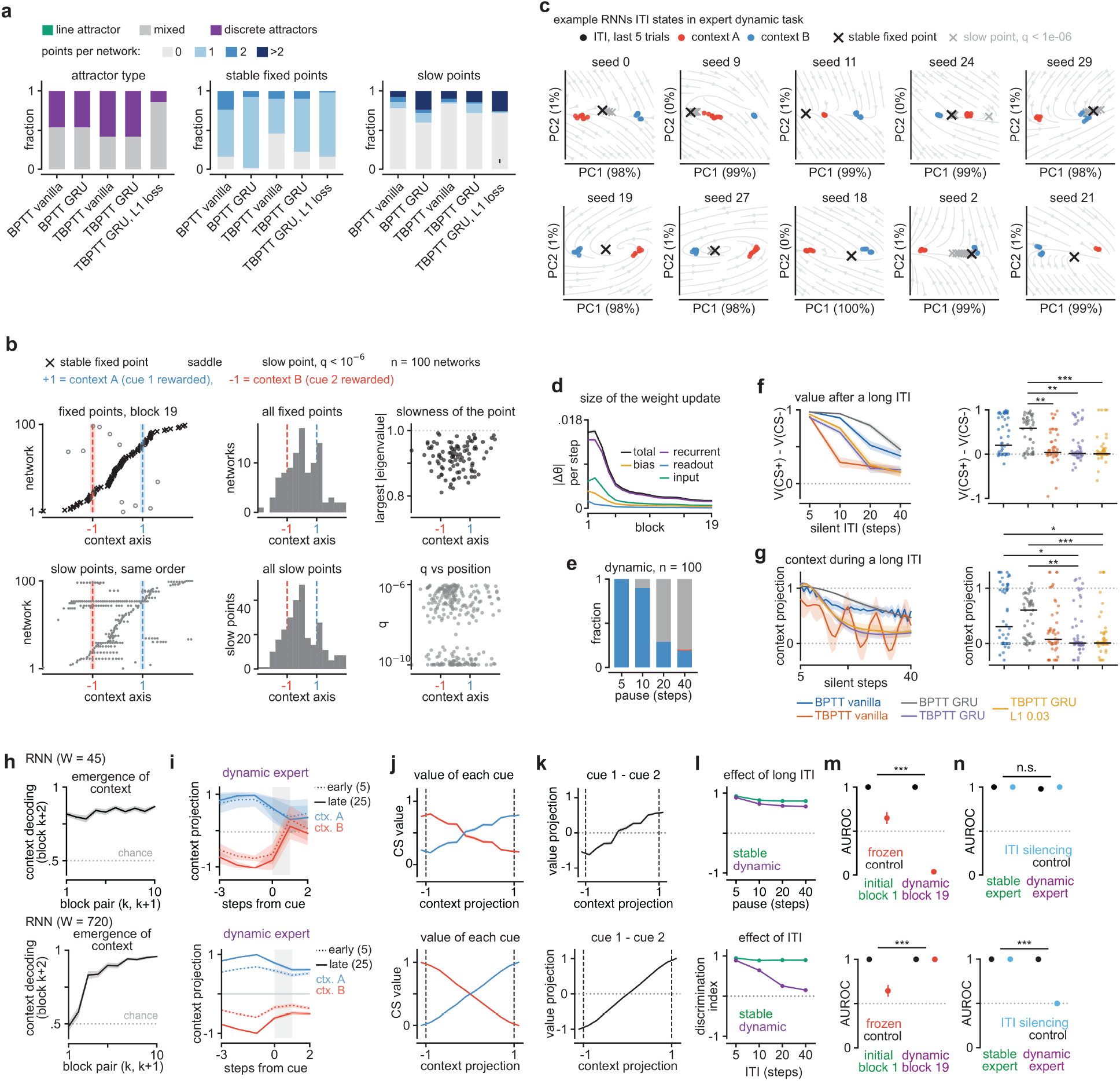
Attractor structure of the trained RNNs and the effect of the TBPTT window. **a**, Autonomous dynamics of networks trained on the dynamic task with five training rules (BPTT, full backpropagation through time; TBPTT, online with W=720; vanilla, tanh RNN; L1 loss, the activity cost of the main figures; see Methods). *left*, Fraction of networks of each attractor type, classified by three criteria (see Methods); *middle*, fraction of networks by number of stable fixed points; *right*, same for slow points (n=50 networks per rule). **b**, Fixed points (top; saddles, open grey circles) and slow points (bottom; grey dots) of expert TBPTT GRU + L1 networks on the context axis (+1, context A; −1, context B; shading, ±1 s.d. of the ITI states of a context): per network, sorted by the position of its stable fixed point (left), histogram (middle), and largest |eigenvalue| (top) or q (bottom) against position (right) (n=100 networks). **c**, ITI states of ten example expert networks (seed numbers) in the plane of their first two principal components (variance explained in parentheses), with the autonomous flow field (grey), as in Fig. 2j; context A, the contingency of the last training block. **d**, Size of the weight update per gradient step (|Δθ|) across training, for all parameters (total) and each parameter group (n=100 networks). **e**, Fraction of expert networks whose cue values after a silent pause kept the pre-pause contingency (blue), flipped it (red) or lay halfway (grey) (see Methods) (n=100 networks). **f**, *left*, V(CS+) − V(CS-) on the first trial after a silent ITI for the five rules of a; *right*, the same per network after 40 silent steps (dots, values beyond the axis drawn at its edge; bar, median) (n=50 networks per rule; two-tailed Mann-Whitney U test on all ten pairs, Holm-corrected; only significant pairs marked). **g**, Same as f for the context projection during the silent ITI (as in Fig. 2p; 1, the settled context; 0, the midpoint; see Methods) (n and statistics as in f). **h-n**, Networks trained with a short (W=45, top) or the standard (W=720, bottom) window. **h**, Emergence of a context code (as in Fig. 2k) (W=45, n=99; W=720, n=100 networks). **i**, Context projection around the cue (as in Fig. 2l) for ctx. A (cue 1 rewarded) and ctx. B (cue 2 rewarded), early (dotted) and late (solid) in a block; grey band, cue-to-outcome interval (W=45, n=99; W=720, n=100 networks). **j**, Value of cue 1 (blue) and cue 2 (red) reported from each ITI state against its context projection (as in Fig. 2m; mean ± s.e.m. over trials pooled across networks); dashed lines, ctx. A (+1) and ctx. B (−1) (W=45, n=99; W=720, n=100 networks). **k**, Value projection, V(cue 1) − V(cue 2), against the context projection (as in Fig. 2n; dashed lines as in j). **l**, Discrimination index after a silent pause (as in Fig. 2r; W=45 dynamic networks probed in the context of their own last block, see Methods) for networks after one block of the stable task (green) and expert dynamic-task networks (purple) (W=45: stable n=50, dynamic n=99; W=720: stable n=50, dynamic n=100 networks). **m**, AUROC of the CS+ against the CS- values with the weights frozen for the whole block (red; as in Fig. 2f) or plastic (control, black) in the initial block (block 1) and an expert reversal (block 19); stars compare the effect of freezing (control minus frozen AUROC) between the blocks (n=50 networks per window; two-tailed Wilcoxon signed-rank test). **n**, AUROC with ITI silencing (light blue; see Methods; W=45 dynamic networks probed in the context of their own last block) or control (black) in stable-expert (green) and dynamic-expert (purple) networks; stars compare the effect of silencing (control minus silenced AUROC) between the groups (n=50 networks per group and window; two-tailed Mann-Whitney U test). *, *P*<0.05; **, *P*<0.01; ***, *P*<0.001; n.s., *P*>0.05. All data shown are mean ± s.e.m. over networks unless stated otherwise.

**Extended Data Fig 5.**
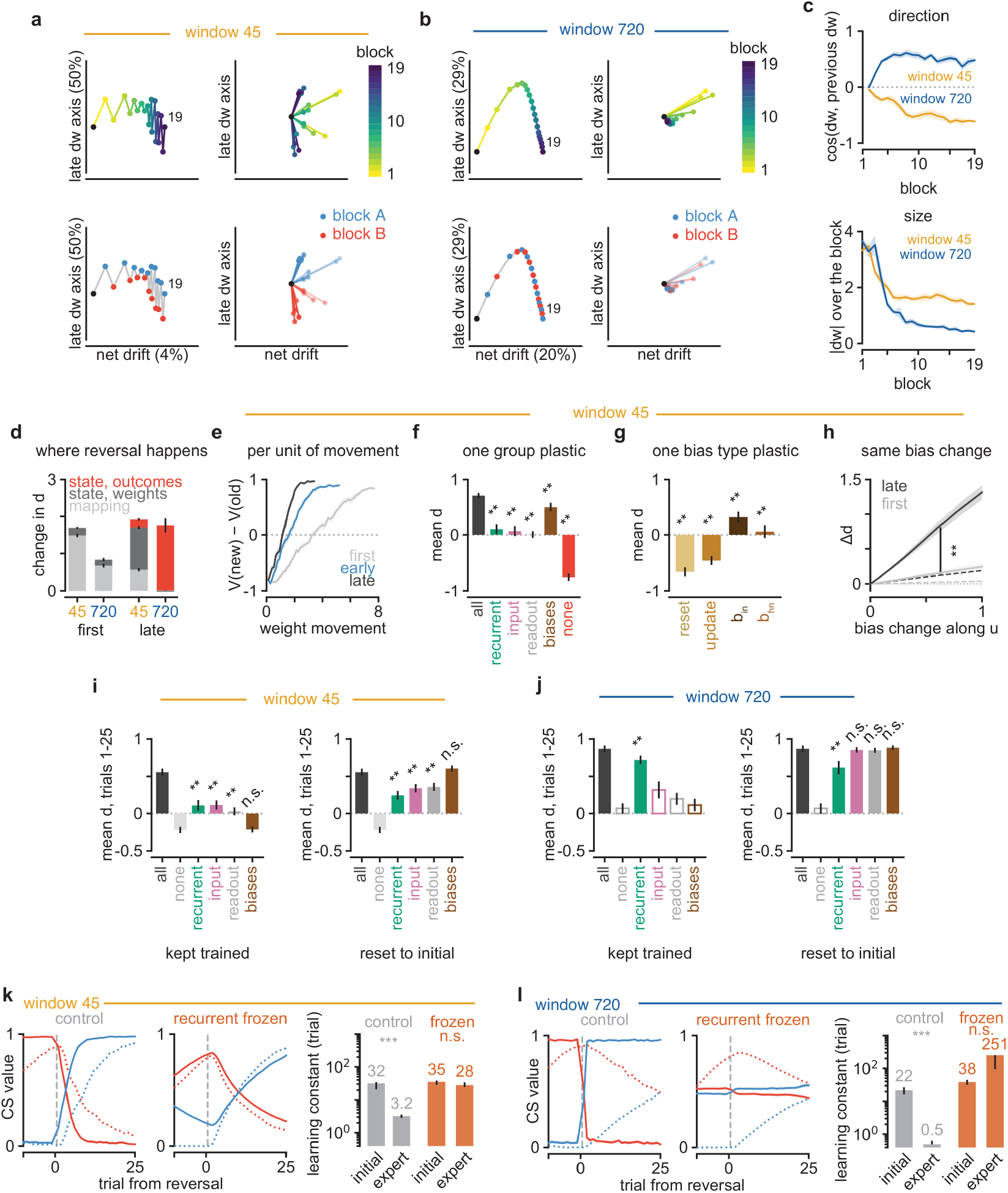
Weight-space mechanism of reversal learning in RNNs with a short versus a long TBPTT window. Networks trained online on the dynamic task with a short (W=45, orange) or a long (W=720, blue) TBPTT window (see Methods). Throughout, d = V(new CS+) − V(old CS+) at the cue; first, early and late, the reversals into block 2, block 4 and blocks 16 and 18 (averaged per network). **a**, Weight-space trajectory of an example network trained with W=45. *top*, *left*, Weight vector at the end of every block projected onto the net-drift and late-dw axes (see Methods), coloured by block (colour scale at right; black dot, initial weights; 19, end of block 19; numbers in parentheses, share of the squared late weight changes along each axis; axes scaled independently). *top*, *right*, Weight change of every block (dw) drawn from a common origin. *bottom*, Same as *top*, coloured by block type (block A, cue 1 rewarded; block B, cue 2 rewarded; grey line, path; fainter, earlier blocks). **b**, Same as a for an example network trained with W=720. **c**, Direction (top; cosine between the weight changes of consecutive blocks) and size (bottom) of the weight change of each block (n=10 networks per window, separate from those of d-j). **d**, Where the reversal happens. Change of d at the resting state across a reversal block, split into the change of the state-to-value mapping and the moves of the resting state produced by the new weights and by the new outcomes alone (see Methods); bars stack the three parts (negative parts drawn downward); error bars, s.e.m. of each part (n=10 networks per window). **e**, Value updating per unit of weight movement. d against the cumulative weight movement within the reversal block (W=45) for the first (grey), early (blue) and late (black) reversals (n=10 networks). **f**, One group plastic. Mean d over block 16 when training was continued from its start with only one parameter group plastic (W=45; none, weights frozen; see Methods); stars, versus all parameters plastic (n=10 networks). **g**, One bias type plastic. Same as f with one type of GRU bias plastic: the reset-gate, update-gate, input-side (b_in_) or hidden-side (b_hn_) candidate biases (n=10 networks). **h**, Same bias change. Change of d at the resting state (Δd) when the biases were moved by ε along the switch axis u with all weights frozen (see Methods), at the start of the first (grey) and the late (black) reversals (W=45); dashed lines, mean |Δd| for random bias directions (first, grey; late, black); vertical bar, first versus late at ε=0.6 (n=10 networks). **i**, Where the learned speed is stored. Mean d over the first 25 trials of block 19 of hybrid networks (W=45) built from the weights at the start of block 18 and trained online through blocks 18 and 19 (see Methods). *left*, One group kept trained and all others reset to their initial values (stars, versus none); *right*, one group reset to its initial values and all others kept trained (stars, versus all) (n=10 networks). **j**, Same as i for W=720; open bars, conditions that did not learn the old contingency in block 18, not tested (n=10 networks). **k**, Meta-learning without recurrent plasticity (W=45). *left* and *middle*, Value of the new CS+ (blue) and the old CS+ (red) at the cue by trial (the n-th presentation of that cue) from the reversal (dashed line), at the first reversal (dotted; into block 2) and at the expert reversals (solid, shading s.e.m.; into blocks 16-19, averaged per network), in control networks (all weights plastic) and in networks whose recurrent weights and biases were frozen at their initial values from the start of training (recurrent frozen; see Methods). *right*, Learning constant (trials, log scale) of the CS+ at initial acquisition (block 1) and of the new CS+ at the expert reversals (per-network median over the reversals into blocks 16-19) (see Methods); bars, median; lines, 95% bootstrap confidence interval; stars, initial versus expert within a condition (n=50 networks per condition). **l**, Same as k for W=720 (n=50 networks per condition). In a-j, dotted lines, 0. Two-tailed Wilcoxon signed-rank test (**, *P*<0.01; ***, *P*<0.001; n.s., *P*>0.05). All data shown are mean ± s.e.m. over networks, except the learning constants in k and l (median with 95% bootstrap confidence interval).

**Extended Data Fig 6.**
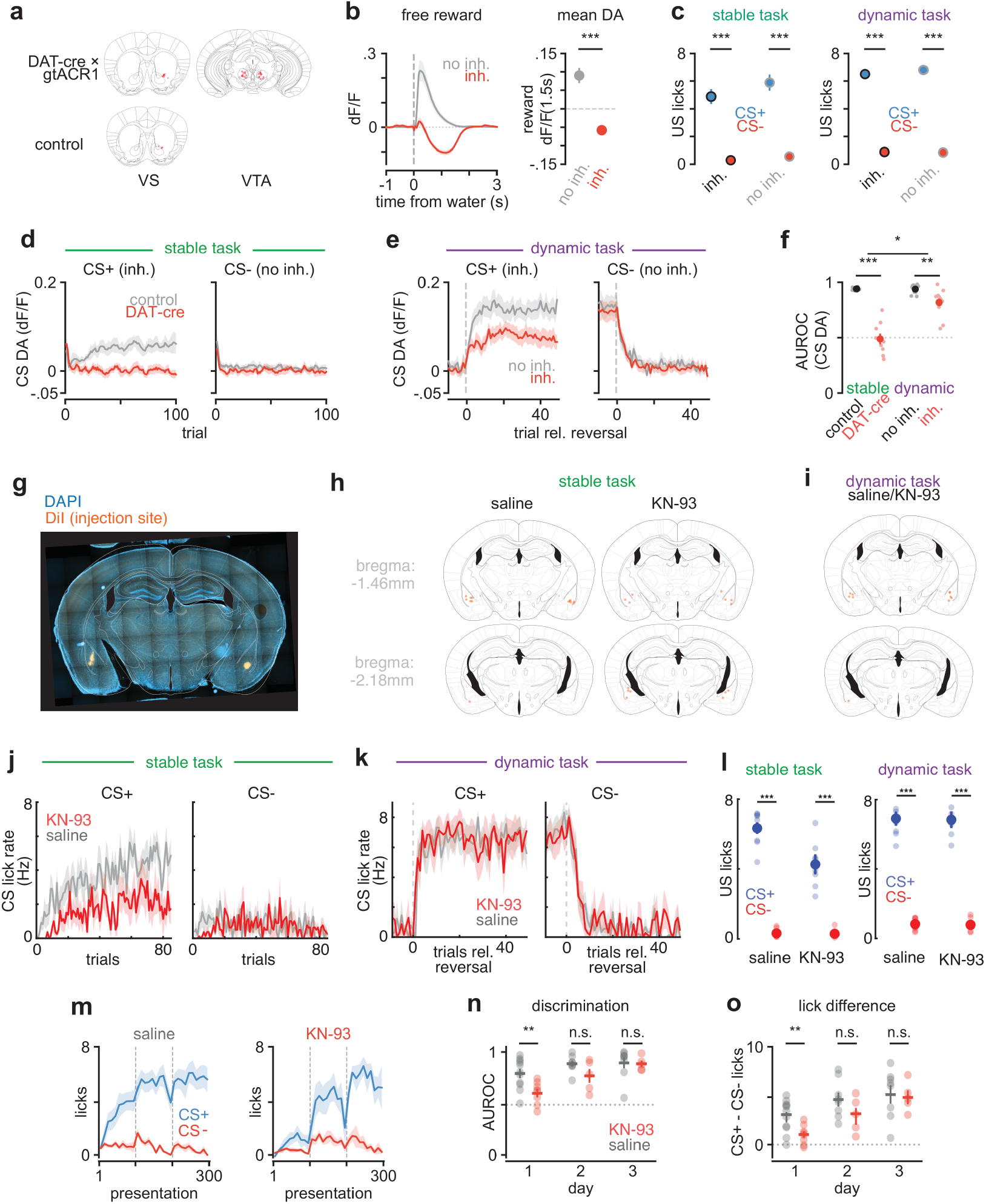
Histology and further quantification of the dopamine-inhibition and KN-93 infusion experiments. **a**, Fiber tips (red dots) above the ventral striatum (VS, left; photometry) of DAT-cre × gtACR1 (*top*, n=11 mice) and control (*bottom*, n=5 mice) mice and above the VTA (right) of DAT-cre × gtACR1 mice (n=11 mice). **b**, Calibration of the dopamine inhibition (see Methods). *left*, Dopamine response to free water (dashed line, water delivery; dotted line, 0) without (no inh.) or with light (inh.). *right*, Mean response over the 1.5 s after the water (dashed line, 0) (n=13 mice; two-tailed paired t-test). **c**, US (consumption) licks on CS+ and CS- trials (see Methods) in the stable task (left; DAT-cre × gtACR1 mice with light at the CS+ outcome, inh., n=9 mice; control mice without the opsin given the same light, no inh., n=7 mice) and in the light (inh.) and no-light (no inh.) sessions of the dynamic task (right; inh., 26 sessions from n=11 mice; no inh., 31 sessions from n=13 mice) (two-tailed paired t-test over mice, CS+ versus CS- within each group). **d**, Cue dopamine response (CS DA) by presentation of each cue (trial, the n-th presentation of that cue) on day 1 of the stable task (control, n=5 mice; DAT-cre × gtACR1, n=9 mice; see Methods). **e**, Same as d for the dynamic task, aligned to the reversal (dashed line; CS+, the odor rewarded after the reversal, also in k); light (inh.) and no-light (no inh.) sessions of the same mice (inh., n=10 mice, 23 sessions; no inh., n=12 mice, 20 sessions). **f**, AUROC of the CS DA between CS+ and CS- trials (see Methods); pale dots, mice; filled dots, mean; dotted line, chance (stable, n=5 control and n=9 DAT-cre mice, two-tailed Student’s t-test; dynamic, n=12 (no inh.) and 10 (inh.) mice, sessions as in e, two-tailed paired t-test on the 10 mice with both conditions; top bracket, task difference of the inhibition effect, label-permutation test). **g**, Example histology of an amygdala infusion site (DiI; see Methods). **h**, Injection sites (orange dots) of the stable-task infusions (Fig. 3i-l) for saline (left) and KN-93 (right) mice. **i**, Same as h for the dynamic task (same mice for both drugs). **j**, Stable task (day 1). CS licks by presentation of each cue (saline, n=11 mice; KN-93, n=8 mice). **k**, Dynamic task (expert mice). CS licks aligned to the reversal (dashed line) on KN-93 and saline sessions of the same mice (n=9 saline and 7 KN-93 sessions from 5 mice). **l**, US licks on CS+ and CS- trials in the stable (left; pale dots, mice; saline, n=11 mice; KN-93, n=8 mice) and dynamic (right; pale dots, sessions; n=9 saline and 7 KN-93 sessions from 5 mice) task; filled dots, mean (two-tailed paired t-test, CS+ versus CS- within each group, over mice in the stable task and over sessions in the dynamic task). **m**, CS licks over days 1-3 of the stable task (dashed lines, day boundaries; see Methods) in the mice that completed all three days (saline, n=8 mice; KN-93, n=5 mice). **n**, Discrimination (AUROC of CS licks, CS+ versus CS-) per day; dots, mice; horizontal bar, mean; vertical line, s.e.m.; dotted line, chance. **o**, CS+ − CS- licks per day (dotted line, 0) (n, o: saline, n=11/8/8 mice and KN-93, n=8/5/5 mice on days 1/2/3; two-tailed Student’s t-test, saline versus KN-93). *, *P*<0.05; **, *P*<0.01; ***, *P*<0.001; n.s., *P*>0.05. All data shown are mean ± s.e.m. (over mice, except over sessions in k and in the dynamic task of l).

**Extended Data Fig 7.**
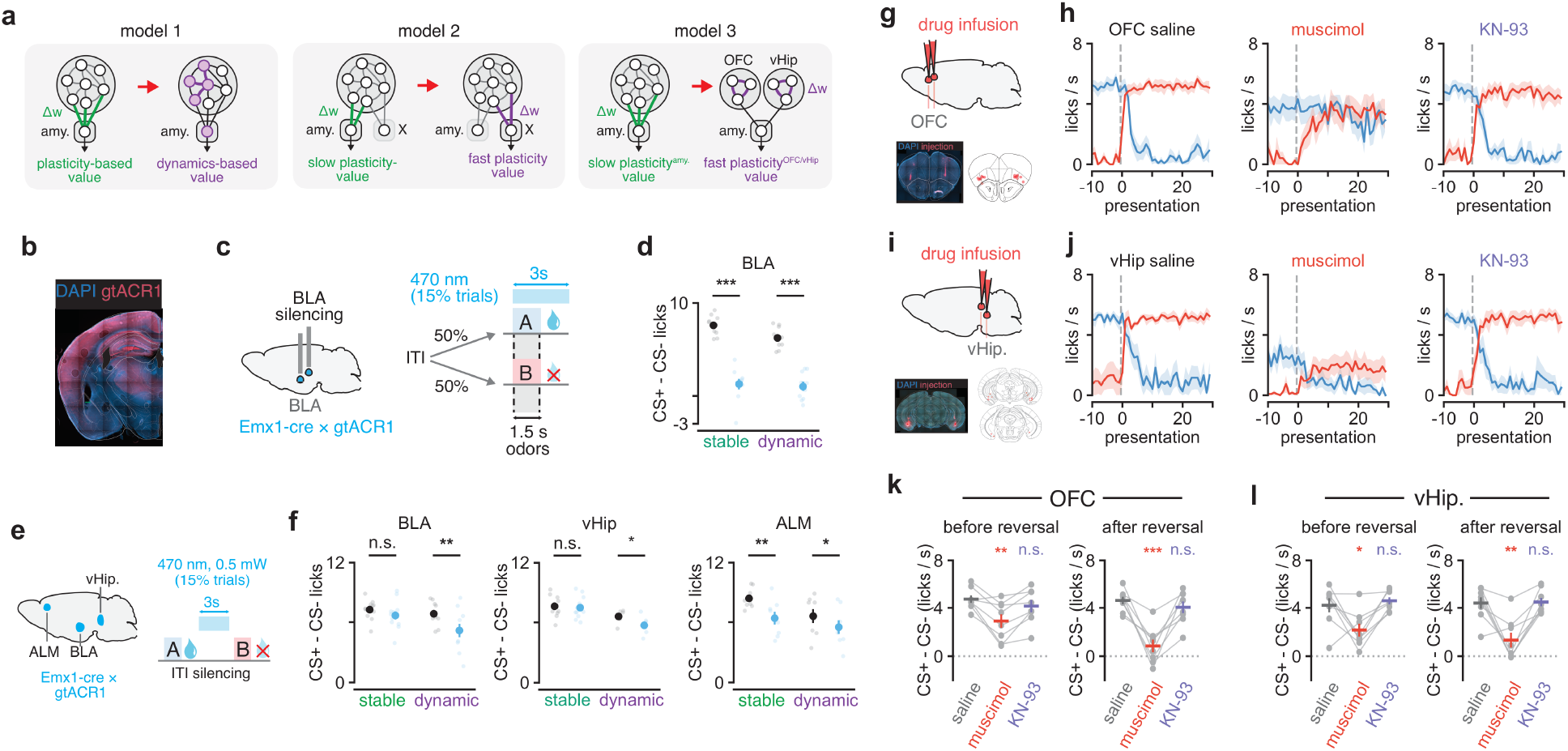
Alternative models of the plasticity-to-dynamics transition and amygdala, vHip, ALM and OFC manipulations. **a**, Three models consistent with the effect of blocking amygdala plasticity (Fig. 3i-l). Plasticity in the amygdala (Δw, green; slow in models 2 and 3) drives value updates in the stable task and, in the expert dynamic task, gives way to recurrent dynamics (dynamics-based value, purple; model 1) or to fast plasticity (purple) in another region X (model 2) or in the orbitofrontal cortex (OFC) or ventral hippocampus (vHip), which project to the amygdala (model 3). Model 2 predicts that amygdala activity is not necessary for the dynamic task. **b**, Example histology of an Emx1-cre × gtACR1 mouse (DAPI, blue; gtACR1, red). **c**, Schematic of cue silencing of the basolateral amygdala (BLA; A, rewarded odor, drop; B, unrewarded odor, cross; see Methods). **d**, Cue discrimination (CS+ − CS- licks) in the stable and dynamic task on trials without light (control, black) and with light (cue silenced, light blue); small dots, sessions; large dots, mean (stable, n=10 sessions from 5 mice; dynamic, n=10 sessions from 5 mice; two-tailed paired t-test, control versus silenced). Not drawn: cue silencing raised CS- licks in both tasks, whereas CS+ licks did not change (two-tailed paired t-test). **e**, Schematic of ITI silencing of the anterior lateral motor cortex (ALM), BLA or vHip (symbols as in c; see Methods). **f**, Same as d for ITI silencing of the BLA (left), vHip (middle) and ALM (right) (control, black; ITI silenced, light blue) (BLA: stable n=12 sessions from 6 mice, dynamic n=10 sessions from 5 mice; vHip: stable n=11 sessions from 7 mice, dynamic n=6 sessions from 6 mice; ALM: stable and dynamic, n=8 sessions from 4 mice each; two-tailed paired t-test, control versus silenced). Not drawn: ITI silencing of the BLA raised CS- licks in both tasks without changing CS+ licks, and ITI silencing of the vHip raised CS- licks in the dynamic task only (two-tailed paired t-test); the reduction of CS+ − CS- licks by silencing was larger in the dynamic than in the stable task for the BLA and the vHip, but not for the ALM (two-tailed Student’s t-test). **g**, Drug infusion in the OFC. *top*, Schematic. *bottom*, *left*, Example coronal slice (DAPI, blue; injection site, red). *bottom*, *right*, Injection sites of all mice (red). **h**, Lick rate around the reversal (dashed line) after saline (left), muscimol (middle) or KN-93 (right) infusion in the OFC, for the odor rewarded before (blue, CS+ → CS-) and after (red, CS- → CS+) the reversal (n=8 mice, one session per mouse and drug; see Methods). **i**, Same as g for the vHip. **j**, Same as h for vHip infusions (n=8 mice). **k**, Discrimination (CS+ − CS- lick rate under the current contingency; see Methods) before (left) and after (right) the reversal for saline (grey), muscimol (red) and KN-93 (purple) infusion in the OFC; grey dots and lines, mice; colored horizontal bar, mean; vertical bar, s.e.m.; dotted line, 0 (n=8 mice; two-tailed paired t-test versus saline; stars colored by drug). **l**, Same as k for the vHip (n=8 mice). *, *P*<0.05; **, *P*<0.01; ***, *P*<0.001; n.s., *P*>0.05. All data shown are mean ± s.e.m. (over sessions in d, f; over mice in h, j-l).

**Extended Data Fig 8.**
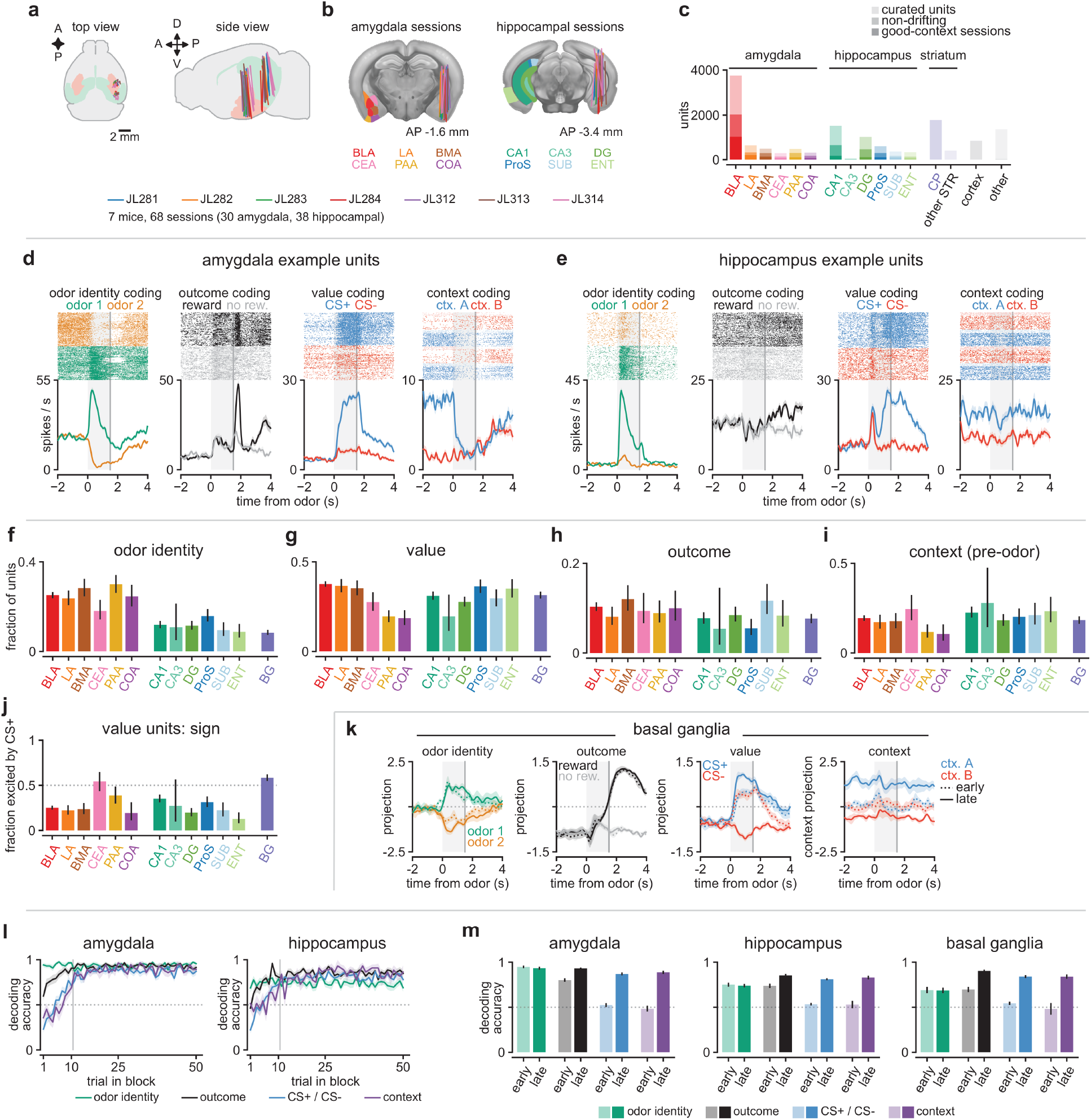
Neuropixels recordings in the amygdala and hippocampus during the dynamic task. **a**, Probe tracks of all sessions registered to the Allen atlas, one line per session colored by mouse (*left*, top view; *right*, side view; amygdala, salmon, and hippocampus, green, shaded) (n=7 mice, 68 sessions: 30 amygdala, 38 hippocampal). **b**, The same tracks on coronal slices at the median anterior-posterior position of the amygdala (left) and hippocampal (right) sessions, with the recorded subregions colored on the opposite hemisphere (abbreviations, see Methods). **c**, Number of units per region: curated units (light), non-drifting units (medium) and non-drifting units of the good-context sessions (dark) (see Methods); striatum (CP, caudoputamen; STR, other striatum), cortex and other regions, curated units only. **d**, Example amygdala units coding odor identity, outcome, value and context: spike rasters (top) and PSTHs (bottom) aligned to odor onset (grey shading, odor; vertical line, outcome); the context raster is in trial order, with the first 10 trials of each block drawn pale and excluded from the PSTH (see Methods). **e**, Same as d for the hippocampus. **f-i**, Fraction of units per subregion (colors as in b; BG, basal ganglia, purple) coding odor identity (f), value (g), outcome (h) and context (i; non-drifting units only) in a trial-by-trial regression (ordinary-least-squares t-test, *P*<0.01; see Methods); error bars, 95% Wilson confidence intervals (f-h, curated units: BLA, n=3,751; LA, n=659; BMA, n=514; CEA, n=285; PAA, n=503; COA, n=308; CA1, n=1,527; CA3, n=56; DG, n=1,032; ProS, n=621; SUB, n=359; ENT, n=333; BG, n=2,192 units; i, non-drifting units: BLA, n=2,023; LA, n=327; BMA, n=295; CEA, n=138; PAA, n=291; COA, n=180; CA1, n=670; CA3, n=25; DG, n=483; ProS, n=289; SUB, n=169; ENT, n=132; BG, n=1,126 units). Not drawn: subregions and regions were compared with chi-square tests (see Methods). **j**, Fraction of the value units of g excited by the CS+ (error bars, 95% Wilson confidence intervals; dotted line, 0.5; BLA, n=1,413; LA, n=242; BMA, n=182; CEA, n=79; PAA, n=98; COA, n=57; CA1, n=476; CA3, n=11; DG, n=287; ProS, n=226; SUB, n=107; ENT, n=117; BG, n=691 value units). Not drawn: subregions were compared with chi-square tests, and each region with 0.5 with a two-tailed binomial test. **k**, Population projections onto the dPCA odor-identity, outcome, value and context axes in the basal ganglia, as in Fig. 4c, e (see Methods). Solid and dotted lines, CS+ and CS- trials (odor identity), the two odors (outcome), and late (last 10 trials of a block) and early (first 10 trials after a reversal) trials (value, context); grey shading, odor; vertical line, outcome; dotted line, 0 (n=21 sessions with ≥20 non-drifting units; context, n=7 good-context sessions). **l**, Decoding accuracy by trial position (trial in block, counting the trials of both odors) in the amygdala (left) and hippocampus (right), for decoders trained on the trials after the first 10 of each block (see Methods); dotted line, chance; vertical line, trial 10 (amygdala, n=29 sessions, context n=14 good-context sessions; hippocampus, n=33 sessions, context n=11 good-context sessions). Not drawn: accuracy on the first trial after a reversal was compared with 0.5 (two-tailed Wilcoxon signed-rank test over sessions). **m**, Balanced decoding accuracy on the early (trials 1-10 after a reversal, light) and held-out late (dark) trials in the amygdala (left), hippocampus (middle) and basal ganglia (right; n=21 sessions, context n=7 good-context sessions); n as in l; dotted line, chance. Not drawn: early versus late accuracy, and early accuracy versus 0.5, two-tailed Wilcoxon signed-rank test over sessions. All data shown are mean ± s.e.m. (d, e, over trials; k-m, over sessions).

**Extended Data Fig 9.**
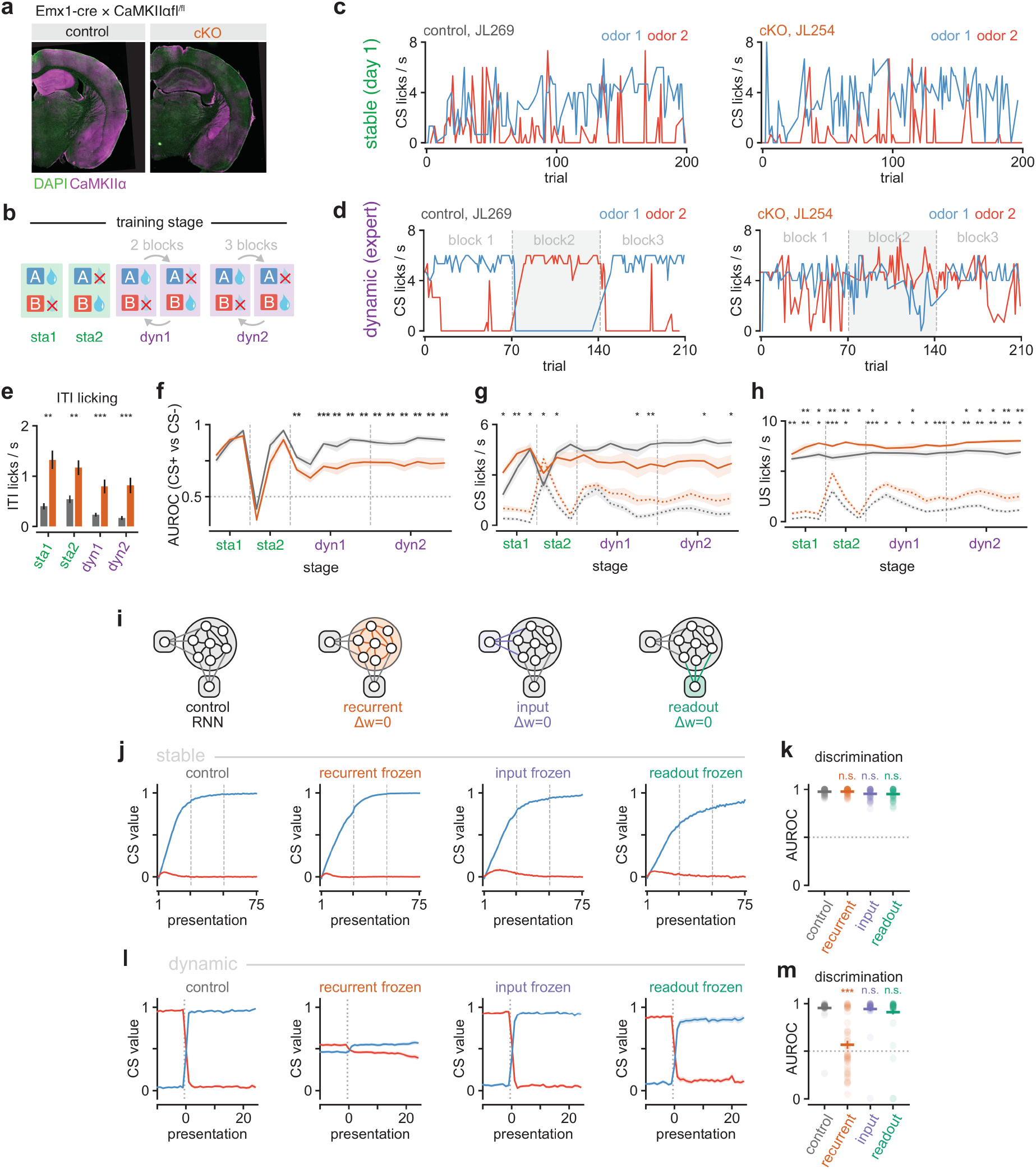
CaMKIIα conditional knockout and freezing the recurrent weights of RNNs spare stable learning but impair the dynamic task. **a**, Example histology of a control (Cre-negative littermate; left) and a conditional knockout (cKO; right) mouse (genotypes, see Methods). **b**, Training stages (see Methods): stable task (sta1), reversed stable task (sta2) and dynamic task with one (dyn1) or two (dyn2) reversals per session; A, odor A; B, odor B; drop, rewarded; cross, unrewarded; arrows, reversals. **c**, Example sessions on day 1 of the stable task (sta1) of a control (left) and a cKO (right) mouse: single-trial CS lick rate against the trial of the session (odor 1, odor A; odor 2, odor B). **d**, Same mice on the last dyn2 session (expert); dashed lines, block boundaries; grey shading, block 2, in which odor 2 was rewarded. **e**, ITI lick rate at each stage for control (grey) and cKO (orange) mice (n=10 mice per group; two-tailed Mann-Whitney U test). **f**, Discrimination (AUROC of the CS lick rate, CS+ versus CS-) on every session of the four stages for control (grey) and cKO (orange) mice; dashed lines, stage boundaries (also in g, h); dotted line, chance (n=10 mice per group, except 9 control mice on sta1 session 3 and dyn2 session 4; two-tailed Mann-Whitney U test per session). **g**, CS lick rate to the CS+ (solid) and CS- (dotted) per session (see Methods); stars, CS- (n as in f; two-tailed Mann-Whitney U test per session). **h**, US lick rate on CS+ (reward, solid) and CS- (omission, dotted) trials per session; stars, CS+ (top row) and CS- (bottom row) (n as in f; two-tailed Mann-Whitney U test per session). **i**, The four online RNNs (see Methods): control (all weights plastic) and networks with the recurrent (RNN^Δw=0^ of Fig. 5e), input or readout weights frozen (Δw=0). **j**, Stable task (3 blocks of 50 trials; dashed lines, block boundaries). Value at the cue for the CS+ (blue) and CS- (red) by presentation of each cue (n=50 networks per condition). **k**, AUROC between CS+ and CS- values in the stable task; dots, networks; bar, mean; line, s.e.m.; dotted line, chance (n=50 networks per condition; two-tailed Mann-Whitney U test versus control). **l**, Expert dynamic task (last reversal, block 18 → 19, as in Fig. 5g). Value at the cue for the new CS+ (blue) and old CS+ (red) aligned to the reversal (dotted line) (n=50 networks per condition). **m**, Same as k for the expert dynamic task (block 19) (n=50 networks per condition; two-tailed Mann-Whitney U test versus control). *, *P*<0.05; **, *P*<0.01; ***, *P*<0.001; n.s., *P*>0.05. All data shown are mean ± s.e.m. (over mice in e-h; over networks in j-m).

## References

1. Sutton, R. S. & Barto, A. G. Reinforcement Learning: An Introduction. (A Bradford Book, Cambridge, Mass, 1998).

2. Botvinick, M., Wang, J. X., Dabney, W., Miller, K. J. & Kurth-Nelson, Z. Deep Reinforcement Learning and Its Neuroscientific Implications. Neuron 107, 603–616 (2020).

3. Lee, D., Seo, H. & Jung, M. W. Neural Basis of Reinforcement Learning and Decision Making. Annual Review of Neuroscience 35, 287–308 (2012).

4. Schultz, W., Dayan, P. & Montague, P. R. A Neural Substrate of Prediction and Reward. Science 275, 1593–1599 (1997).

5. Reynolds, J. N. J., Hyland, B. I. & Wickens, J. R. A cellular mechanism of reward-related learning. Nature 413, 67–70 (2001).

6. Hong, S. & Hikosaka, O. Dopamine-Mediated Learning and Switching in Cortico-Striatal Circuit Explain Behavioral Changes in Reinforcement Learning. Front. Behav. Neurosci. 5, (2011).

7. Tye, K. M., Stuber, G. D., de Ridder, B., Bonci, A. & Janak, P. H. Rapid strengthening of thalamo-amygdala synapses mediates cue–reward learning. Nature 453, 1253–1257 (2008).

8. Yamaguchi, K. et al. A behavioural correlate of the synaptic eligibility trace in the nucleus accumbens. Sci Rep 12, 1921 (2022).

9. Xiong, Q., Znamenskiy, P. & Zador, A. M. Selective corticostriatal plasticity during acquisition of an auditory discrimination task. Nature 521, 348–351 (2015).

10. Tsutsui-Kimura, I. et al. Dopamine in the tail of the striatum facilitates avoidance in threat– reward conflicts. Nat Neurosci 28, 795–810 (2025).

11. Rogan, M. T., Stäubli, U. V. & LeDoux, J. E. Fear conditioning induces associative long-term potentiation in the amygdala. Nature 390, 604–607 (1997).

12. Reynolds, J. N. J. & Wickens, J. R. Dopamine-dependent plasticity of corticostriatal synapses. Neural Networks 15, 507–521 (2002).

13. Yagishita, S. et al. A critical time window for dopamine actions on the structural plasticity of dendritic spines. Science 345, 1616–1620 (2014).

14. Wang, J. X. et al. Prefrontal cortex as a meta-reinforcement learning system. Nat Neurosci 21, 860–868 (2018).

15. Hattori, R. et al. Meta-reinforcement learning via orbitofrontal cortex. Nat Neurosci 26, 2182–2191 (2023).

16. Bartolo, R. & Averbeck, B. B. Prefrontal Cortex Predicts State Switches during Reversal Learning. Neuron 106, 1044–1054.e4 (2020).

17. Vertechi, P. et al. Inference-Based Decisions in a Hidden State Foraging Task: Differential Contributions of Prefrontal Cortical Areas. Neuron 106, 166–176.e6 (2020).

18. Mishchanchuk, K. et al. Hidden state inference requires abstract contextual representations in the ventral hippocampus. Science 386, 926–932 (2024).

19. Hattori, R., Danskin, B., Babic, Z., Mlynaryk, N. & Komiyama, T. Area-Specificity and Plasticity of History-Dependent Value Coding During Learning. Cell 177, 1858–1872.e15 (2019).

20. Kim, C. M., Chow, C. C. & Averbeck, B. B. Neural dynamics of reversal learning in the prefrontal cortex and recurrent neural networks. eLife 13, (2025).

21. Bari, B. A. et al. Stable Representations of Decision Variables for Flexible Behavior. Neuron 103, 922–933.e7 (2019).

22. Bittner, K. C., Milstein, A. D., Grienberger, C., Romani, S. & Magee, J. C. Behavioral time scale synaptic plasticity underlies CA1 place fields. Science 357, 1033–1036 (2017).

23. Milstein, A. D. et al. Bidirectional synaptic plasticity rapidly modifies hippocampal representations. eLife 10, e73046 (2021).

24. Magee, J. C. Behavioral timescale synaptic plasticity: properties, elements and functions. Nat Neurosci 29, 520–534 (2026).

25. Yaeger, C. E., Soto-Albors, R. M., Liu, W., Beltramini, A. & Harnett, M. T. Plateau potentials are instructive signals for behavioral timescale synaptic plasticity in the neocortex. Preprint at 10.1101/2025.11.07.687250 (2025).

26. Pereira-Obilinovic, U., Hou, H., Svoboda, K. & Wang, X.-J. Brain mechanism of foraging: Reward-dependent synaptic plasticity versus neural integration of values. Proceedings of the National Academy of Sciences 121, e2318521121 (2024).

27. Parker, N. F., et al. Choice-selective sequences dominate in cortical relative to thalamic inputs to NAc to support reinforcement learning. Cell Reports 39, (2022).

28. Hennig, J. A. et al. Emergence of belief-like representations through reinforcement learning. PLOS Computational Biology 19, e1011067 (2023).

29. Qian, L. et al. Prospective contingency explains behavior and dopamine signals during associative learning. Nat Neurosci 28, 1280–1292 (2025).

30. Mante, V., Sussillo, D., Shenoy, K. V. & Newsome, W. T. Context-dependent computation by recurrent dynamics in prefrontal cortex. Nature 503, 78–84 (2013).

31. Driscoll, L. N., Shenoy, K. & Sussillo, D. Flexible multitask computation in recurrent networks utilizes shared dynamical motifs. Nat Neurosci 27, 1349–1363 (2024).

32. Steinberg, E. E. et al. A causal link between prediction errors, dopamine neurons and learning. Nat Neurosci 16, 966–973 (2013).

33. Iino, Y. et al. Dopamine D2 receptors in discrimination learning and spine enlargement. Nature 579, 555–560 (2020).

34. Lee, S. J. et al. Cell-type-specific asynchronous modulation of PKA by dopamine in learning. Nature 590, 451–456 (2021).

35. Lahiri, A. K. & Bevan, M. D. Dopaminergic Transmission Rapidly and Persistently Enhances Excitability of D1 Receptor-Expressing Striatal Projection Neurons. Neuron 106, 277–290.e6 (2020).

36. Ericsson, J. et al. Dopamine Differentially Modulates the Excitability of Striatal Neurons of the Direct and Indirect Pathways in Lamprey. J. Neurosci. 33, 8045–8054 (2013).

37. Maltese, M., March, J. R., Bashaw, A. G. & Tritsch, N. X. Dopamine differentially modulates the size of projection neuron ensembles in the intact and dopamine-depleted striatum. eLife 10, e68041 (2021).

38. Beyeler, A. et al. Organization of Valence-Encoding and Projection-Defined Neurons in the Basolateral Amygdala. Cell Reports 22, 905–918 (2018).

39. Paton, J. J., Belova, M. A., Morrison, S. E. & Salzman, C. D. The primate amygdala represents the positive and negative value of visual stimuli during learning. Nature 439, 865– 870 (2006).

40. Zhang, X. & Li, B. Population coding of valence in the basolateral amygdala. Nat Commun 9, 5195 (2018).

41. Zhang, X. et al. Genetically identified amygdala–striatal circuits for valence-specific behaviors. Nat Neurosci 24, 1586–1600 (2021).

42. Radiske, A., de Castro, C. M., Rossato, J. I., Gonzalez, M. C. & Cammarota, M. Hippocampal CaMKII inhibition induces reactivation-dependent amnesia for extinction memory and causes fear relapse. Sci Rep 13, 21712 (2023).

43. Adler, A., Zhao, R., Shin, M. E., Yasuda, R. & Gan, W.-B. Somatostatin-Expressing Interneurons Enable and Maintain Learning-Dependent Sequential Activation of Pyramidal Neurons. Neuron 102, 202–216.e7 (2019).

44. Cichon, J. & Gan, W.-B. Branch-specific dendritic Ca2+ spikes cause persistent synaptic plasticity. Nature 520, 180–185 (2015).

45. Radiske, A. et al. Avoidance memory requires CaMKII activity to persist after recall. Molecular Brain 14, 167 (2021).

46. Wilson, R. C., Takahashi, Y. K., Schoenbaum, G. & Niv, Y. Orbitofrontal Cortex as a Cognitive Map of Task Space. Neuron 81, 267–279 (2014).

47. Sanchez, A. N., Ford, C. F., Balewski, Z. Z. & Wallis, J. D. Orbitofrontal-Hippocampal State Coding Dynamics during Reversal Learning. J. Neurosci. 45, (2025).

48. Guo, Z. V. et al. Flow of Cortical Activity Underlying a Tactile Decision in Mice. Neuron 81, 179–194 (2014).

49. Li, N., Chen, T.-W., Guo, Z. V., Gerfen, C. R. & Svoboda, K. A motor cortex circuit for motor planning and movement. Nature 519, 51–56 (2015).

50. Achterberg, K. G. et al. Temporal and Region-Specific Requirements of αCaMKII in Spatial and Contextual Learning. J. Neurosci. 34, 11180–11187 (2014).

51. Abraham, W. C. & Bear, M. F. Metaplasticity: the plasticity of synaptic plasticity. Trends in Neurosciences 19, 126–130 (1996).

52. Saar, D., Grossman, Y. & Barkai, E. Reduced after-hyperpolarization in rat piriform cortex pyramidal neurons is associated with increased learning capability during operant conditioning. European Journal of Neuroscience 10, 1518–1523 (1998).

53. Cohen, Y., Avramoav, S., Barkai, E. & Maroun, M. Olfactory learning-induced enhancement of the predisposition for LTP induction. Learn. Mem. 18, 594–597 (2011).

54. Kundu, S., Paul, B., Reuevni, I., Lamprecht, R. & Barkai, E. Learning-induced bidirectional enhancement of inhibitory synaptic metaplasticity. The Journal of Physiology 602, 2343– 2358 (2024).

55. Harvey, C. D. & Svoboda, K. Locally dynamic synaptic learning rules in pyramidal neuron dendrites. Nature 450, 1195–1200 (2007).

56. Lillicrap, T. P., Santoro, A., Marris, L., Akerman, C. J. & Hinton, G. Backpropagation and the brain. Nat Rev Neurosci 21, 335–346 (2020).

57. Murray, J. M. Local online learning in recurrent networks with random feedback. eLife 8, e43299 (2019).

58. Bellec, G. et al. A solution to the learning dilemma for recurrent networks of spiking neurons. Nat Commun 11, 3625 (2020).

59. Frey, U. & Morris, R. G. M. Synaptic tagging and long-term potentiation. Nature 385, 533– 536 (1997).

60. Foster, D. J. & Wilson, M. A. Reverse replay of behavioural sequences in hippocampal place cells during the awake state. Nature 440, 680–683 (2006).

61. Ambrose, R. E., Pfeiffer, B. E. & Foster, D. J. Reverse Replay of Hippocampal Place Cells Is Uniquely Modulated by Changing Reward. Neuron 91, 1124–1136 (2016).

62. Blanco-Pozo, M., Akam, T. & Walton, M. E. Dopamine-independent effect of rewards on choices through hidden-state inference. Nat Neurosci 27, 286–297 (2024).

63. Parker, N. F. et al. Reward and choice encoding in terminals of midbrain dopamine neurons depends on striatal target. Nat Neurosci 19, 845–854 (2016).

64. Pan-Vazquez, A. et al. VTA dopamine neuron activity produces spatially organized stimulus and action value representations through conditioned reinforcement. Preprint at 10.1101/2025.11.04.685995 (2025).

65. Yu, K. et al. The central amygdala controls learning in the lateral amygdala. Nat Neurosci 20, 1680–1685 (2017).

66. Yang, T. et al. Plastic and stimulus-specific coding of salient events in the central amygdala. Nature 616, 510–519 (2023).

67. Saez, A., Rigotti, M., Ostojic, S., Fusi, S. & Salzman, C. D. Abstract Context Representations in Primate Amygdala and Prefrontal Cortex. Neuron 87, 869–881 (2015).

68. Mongillo, G., Barak, O. & Tsodyks, M. Synaptic Theory of Working Memory. Science 319, 1543–1546 (2008).

69. Masse, N. Y., Yang, G. R., Song, H. F., Wang, X.-J. & Freedman, D. J. Circuit mechanisms for the maintenance and manipulation of information in working memory. Nature Neuroscience 22, 1159–1167 (2019).

70. Stokes, M. G. ‘Activity-silent’ working memory in prefrontal cortex: a dynamic coding framework. Trends in Cognitive Sciences 19, 394–405 (2015).

71. Panichello, M. F. et al. Intermittent rate coding and cue-specific ensembles support working memory. Nature 636, 422–429 (2024).

72. Zucker, R. S. & Regehr, W. G. Short-Term Synaptic Plasticity. Annual Review of Physiology 64, 355–405 (2002).

73. Fioravante, D. & Regehr, W. G. Short-term forms of presynaptic plasticity. Current Opinion in Neurobiology 21, 269–274 (2011).

74. Chan, S. C. Y. et al. Data distributional properties drive emergent in-context learning in transformers. in Advances in Neural Information Processing Systems vol. 35 18878–18891 (Curran Associates, 2022).

75. Pesnot Lerousseau, J. & Summerfield, C. Shared sensitivity to data distribution during learning in humans and transformer networks. Nature Human Behaviour 10, 601–614 (2025).

76. Russin, J., Pavlick, E. & Frank, M. J. Parallel trade-offs in human cognition and neural networks: The dynamic interplay between in-context and in-weight learning. Proceedings of the National Academy of Sciences 122, e2510270122 (2025).

77. Rmus, M., McDougle, S. D. & Collins, A. G. The role of executive function in shaping reinforcement learning. Current Opinion in Behavioral Sciences 38, 66–73 (2021).

78. Frank, M. J. & Claus, E. D. Anatomy of a decision: Striato-orbitofrontal interactions in reinforcement learning, decision making, and reversal. Psychological Review 113, 300–326 (2006).

79. Razmi, N. & Nassar, M. R. Adaptive Learning through Temporal Dynamics of State Representation. The Journal of Neuroscience 42, 2524–2538 (2022).

80. Whittington, J. C. R., Dorrell, W., Behrens, T. E. J., Ganguli, S. & El-Gaby, M. A tale of two algorithms: Structured slots explain prefrontal sequence memory and are unified with hippocampal cognitive maps. Neuron 113, 321–333.e6 (2025).

81. Finn, C., Abbeel, P. & Levine, S. Model-Agnostic Meta-Learning for Fast Adaptation of Deep Networks. in Proceedings of the 34th International Conference on Machine Learning 1126–1135 (PMLR, 2017).

82. Brown, T. B. et al. Language models are few-shot learners. in Advances in Neural Information Processing Systems vol. 33 1877–1901 (Curran Associates, 2020).

83. Lee, J. & Sabatini, B. L. Striatal indirect pathway mediates exploration via collicular competition. Nature 1–5 (2021) doi:10.1038/s41586-021-04055-4.

84. Sumi, M. et al. The newly synthesized selective Ca2+calmodulin dependent protein kinase II inhibitor KN-93 reduces dopamine contents in PC12h cells. Biochemical and Biophysical Research Communications 181, 968–975 (1991).

85. Kobak, D. et al. Demixed principal component analysis of neural population data. eLife 5, e10989 (2016).

86. Werbos, P. J. Backpropagation through time: what it does and how to do it. Proceedings of the IEEE 78, 1550–1560 (1990).

87. Williams, R. J. & Zipser, D. Gradient-based learning algorithms for recurrent networks and their computational complexity. in Backpropagation: theory, architectures, and applications 433–486 (L. Erlbaum Associates Inc., USA, 1995).

88. Sussillo, D. & Barak, O. Opening the Black Box: Low-Dimensional Dynamics in High-Dimensional Recurrent Neural Networks. Neural Computation 25, 626–649 (2013).

